# The Arabidopsis histone H3K4me3-binding ALFIN-like proteins mediate histone H2A ubiquitination and coordinate diverse chromatin modifications

**DOI:** 10.1101/2024.09.12.612777

**Authors:** Xiao-Min Su, Dan-Yang Yuan, Lin Li, Minqi Yang, She Chen, Yue Zhou, Xin-Jian He

**Author notes:** Corresponding author: Xin-Jian He.

## Abstract

The histone H3K4 trimethylation (H3K4me3) is widely distributed at numerous actively transcribed protein-coding genes throughout the genome. However, the interplay between H3K4me3 and other chromatin modifications remains poorly understood in plants. In this study, we find that the *Arabidopsis thaliana* H3K4me3-binding ALFIN-LIKE (AL) proteins are associated with H3K4me3-enriched genes at the whole-genome level. The AL proteins contain a C-terminal PHD finger, which has a conserved role in recognizing H3K4me3, and a PHD-associated AL (PAL) domain, which is responsible for binding to diverse chromatin-related proteins. We demonstrate that the AL proteins not only act as subunits of the Polycomb repressive complex 1 (PRC1) to mediate H2A ubiquitination at H3K4me3-enriched genes but also interact with a variety of other chromatin-related proteins. Furthermore, we elucidate the mechanisms by which AL proteins interact with other chromatin-associated proteins to integrate H3K4me3, H2A ubiquitination, H2A.Z deposition, H3K27 demethylation, and chromatin accessibility across the genome. These findings underscore the critical role of AL proteins in linking H3K4me3 with a variety of other chromatin modifications in plants.

## Introduction

In eukaryotic cells, the nucleosome serves as the fundamental unit of chromatin, consisting of 147 base pairs of DNA wrapped around a histone octamer, which is composed of two molecules each of H2A, H2B, H3, and H4 (Peterson and Laniel, 2004). These nucleosomes are compacted into higher-order chromatin structures, which are essential for condensing genomic DNA into the confined space of the nucleus (Luger et al., 1997; Jansen and Verstrepen, 2011). However, this compaction also restricts DNA accessibility and influences various processes associated with chromatin, including gene transcription, DNA replication, and DNA damage repair (Li and Reinberg, 2011). Therefore, the dynamic regulation of chromatin is crucial to balance its compaction with the necessity for accessibility. Eukaryotes have evolved several mechanisms for this dynamic regulation, including histone modifications, deposition of histone variants, and chromatin remodeling (Clapier and Cairns, 2009; Bannister and Kouzarides, 2011; Weber and Henikoff, 2014).

Histone modifications encompass a variety of post-translational changes such as acetylation, methylation, ubiquitination, phosphorylation, and SUMOylation (Portela and Esteller, 2010; Bannister and Kouzarides, 2011). These modifications can impact the chromatin state either by directly altering the interaction between DNA and histones or indirectly through the action of "reader" proteins that recognize these modifications (Liu et al., 2010). Histone methylation occurs at lysine and arginine residues of histone tails (Lachner and Jenuwein, 2002; Di Lorenzo and Bedford, 2011). Methylation of different lysine sites has distinct effects on transcriptional regulation, with certain marks like H3K4me3 and H3K36me3 promoting transcriptional activation, while others such as H3K9me2 and H3K27me3 are associated with repression (Dambacher et al., 2010; Suganuma and Workman, 2011). The influence of histone methylation on chromatin is primarily mediated by "reader" proteins, with domains like PHD, Chromo, BAH, and Tudor identified as “readers” of histone methylation (Bannister and Kouzarides, 2011; Liu et al., 2018).

ALFIN-LIKE (AL) proteins are identified across a broad spectrum of plant species, ranging from unicellular algae to angiosperms, yet they are absent in metazoans and fungi (Lee et al., 2009; Kayum et al., 2015; Zhou et al., 2016; Wang et al., 2023). In Arabidopsis, seven AL proteins (AL1-AL7) share a common structural feature: a conserved C-terminal PHD finger and an N-terminal PHD-associated AL (PAL) domain, with the PHD finger capable of binding the H3K4me3 mark (Lee et al., 2009; Liang et al., 2018). These AL proteins have been implicated in various biological processes, including ABA signaling, seed germination, and responses to biotic and abiotic stresses (Wei et al., 2015; Yan et al., 2022; Wang et al., 2023; Zhang et al., 2023; Jin et al., 2024). However, due to limitations in generating higher-order mutants, previous studies have predominantly focused on individual or a few AL proteins, necessitating a comprehensive investigation into the collective function of all seven Arabidopsis AL proteins.

Four Arabidopsis AL proteins (AL4, AL5, AL6, and AL7) have been reported to be co-purified with the SWI2/SNF2-Related 1 (SWR1) complex component ACTIN-RELATED PROTEIN 6 (ARP6) (Potok et al., 2019). Given the role of the SWR1 complex in the deposition of the histone variant H2A.Z into chromatin (March-Dīaz and Reyes, 2009; Gómez-Zambrano et al., 2018; Shang and He, 2022), this co-purification suggests a potential link between H2A.Z deposition and the H3K4me3 mark. Additionally, AL6 and AL7 have been shown to interact with components of the Polycomb Repressive Complex 1 (PRC1), RING1A and BMI1B, through their PAL domain, facilitating the transition from H3K4me3 to H3K27me3 at seed development-related genes during germination (Molitor et al., 2014; Peng et al., 2018). In Arabidopsis, the PRC1 complex, which is responsible for H2A ubiquitination (H2Aub), includes two RING1 proteins (RING1A/B) and three BMI1 proteins (BMI1A/B/C) (Bratzel et al., 2010; Chen et al., 2010; Grossniklaus and Paro, 2014; Li et al., 2017). Although earlier studies have indicated that AL proteins interact with components of SWR1 and PRC1 complexes, it is yet to be determined whether AL proteins affect SWR1-mediated H2A.Z deposition or PRC1-mediated H2Aub and whether AL proteins interact any other chromatin-related proteins in addition to SWR1 and PRC1 components.

This study presents an extensive examination of all seven AL proteins in Arabidopsis. Utilizing T-DNA mutants, CRISPR/Cas9-induced mutagenesis, and genetic crossing, we generated single and multi-mutants for each AL protein. Despite our success in creating a range of multi-mutants, a septuple mutant disrupting all seven AL proteins could not be obtained, highlighting the essential and redundant roles of AL proteins for plant viability. Several viable high-order mutants exhibited severe defects in both vegetative and reproductive development. By performing chromatin immunoprecipitation followed by sequencing, we found that the Arabidopsis AL proteins associate with genomic regions near the transcription start site (TSS) of genes enriched with H3K4me3. Both the H3K4me3-binding PHD finger and the PAL domain contribute to the association of AL proteins with chromatin. Furthermore, we found that AL proteins serve as PRC1 subunits and are crucial for PRC1-mediated H2Aub, revealing a plant-specific link between H3K4me3 and H2Aub. Moreover, we found that AL proteins interact with chromatin-related components involved in regulating multiple histone modifications, H2A.Z deposition, and chromatin remodeling, shedding light on the role of AL proteins in orchestrating multiple chromatin modifications.

## Results

### Arabidopsis AL proteins are essential for plant growth and development

The Arabidopsis ALFIN-LIKE (AL) proteins have been demonstrated to influence seed germination and stress tolerance, with minimal impact on post-germination plant development (Molitor et al., 2014; Wei et al., 2015). However, previous studies have primarily focused on one or two family members, leaving the essentiality of AL proteins in Arabidopsis largely unexplored. In this study, we obtained each *al* single mutant from the Arabidopsis Biological Resource Center or generated them through CRISPR/Cas9-induced mutagenesis (Supplemental Figure 1). Subsequently, we investigated a range of phenotypic traits in these mutants, including days to bolting, the number of rosette leaves, rosette diameter, fresh weight, and silique length. We found that the *al2* and *al4* single mutants exhibited a mild late-flowering phenotype, characterized by a delayed bolting and an increased number of rosette leaves, they did not display any additional developmental abnormalities (Supplemental Figure 2). In contrast, no flowering-time delays or developmental phenotypes were observed in the other *al* single mutants (Supplemental Figure 2).

We utilized CRISPR/Cas9-induced mutagenesis and genetic crossing to generate various combinations of *al* mutants. As a result, we obtained a range of double (*al1/3*, *al2/3*, *al3/4*, *al2/4*, *al2/7* and *al6/7*), triple (*al2/3/4*, *al2/4/5*, *al2/4/7*, *al2/6/7*, *al4/5/6* and *al4/5/7*), quadruple (*al1/2/3/4*, *al2/3/4/5*, *al2/4/5/6*, *al2/4/5/7*, *al2/4/6/7* and *al4/5/6/7*), and quintuple mutants (*al1/2/3/4/5*, *al3/4/5/6/7*, *al2/4/5/6/7*, *al1/3/4/5/7* and *al2/3/4/5/7*) for all seven Arabidopsis AL proteins. We observed that the *al2/3*, *al2/4* and *al2/7* double mutants, but not the other double mutants, displayed varying degrees of late flowering, while the other double mutants tested did not show noticeable abnormalities compared to the wild type (Supplemental Figure 3). The late-flowering phenotype in the *al2/4* double mutant were further validated through complementation testing (Supplemental Figure 4), confirming the role of AL proteins in regulating flowering time.

Except for the *al4/5/6* and *al4/5/7* triple mutants, all the other tested triple mutants showed late flowering (Supplemental Figure 5). The late-flowering phenotype was more pronounced in the *al2/4/7* mutant than in the other triple mutants (Supplemental Figure 5). Additionally, the *al2/4/7* mutant, but not the other triple mutants, also showed developmental defects, including smaller plant stature and shorter siliques (Supplemental Figure 5). In comparison to the developmental defects in the *al2/4/7* triple mutant, the defects were exacerbated in several quadruple mutants and to a greater extent in quintuple *al* mutants, as indicated by smaller plant stature, aberrant floral organs, short siliques, and poor fertility (Figure 1A-C, Supplemental Figure 6-7). This suggests that different AL proteins exhibit redundancy in regulating plant development. Differing from the developmental defects that are consistently exacerbated with increasing *al* mutations, the late-flowering phenotype was enhanced in quadruple mutants compared to triple mutants, but was not further enhanced or impaired in quintuple mutants (Supplemental Figure 5-7), which suggests that the late-flowering phenotype is influenced by the severe developmental defects in quintuple mutants.

**Figure 1.**
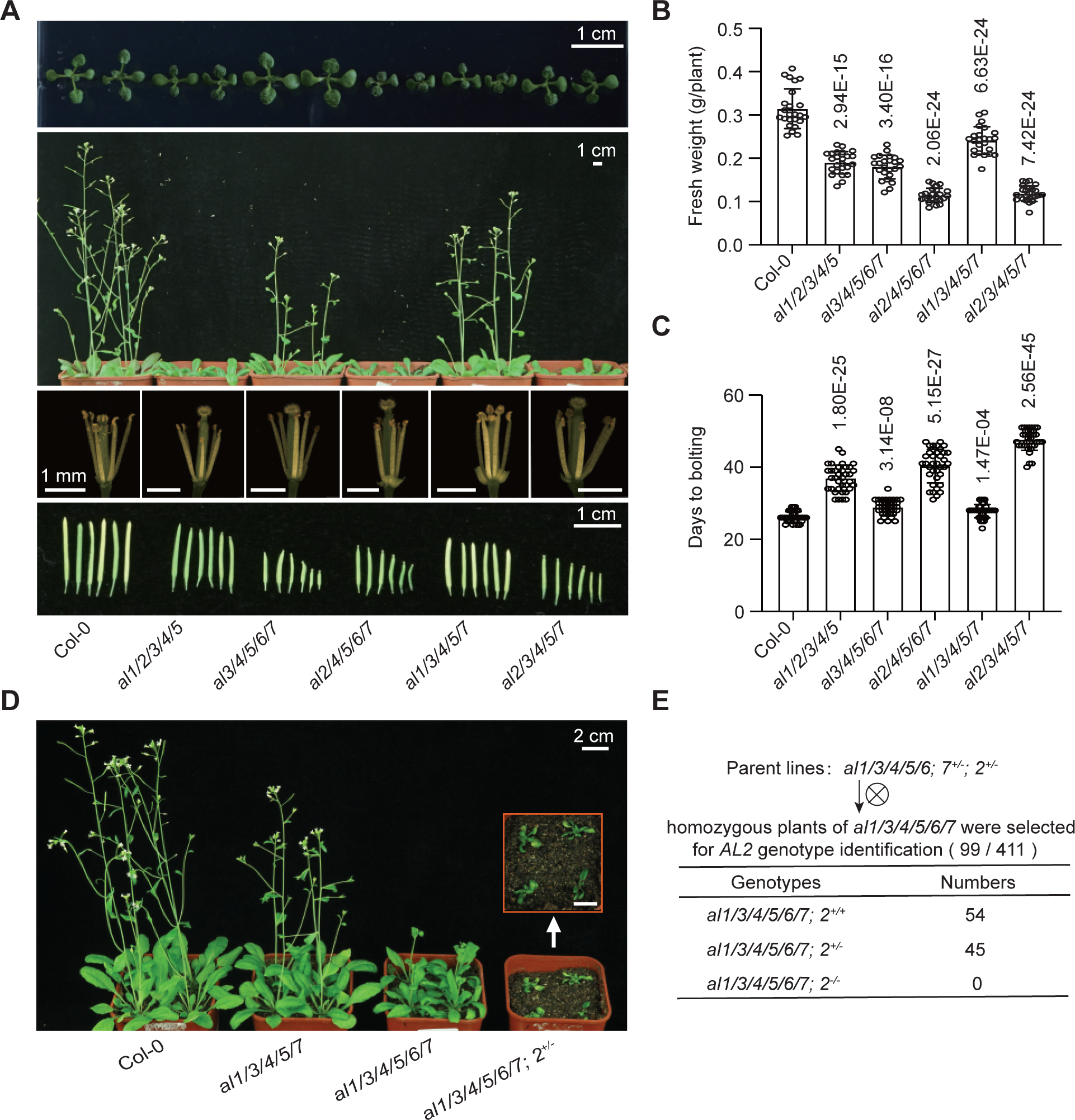
Arabidopsis AL proteins are essential for plant growth and development. **(A)** Morphological phenotypes of the *al* quintuple mutants on plant growth and development are shown. From top to bottom, the phenotype of 11-day-old plants (top), 33-day-old plants, stamens and pistils under a microscope, and mature siliques (bottom) are presented for both wild-type and *al* quintuple mutants. Scale bars, from top to bottom, are 1 cm, 1 cm, 1 mm, and 1 cm. **(B)** The statistical results of fresh weight per plant (25-day-old plants, *n* = 24). Values are mean ± SD. *P* values determined by two-tailed Student’s t-test indicate the difference between the indicated mutants and the wild-type Col-0 control. **(C)** The statistical results of days to bolting (*n* = 36). Values are mean ± SD. *P* values determined by two-tailed Student’s t-test indicate the difference between the indicated mutants and the wild-type Col-0 control. **(D)** Phenotypes of *al* multiple mutants with indicated genotypes. **(E)** Statistical analyses of the progeny of self-bred *al1/3/4/5/6; 7^+/-^; 2^+/-^* plants. Homozygous plants of *al1/3/4/5/6/7* were selected for *AL2* genotype identification. The observed numbers with indicated genotypes are shown.

Through the combination of CRISPR/Cas9-induced mutagenesis and genetic crossing, we obtained the “*al1/3/4/5/6* homozygous while *al2* and *al7* heterozygous” (*al1/3/4/5/6;2^+/-^;7^+/-^*) mutant. In the self-fed progeny of the *al1/3/4/5/6;2^+/-^;7^+/-^*mutant, we obtained numerous *al1/3/4/5/6/7* sextuple mutant plants and the “*al1/3/4/5/6/7* homozygous while *al2* heterozygous” (*al1/3/4/5/6/7;2^+/-^*) mutant plants but failed to obtain the *al1/2/3/4/5/6/7* septuple mutant plants (Figure 1D and E). While the *al1/3/4/5/6/7* mutant plants showed reduced stature and diminished fertility, the *al1/3/4/5/6/7;2^+/-^* mutant plants exhibited more pronounced developmental defects, with a failure to bolt (Figure 1D and E). These results suggest that the redundant AL proteins play an essential role in the regulation of plant growth and development.

To elucidate the potential functional specificities of AL proteins, we conducted an integrated analysis of various phenotypic traits, including days to bolting, the number of rosette leaves, rosette diameter, fresh weight, and silique length, across all the *al* mutants tested in this study. By comparing the phenotype changes between the mutants and the wild type, we obtained a contribution coefficient for each *al* mutation, this coefficient was derived from the statistical significance of the observed changes, as represented by the -log_10_ (*P* value) (Supplemental Figure 8). Our analysis revealed that while AL proteins exhibit redundancy in the regulation of both flowering time and plant development, certain AL proteins also show distinct functional specificities, with AL2 primarily influencing flowering time and AL3 mainly contributing to plant development.

### AL proteins interact with multiple chromatin-related proteins in Arabidopsis plants

To identify AL-interacting proteins in Arabidopsis plants, we conducted affinity purification followed by mass spectrometry (AP–MS) in transgenic plants individually expressing seven FLAG-tagged AL proteins (AL1, AL2, AL3, AL4, AL5, AL6, and AL7). We found that all the AL proteins were co-purified with each of the FLAG-tagged AL proteins (Figure 2A-B; Supplemental Dataset 1), indicating that the AL proteins interact with each other in Arabidopsis plants.

**Figure 2.**
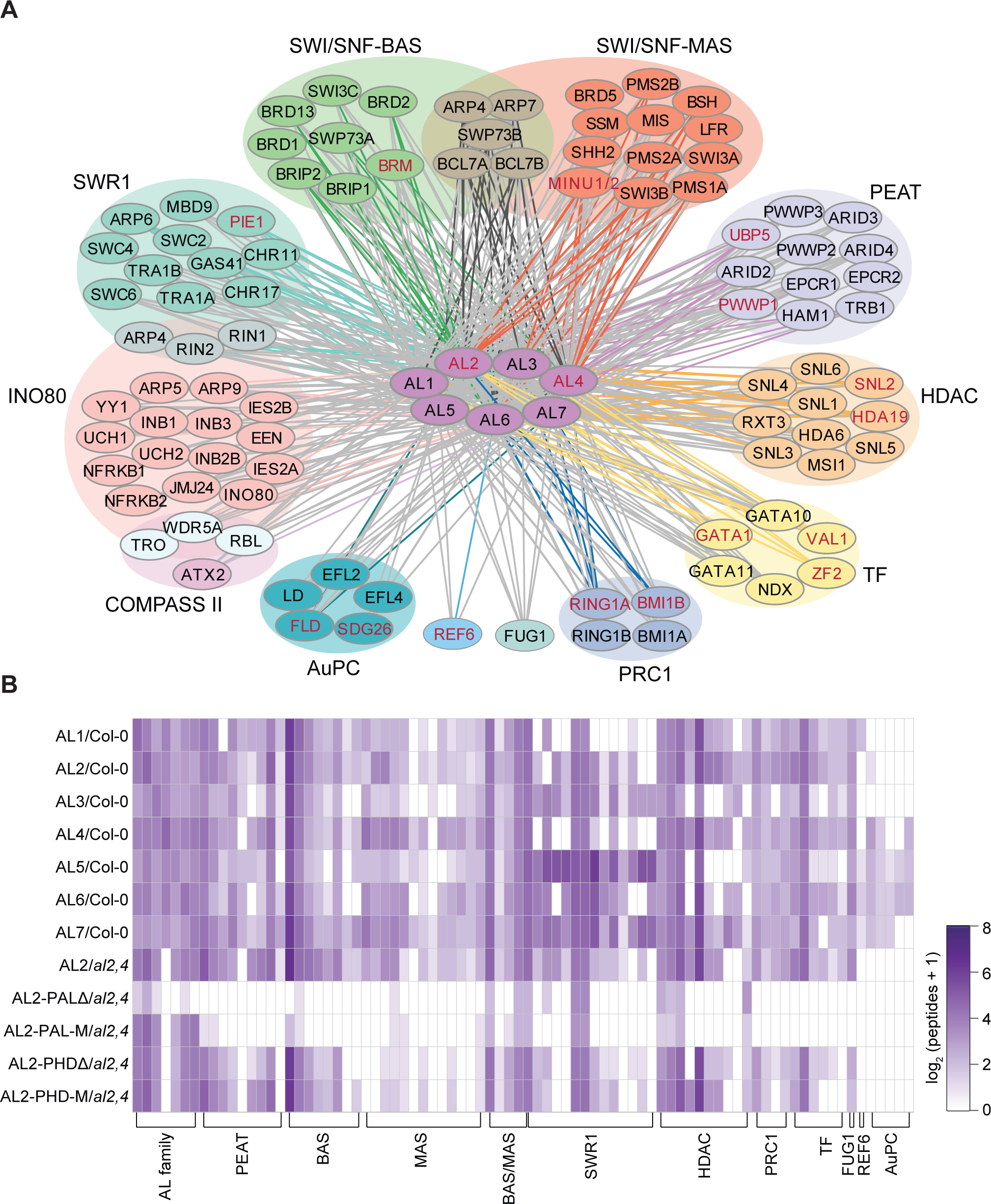
AL proteins interact with multiple chromatin-related proteins in Arabidopsis plants. **(A)** The diagram indicates the interactions between the AL proteins and various chromatin-related proteins as determined by AP-MS. The diagram was drawn with Cytoscape (v3.6.1). In the diagram, each unique color represents a distinct class of chromatin-related proteins. The gray lines denote chromatin-related proteins that are co-purified with the AL proteins in the AP-MS assays, while the colored lines indicate that the interactions between the AL proteins and chromatin-related proteins are validated by bidirectional AP-MS assays. The chromatin-related proteins highlighted in red have been demonstrated to interact with the AL2 or AL4 through pull-down assays conducted in this study. **(B)** The heatmap displays the relative abundance of chromatin-related proteins co-purified with AL proteins. The color intensity represents the enrichment of normalized peptides identified by AP-MS. Transgenic plants expressing either wild-type or mutated AL proteins were subjected to AP-MS.

Furthermore, we discovered that numerous chromatin-related proteins were co-purified with AL proteins, including the SWI/SNF complexes, which facilitate chromatin accessibility (Guo et al., 2022; Fu et al., 2023), the PEAT complex involved in H2A deubiquitination and H4K5 acetylation (Zheng et al., 2023; Godwin et al., 2024), the SWR1 complex involved in the histone variant H2A.Z deposition (Gómez-Zambrano et al., 2018; Potok et al., 2019; Sijacic et al., 2019; Luo et al., 2020), the INO80 complex involved in genome stability maintenance and exchanging between H2A and H2A.Z (Zhang et al., 2015; Yang et al., 2020; Shang et al., 2021; Xue et al., 2021), the HDAC complex responsible for histone deacetylation (Perrella et al., 2013; Mehdi et al., 2016; Ning et al., 2019; Feng et al., 2021), the AuPC complex linked to the autonomous flowering regulation pathway (Qi et al., 2022), the PRC1 complex responsible for H2A monoubiquitination (Bratzel et al., 2010; Yang et al., 2013; Barbour et al., 2020), and the histone H3K27me3 demethylase RELATIVE OF EARLY FLOWERING 6 (REF6) (Lu et al., 2011) (Figure 2A, Supplemental Dataset 1).

Reciprocally, based on our previous AP-MS results (Ning et al., 2019; Shang et al., 2021; Guo et al., 2022; Qi et al., 2022; Zheng et al., 2023), AL proteins were also co-purified with chromatin-related proteins (Supplemental Figure 9), although the identification of AL proteins were not noted in those studies, thereby confirming the interaction of AL proteins with these chromatin-related proteins. Earlier studies from other groups also indicated that AL proteins interact with the SUMO protease FUG1 (Sureshkumar et al., 2024), the SWR1 complex component ARP6 (Potok et al., 2019), and the PRC1 complex components RING1A and BMI1B (Molitor et al., 2014; Peng et al., 2018). Our AP-MS analysis, using transgenic plants that expressed FLAG-tagged AL, RING1A, and BMI1B proteins, confirmed the interaction between AL proteins and the PRC1 complex components RING1A, RING1B, BMI1A, and BMI1B (Supplemental Figure 10, Supplemental Dataset 1). All these proteins were also identified as AL-interacting proteins in our AP-MS analysis (Figure 2A, Supplemental Figure 9-10, Supplemental Dataset 1), affirming the interaction of AL proteins with these components.

Moreover, our AP-MS data revealed that several transcription factors were co-purified with AL proteins. These include the previously characterized transcriptional repressors VAL1 and NDX (Karanyi et al., 2022; Liang et al., 2022), as well as the previously uncharacterized transcription factors GATA1, GATA10, GATA11, and ZF2 (Figure 2A, Supplemental Figure 10, Supplemental Dataset 1). In addition, we identified several putative transcription factors that have not yet been formally named (Supplemental Figure 10). To validate the interaction between AL proteins and the transcription factors, we generated transgenic plants that individually expressed *GATA1-FLAG, VAL1-FLAG* and *ZF2-FLAG* transgenes, and then performed AP-MS analysis in the transgenic plants. The results indicated that AL proteins were co-purified with GATA1, VAL1 and ZF2 in the corresponding transgenic plants (Supplemental Figure 10, Supplemental Dataset 1), confirming that the AL proteins interact with the transcription factors in Arabidopsis plants. Furthermore, we conducted pull-down assays to investigate whether the AL proteins directly interact with the transcription factors. We found that the AL proteins directly interact with GATA1, ZF2, and VAL1, but not with NDX (Supplemental Figure 11).

### The PAL domain of AL proteins is responsible for interacting with chromatin-related proteins

Phylogenetic analysis indicated that seven Arabidopsis AL proteins are highly similar and can be divided into three subclasses (subclass I: AL1 and AL2; subclass II: AL3, AL4, and AL5; subclass III: AL6 and AL7) (Supplemental Figure 12). All the AL proteins contain a C-terminal PHD finger and an N-terminal PHD-associated AL (PAL) domain (Figure 3A; Supplemental Figure 12). The PHD and PAL domains were found in the AL proteins from different plant species, including algae, bryophytes, ferns, gymnosperms, and angiosperms (Supplemental Figure 13), which is consistent with the previous reports (Kayum et al., 2015; Jin et al., 2024).

**Figure 3.**
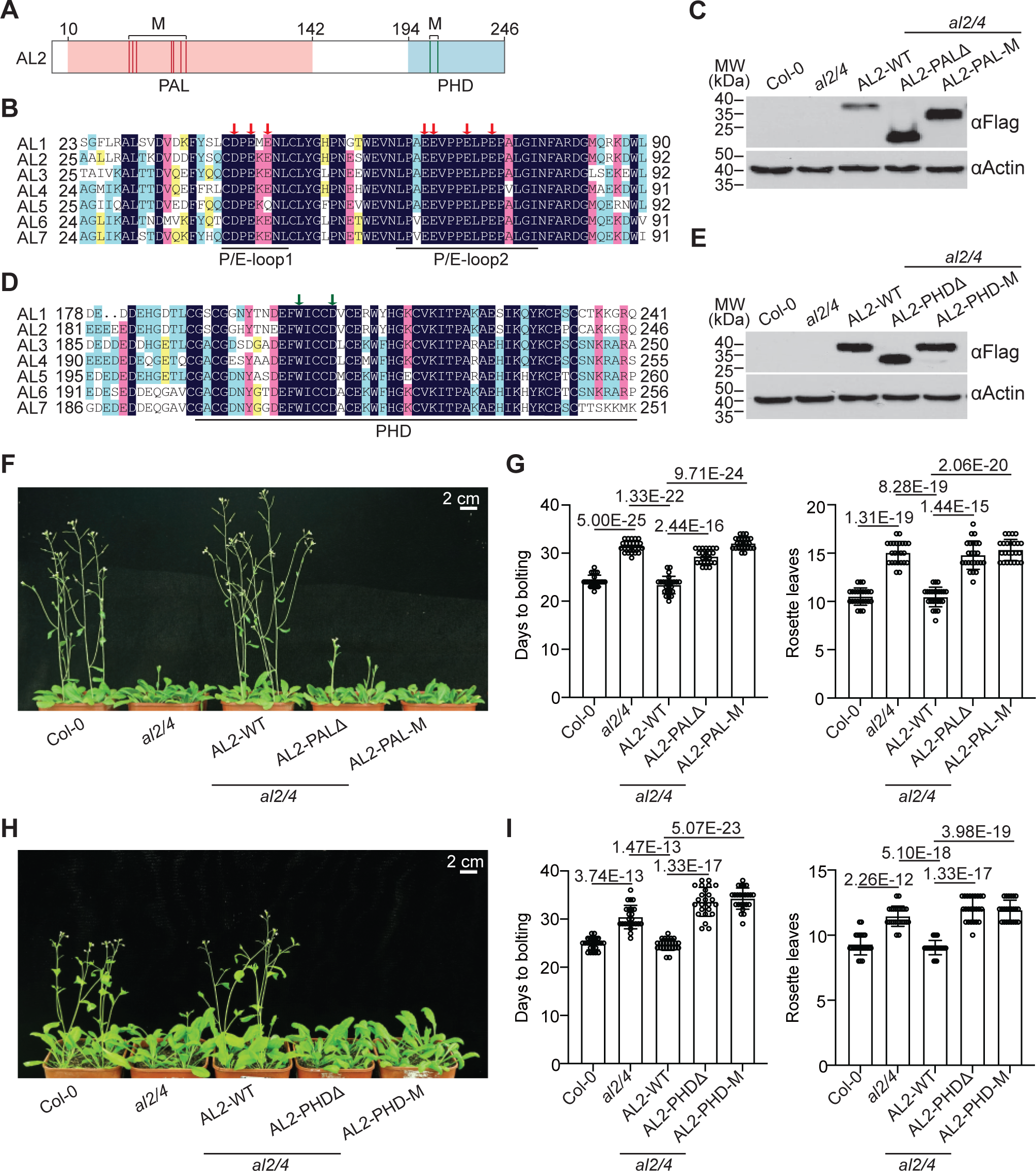
Essential roles of the PHD and PAL domains in Arabidopsis AL proteins. **(A)** Schematic representation of the PAL and PHD domains of AL2. Seven conserved acidic residues within the PAL domain (D43, E45, E47, E66, E67, E71, and E74) are denoted by red lines, while the two conserved residues in the PHD finger (W206 and D210) are indicated with green lines. **(B)** Sequence alignment of the PAL domain in Arabidopsis AL proteins. Seven conserved acidic residues, marked with red arrows, were all mutated to alanine (A) in the *AL2-PAL-M* construct. **(C)** The expression levels of *AL2-WT*, *AL2-PAL*Δ, and *AL2-PAL-M* in transgenic plants were determined by western blotting. The Actin signal is indicated as a loading control. **(D)** Sequence alignment of the PHD finger in Arabidopsis AL proteins. The conserved residues W206 and D210, marked with green arrows, were mutated to alanine (A) and asparagine (N), respectively, in the *AL2-PHD-M* construct. **(E)** The expression levels of *AL2-WT*, *AL2-PHD*Δ, and *AL2-PHD-M* in transgenic plants were determined by western blotting, with Actin as a loading control. **(F)** Determination of the restoration of the late-flowering phenotype of the *al2/4* mutant by *AL2-WT*, *AL2-PAL*Δ, and *AL2-PAL-M* transgenes. The morphological phenotypes of the plants with indicated genotypes are shown. **(G)** Statistical analysis of the flowering time in *AL2-WT*, *AL2-PAL*Δ, and *AL2-PAL-M* complementation lines. The number of days to bolting (left) and the number of rosette leaves (right) are shown. The sample size is 24, with values shown as mean ± SD. *P* values were determined by two-tailed Student’s t-test. **(H)** Determination of the restoration of the late-flowering phenotype of the *al2/4* mutant by *AL2-WT*, *AL2-PHD*Δ, and *AL2-PHD-M* transgenes. The morphological phenotypes of plants with indicated genotypes are shown. **(I)** Statistical analysis of the flowering time in *AL2-WT*, *AL2-PHD*Δ, and *AL2-PHD-M* complementation lines. The number of days to bolting (left) and the number of rosette leaves (right) are shown. The sample size is 24, with values presented as mean ± SD. *P* values were determined by two-tailed Student’s t-test.

To investigate whether the PHD finger of AL proteins is involved in the interaction of AL proteins with chromatin-related proteins identified in our AP-MS analysis, we generated transgenic plants expressing FLAG-tagged PHD-truncated (AL2-PHDΔ) and PHD-mutated (W206 and D210 mutated to alanine, AL2-PHD-M) versions of AL2 (Figure 3A, 3D). By performing AP-MS using the transgenic plants, we found that most of the chromatin-related proteins co-purified with the wild-type AL2 were also co-purified with AL2-PHDΔ and AL2-PHD-M, suggesting that the PHD finger is unlikely to play a major role in the interaction of AL2 with chromatin-related proteins (Figure 2B, Supplemental Dataset 1).

The PAL domain of AL proteins have been shown interact with RING1A and BMI1B, two conserved components of PRC1 complex (Molitor et al., 2014). Structural analysis has revealed that the PAL domain of AL2 exists as a dimer and forms an acidic pocket between two proline- and glutamic acid-rich loops (P/E-loop1 and 2), which facilitates the interaction of ALs with RING1A and BMI1B (Peng et al., 2018). To investigate whether the acidic pocket in the PAL domain is necessary for the interaction of AL proteins with chromatin-related proteins, we conducted AP-MS using transgenic plants expressing PAL-depleted (AL2-PALΔ) and PAL-mutated (seven acidic residues mutated to alanine, AL2-PAL-M) versions of AL2 (Figure 3A, 3B). We found that most of the chromatin-related proteins that were co-purified with the wild-type AL2 were not co-purified with AL2-PALΔ or AL2-PAL-M (Figure 2B, Supplemental Dataset 1). These results indicate that the acidic pocket in the PAL domain is essential for AL proteins to bind to multiple chromatin-related proteins in Arabidopsis plants.

We performed *in vitro* pull-down assays to investigate the protein-protein interactions identified by AP-MS analysis. We observed that that the AL proteins can interact with each other through both the PHD and PAL domains (Supplemental Figure 14). Furthermore, our pull-down assay demonstrated that that AL2 directly interacts with multiple chromatin-related proteins, including the BAS-type SWI/SNF chromatin remodeler BRM (Guo et al., 2022; Fu et al., 2023) (Supplemental Figure 11), the MAS-type SWI/SNF chromatin remodelers MINU1 and MINU2 (Diego-Martin et al., 2022; Guo et al., 2022; Fu et al., 2023) (Supplemental Figure 11), the SWR1 catalytic subunit PIE1 (Noh and Amasino, 2003; Rosana March-Díaz and Reyes, 2007), the HDAC catalytic subunit HDA19 (Ning et al., 2019) (Supplemental Figure 15), the HDAC accessory subunit SNL2 (Wang et al., 2013) (Supplemental Figure 15), the LSD-like histone demethylase FLD (He et al., 2003; Spedaletti et al., 2008; Qi et al., 2022) (Supplemental Figure 15), the histone methyltransferase SDG26 (also known as a subunit of the AuPC complex) (Berr et al., 2015; Qi et al., 2022) (Supplemental Figure 15), the PEAT complex subunit PWWP1 (Hohenstatt et al., 2018) (Supplemental Figure 16), the H2A deubiquitinase UBP5 (also known as a subunit of the PEAT complex) (Zheng et al., 2023; Godwin et al., 2024) (Supplemental Figure 16), the histone H3K27 demethylase REF6 (Lu et al., 2011) (Supplemental Figure 16), the PRC1 complex subunits RING1A and BMI1B (Chen et al., 2010; Molitor and Shen, 2013) (Supplemental Figure 16), as well as the transcription factors GATA1, VAL1, and ZF2 (Chiang et al., 2012; Liang et al., 2022) (Figure 2A; Supplemental Figure 11).

Nevertheless, our pull-down assay failed to detect a direct interaction of AL2 with the ISWI chromatin remodeler CHR11 and the accessory subunit of the SWI/SNF complex SWI3B (Supplemental Figure 11, 15). Although our AP-MS analysis has revealed that AL2 interacts with both CHR11 and SWI3B in Arabidopsis plants (Figure 2A, Supplemental Dataset 1), the interactions are likely mediated by other proteins. Given that CHR11 is a constituent of the SWR1 complex and SWI3B is part of the MAS-type SWI/SNF complex (Luo et al., 2020; Guo et al., 2022), the direct interactions of AL2 with the SWR1 catalytic subunit PIE1 and the MAS catalytic subunit MINU2, as evidenced by our pull-down assay (Figure 7A, Supplemental Figure 11), likely facilitate the association of AL2 with CHR11 and SWI3B, respectively, in Arabidopsis plants.

### Both the PHD and PAL domains are essential for the function of AL proteins in Arabidopsis plants

The PHD finger of AL proteins has been previously reported to possess the binding ability for histone H3K4me3 and to a lesser extent for H3K4me2 (Lee et al., 2009). Our *in vitro* pull-down assay confirmed the binding ability of the PHD finger of AL2 for H3K4me3 and H3K4me2, and demonstrated that the W206A and D210A mutations in the PHD finger (AL2-PHD-M) completely disrupted the binding (Supplemental Figure 17). To investigate the necessity of the PHD finger for the function of AL proteins in Arabidopsis plants, we introduced the PHD-depleted *AL2-PHD*Δ, the PHD-mutated *AL2-PHD-M*, and the wild-type *AL2* (*AL2-WT*) into the *al2/4* mutant for complementation testing (Figure 3E). Our findings showed that the wild-type *AL2* transgene, but not the *AL2-PHD*Δ or *AL2-PHD-M* transgenes, complemented the late-flowering phenotype of the *al2/4* double mutant (Figure 3H, 3I). These results suggest that the conserved H3K4me3-binding PHD finger is essential for the function of AL proteins in Arabidopsis plants.

Given that the PAL-depleted AL2-PALΔ and the acidic residue-mutated AL2-PAL-M disrupted the interaction of AL2 with multiple chromatin-related proteins in Arabidopsis plants (Figure 2B; Supplemental Dataset 1), we investigated whether these mutations affect the biological function of AL2 by transforming the *AL2-PAL*Δ, *AL2-PAL-M*, and *AL2-WT* constructs into the *al2/4* mutant for complementation testing (Figure 3C). Our results demonstrated that the late-flowering phenotype of the *al2/4* mutant was restored by the *AL2-WT* transgene but not by the *AL2-PAL*Δ or *AL2-PAL-M* transgenes (Figure 3F, 3G). This indicates that the conserved acidic residues within the PAL domain are indispensable for the function of AL proteins in Arabidopsis plants.

### Different AL proteins co-occupy chromatin at the whole-genome level

To explore the genome-wide distribution of AL proteins, we conducted chromatin immunoprecipitation followed by sequencing (ChIP-seq) for the Arabidopsis AL proteins, including AL1, AL2, AL3, AL4, and AL6, which are representatives of three different subclasses. Based on two independent replicates of ChIP-seq data, we found that all the tested AL proteins formed distinct peaks at regions proximal to the transcription start site (TSS) (Supplemental Figure 18). The ChIP-seq signals of the three subclasses of AL proteins were shown to have high correlation and indistinguishable, indicating that they co-occupy chromatin at the whole-genome level (Supplemental Figure 18, Supplemental Dataset 2).

We selected AL2, AL4 and AL6 as representatives of the three subclasses of AL proteins for further investigation. Our ChIP-seq analysis identified 16,220, 20,688, and 16,523 genes enriched with AL2, AL4, and AL6, respectively, with the majority of them overlapping (Figure 4A). These overlapping genes were designated as AL-enriched genes (Figure 4A). Notably, the peaks of AL2-, AL4-, and AL6 were primarily located at the TSS-proximal regions and the intragenic regions (Figure 4B). Theses peaks exhibited high levels of H3K4me3, H3K36me3, H3K9Ac, and H2A.Z, which are associated with transcriptional activation, while showing low levels of H3K27me3 associated with transcriptional repression (Figure 4C). Intriguingly, the AL2, AL4, and AL6 peaks also displayed high levels of H2Aub (Figure 4C), which is thought to be closely associated with H3K27me3 (Zhou et al., 2017; Yin et al., 2023).

**Figure 4.**
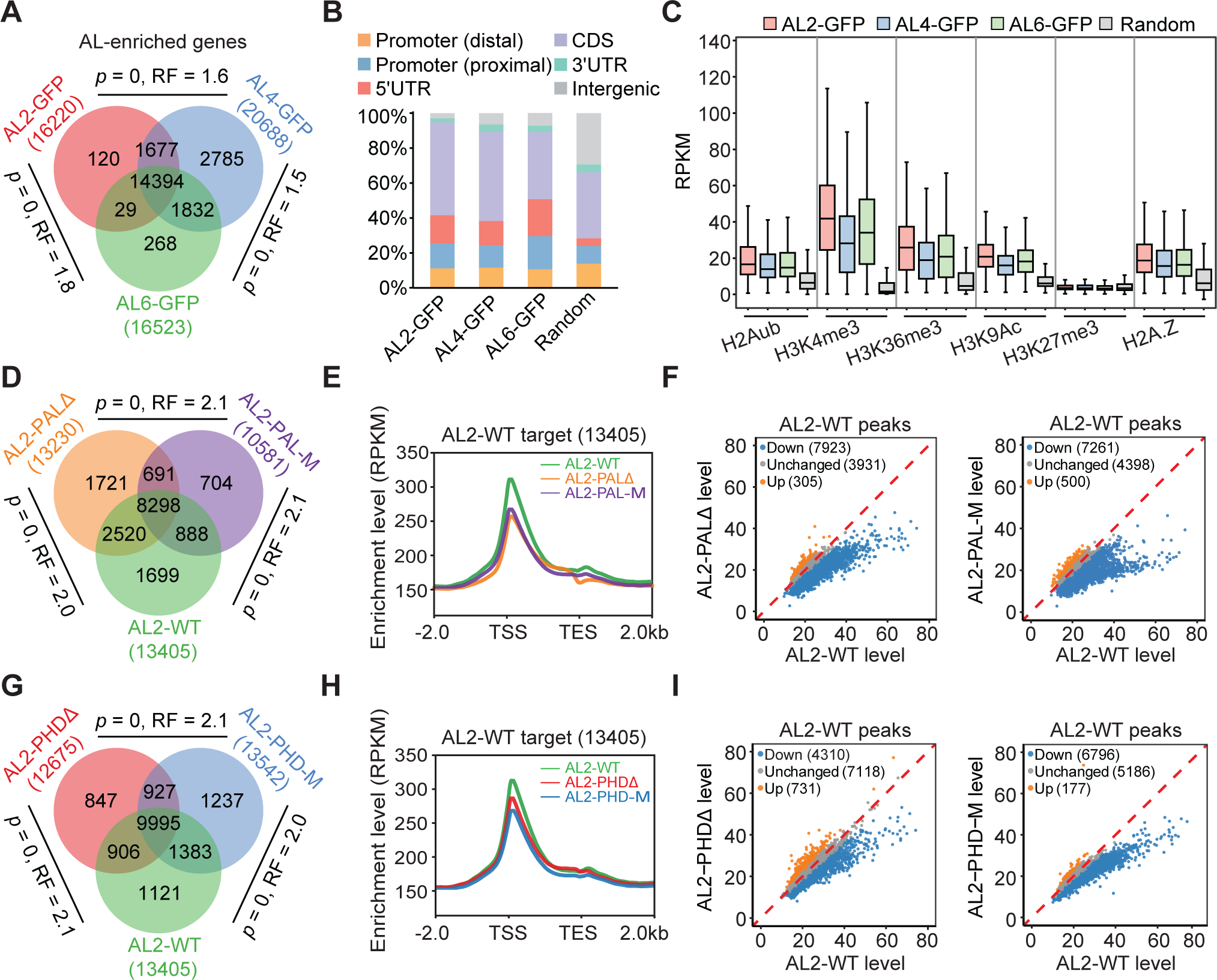
Involvement of the PHD and PAL domains in the association of AL proteins with chromatin. **(A)** The overlap between AL2-, AL4- and AL6-enriched genes. Genes common to all three groups are termed AL-enriched genes. *P* values were determined by one-tailed hypergeometric test. The representation factor (RF) represents the ratio of the observed number of overlapping genes to the expected number drawn from two independent groups. The ChIP-seq data of AL2-GFP, AL4-GFP and AL6-GFP are derived from two independent biological replicates. **(B)** Distribution of AL2, AL4 and AL6 ChIP-seq peaks in distal promoter, proximal promoter, 5’ UTR, CDS (coding sequence), 3’ UTR, and intergenic regions. Distal and proximal promoters represent 401-1,000 bp and 1-400 bp upstream of transcription start sites, respectively. The distribution of random genomic regions is shown as a control. **(C)** Box plots showing the levels of H2Aub, H3K4me3, H3K36me3, H3K9Ac, H3K27me3, H2A.Z in genomic regions bound by AL2, AL4 and AL6. Random: 10,000 random genomic regions. In the box plots, the center lines and box edges represent the medians and interquartile range (IQR), respectively. Whiskers extend to values within 1.5 times the IQR. **(D)** The overlap between AL2-WT, AL2-PALΔ and AL2-PAL-M enriched genes. *P* values were determined by one-tailed hypergeometric test. RF represents the ratio of the observed number of overlapping genes to the number expected by chance. The ChIP-seq data of AL2-WT, AL2-PALΔ and AL2-PAL-M are from two independent biological replicates. **(E)** Meta plots showing the whole-genomic enrichment levels of AL2-WT, AL2-PALΔ and AL2-PAL-M at AL2-WT target genes. TSS, transcription start site; TES, transcription end site. **(F)** Scatter plots showing the effect of AL2-PALΔ or AL2-PAL-M on the enrichment level of AL2 at AL2-bound genomic regions. Blue, orange, and gray dots represent down-regulated, up-regulated, and unchanged regions, respectively. **(G)** The overlap between AL2-WT, AL2-PHDΔ, and AL2-PHD-M enriched genes. *P* values were determined by one-tailed hypergeometric test. RF represents the ratio of the observed number of overlapping genes to the number expected by chance. The ChIP-seq data of AL2-WT, AL2-PHDΔ and AL2-PHD-M are from two independent biological replicates. **(H)** Meta plots showing the whole-genomic enrichment levels of AL2-WT, AL2-PHDΔ and AL2-PHD-M at AL2-bound genomic regions. TSS, transcription start site; TES, transcription end site. **(I)** Scatter plots showing the effect of AL2-PHDΔ or AL2-PHD-M on the enrichment level of AL2 at AL2-bound genomic regions. Blue, orange, and gray dots represent down-regulated, up-regulated, and unchanged regions, respectively.

### Both the PHD and PAL domains are involved in the association of AL proteins with chromatin

Given that the PHD finger of AL proteins binds to H3K4me3 and is crucial for their function in Arabidopsis plants (Figure 3H, 3I), we conducted ChIP-seq using *AL2-PHD*Δ and *AL2-PHD-M* transgenic plants to investigate the involvement of the PHD finger in the association of AL2 with chromatin. Our findings revealed that, although genes enriched with AL2-PHDΔ, AL2-PHD-M, and AL2-WT showed significant overlap (Figure 4G), the deletion or mutation of the PHD finger led to a reduction in the average enrichment of AL2 on its target genes at the whole-genome level (Figure 4H). Moreover, the deletion or mutation of the PHD finger significantly impaired the enrichment of AL2 at a substantial subset of its target loci (Figure 4I), suggesting that the PHD finger plays a crucial role in the association of AL proteins with chromatin. We further analyzed whether the PHD finger contributes to the association of AL proteins with H3K4me3-enriched genes. Our ChIP-seq analysis revealed that the deletion or mutation of the PHD finger tends to reduce the association of AL2 with a subset of AL2 target genes that exhibited relatively high levels of H3K4me3 (Supplemental Figure 19). These findings support the notion that the PHD finger recognizes H3K4me3 and thereby facilitates the association of AL proteins with H3K4me3-enriched target genes.

The interaction of the PAL domain of AL proteins with various chromatin-related proteins prompted us to explore whether AL proteins associate with chromatin through interacting with other chromatin-related proteins in a PAL-dependent manner. To investigate this, we carried out ChIP-seq using *AL2-PAL*Δ, *AL2-PAL-M*, and *AL2-WT* transgenic plants to examine the effect of PAL deletion or mutation on the association of AL2 with chromatin. Although the majority of genes enriched with AL2-PALΔ, AL2-PAL-M, and AL2-WT exhibited significant overlap (Figure 4D), the average enrichment of AL2-PALΔ and AL2-PAL-M was notably lower than that of AL2-WT at the whole-genome level (Figure 4E, 4F). These results suggest that the PAL domain is involved in the association of AL2 with chromatin, supporting the notion that the interaction of the PAL domain with multiple chromatin-related proteins contributes to the association of AL proteins with chromatin.

### Comparison of the genome-wide distributions of AL proteins and RING1A

Previous *in vitro* assays showed that the PAL domain of AL proteins directly interacts with RING1A and BMI1B (Molitor et al., 2014; Peng et al., 2018). Our pull-down assay not only confirmed the interaction of the AL2 PAL domain with RING1A and BMI1B (Supplemental Figure 16), but also identified that the conserved RING-finger domain of RING1A and BMI1B mediates the RING1A-BMI1B interaction (Supplemental Figure 16). These results suggest that the Arabidopsis PRC1 complex is composed of AL, RING1 and BMI1 proteins.

We conducted ChIP-seq using *RING1A-FLAG* transgenic plants and then compared the distributions of AL proteins and RING1A on chromatin throughout the Arabidopsis genome. A correlation analysis of the ChIP-seq data revealed a high positive correlation between RING1A and AL proteins (Figure 5A, Supplemental Dataset 2). Similar to AL proteins, RING1A was primarily enriched at the TSS-proximal regions and intragenic regions (Figure 5C). The majority (5,252/6,125 = 85.7 %) of RING1A-enriched genes were also enriched with AL proteins (Figure 5B), indicating that AL proteins and RING1A co-occupy PCR1 target genes at the whole-genome level. Notably, a substantial subset (9,142/14,394) of AL-enriched genes were not enriched by RING1A, which is consistent with the finding that AL proteins also interact with other chromatin-related proteins and have a PRC1-independent role.

**Figure 5.**
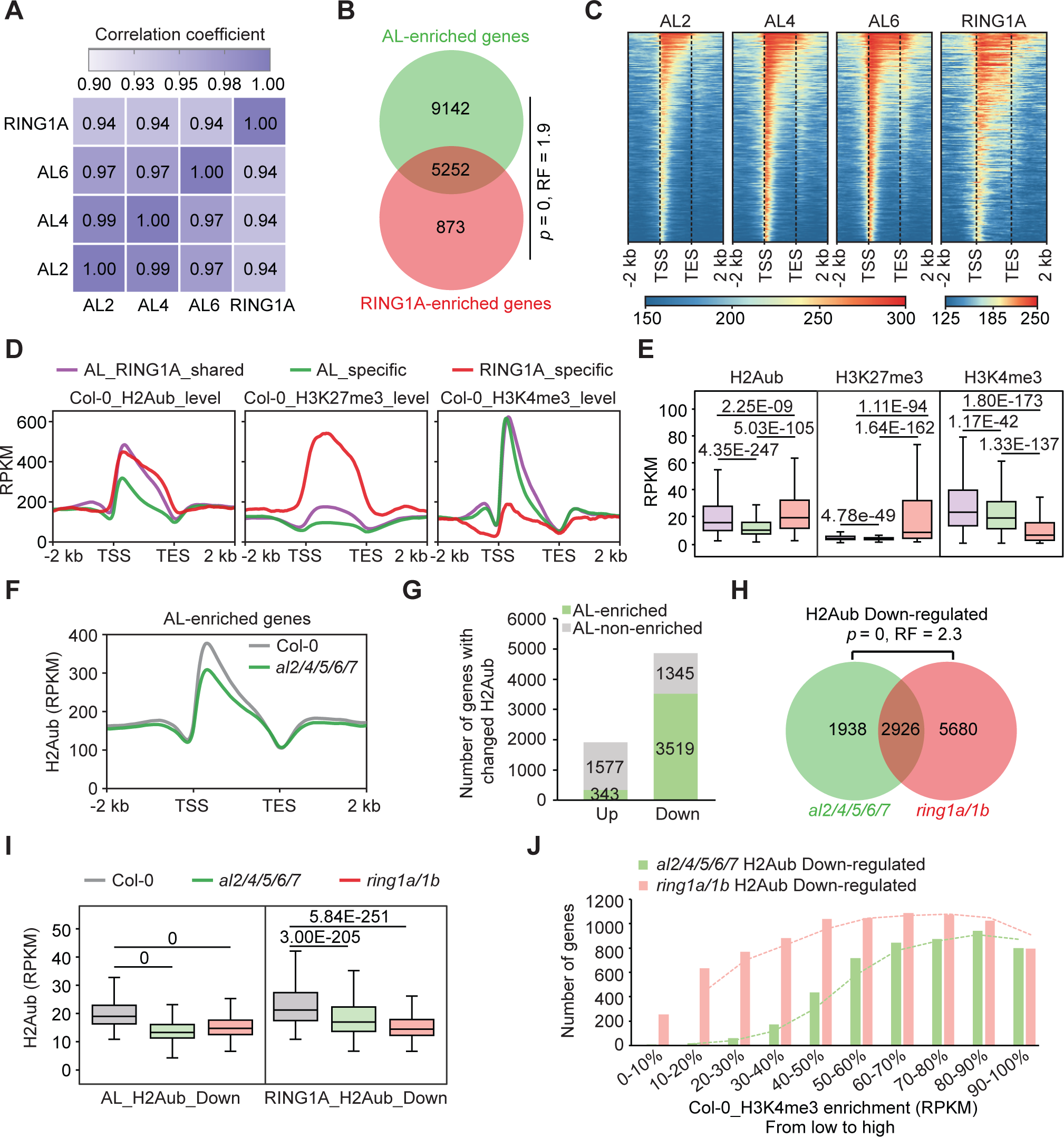
AL proteins are essential for PRC1-mediated H2Aub. **(A)** Correlation analysis of ChIP-seq signals between pairwise AL2, AL4, AL6 and RING1A. Pearson correlation coefficients are shown. The ChIP-seq data of AL2-GFP, AL4-GFP, AL6-GFP and RING1A-FLAG are from two independent biological replicates. **(B)** The overlap between AL-enriched genes and RING1A-enriched genes. Statistical significance was determined by one-tailed the hypergeometric test. RF represents the ratio of the observed number of overlapping genes to the number expected by chance. The ChIP-seq data of RING1A are from two independent biological replicates. **(C)** Heatmap showing the enrichment levels of AL2, AL4, AL6, and RING1A at their respective target genes. The genes are organized in descending order of their enrichment levels, from the highest at the top to the lowest at the bottom. **(D)** Meta plots showing the distribution patterns of H2Aub, H3K27me3 and H3K4me3 across AL- and RING1A-shared genes, AL-specific genes, and RING1A-specific genes. TSS, transcription start site; TES, transcription end site. **(E)** The relative enrichment levels of H2Aub, H3K27me3 and H3K4me3 at AL-RING1A-shared genes, AL-specific genes, and RING1A-specific genes. In the box plots, the center lines and box edges represent the medians and interquartile range (IQR), respectively. Whiskers extend to values within 1.5 times the IQR. Purple, green, and red boxes represent the histone modification levels for AL and RING1A-shared genes, AL-specific genes, and RING1A-specific genes, respectively. *P* values were determined by two-tailed Mann Whitney U test (no-paired). **(F)** Meta plots showing the distribution patterns of H2Aub in wild type and *al2/4/5/6/7* mutant at AL-enriched genes. TSS, transcription start site; TES, transcription end site. **(G)** The numbers of genes with up- and down-regulated H2Aub levels in *al2/4/5/6/7* mutant relative to the wild type. Green represents AL-enriched genes, while gray indicates AL-non-enriched genes. **(H)** The overlap of genes with down-regulated H2Aub levels in *al2/4/5/6/7* and *ring1a/1b* relative to the wild type. *P* values were determined by hypergeometric test. RF is the ratio of the number of observed overlapping genes to the number expected by chance. **(I)** Box plots showing the H2Aub levels in wild type, *al2/4/5/6/7* and *ring1a/1b* mutants at AL_H2Aub_down and RING1A_H2Aub_down sites. AL_H2Aub_down refers to AL-enriched genes with H2Aub down-regulation in *al2/4/5/6/7* (3,519 genes), and RING1A_H2Aub_down refers to RING1A-enriched genes with H2Aub down-regulation in *ring1a/1b* H2Aub (1,486 genes). In the box plots, the center lines and box edges represent the medians and interquartile range (IQR), respectively. Whiskers extend to values within 1.5 times the IQR. *P* values were determined by two-tailed Mann Whitney U test (paired). **(J)** The graph illustrating the count of genes exhibiting reduced levels of H2Aub levels in the *al2/4/5/6/7* and *ring1a/1b* mutants within deciles of increasing H3K4me3 levels. The total number of Arabidopsis genes (*n* = 32548) were divided into the deciles with ascending levels of H3K4me3.

### AL proteins play a crucial role in PRC1-mediated H2Aub

We compared the genes enriched with AL proteins and RING1A and classified them into AL- and RING1A-shared genes, AL-specific genes, and RING1A-specific genes. We then examined the H3K4me3, H3K27me3, and H2Aub levels of these genes in the wild type. Our analyses revealed that AL- and RING1A-shared genes, as well as AL-specific genes, exhibited high levels of H3K4me3 compared to RING1A-specific genes, supporting the notion that AL proteins have the ability to bind H3K4me3. Consistent with the previously observed mutual exclusivity of H3K27me3 and H3K4me3 at numerous genes in Arabidopsis plants (Zhang et al., 2009; Roudier et al., 2014; Zhu et al., 2023), AL- and RING1A-shared genes and AL-specific genes displayed extremely low levels of H3K27me3 compared to RING1A-specific genes (Figure 5D, 5E). The level of H2Aub was significantly higher in AL- and RING1A-shared genes, as well as RING1A-specific genes, than in AL-specific genes (Figure 5D, 5E). Given that the PRC1 complex is responsible for catalyzing H2Aub (Cao et al., 2005; Yang et al., 2013), we predict that AL proteins facilitate the accumulation of H2Aub at H3K4me3-enriched genes and thereby link the two histone modifications in Arabidopsis.

We therefore performed ChIP-seq for H2Aub in the wild type, *al2/4/5/6/7*, and *ring1a/1b* mutants to investigate the involvement of AL proteins in PRC1-mediated H2Aub. We found that the average H2Aub level was significantly reduced in the *al2/4/5/6/7* mutant as well as in the *ring1a/1b* mutant compared to the wild type (Figure 5F, Supplemental Figure 20). In total, we identified 4,864 genes with decreased H2Aub levels (FC < 0.8, FDR < 0.05) and 1,920 genes with increased H2Aub levels (FC > 1.2, FDR < 0.05) in the *al2/4/5/6/7* mutant compared to the wild type (Figure 5G, Supplemental Dataset 3). We found that the majority (3,519/4,864) of genes with decreased H2Aub levels are AL-enriched genes, while only the minority (343/1,920) of genes with increased H2Aub levels are AL-enriched genes (Figure 5G). Moreover, AL-enriched genes with decreased levels of H2Aub are more abundant than those with increased levels of H2Aub in the *al2/4/5/6/7* mutant, which is consistent with the finding that RING1A-enriched genes exhibiting decreased levels of H2Aub are more abundant than those with increased levels of H2Aub in the *ring1a/1b* mutant (Supplemental Figure 20). Consistent with the conserved role of the PRC1 components RING1A/1B in mediating H2Aub, our H2Aub ChIP-seq analysis identified 8,606 genes with decreased H2Aub levels and only 754 genes with increased H2Aub levels (Figure 5H, Supplemental Dataset 3). Although genes with decreased levels of H2Aub identified in the *al2/4/5/6/7* mutant are less abundant than those identified in the *ring1a/1b* mutant, the majority of genes with decreased levels of H2Aub in the *al2/4/5/6/7* mutant also showed decreased levels of H2Aub in the *ring1a/1b* mutant (Figure 5I). The overall H2Aub levels of both AL- and RING1A-target genes were co-regulated in the *al2/4/5/6/7* and *ring1a/1b* mutants (Figure 5I). The impact of *al2/4/5/6/7* mutations on H2Aub levels was clearly evidenced at representative H2Aub target loci, revealing both AL-dependent and -independent H2Aub target loci (Supplemental Figure 20). These analyses indicate that AL proteins are involved in mediating H2Aub at a subset of PRC1 target loci.

To investigate the potential correlation between H2Aub and H3K4me3 levels across the Arabidopsis genome, we quantified the levels of H2Aub and H3K4me3 for all Arabidopsis genes in wild-type plants. After categorizing these genes into deciles based on increasing H2Aub levels, we observed that last four deciles with increasing H2Aub also exhibited a consistent rise in H3K4me3 levels, indicating a genome-wide positive correlation between H2Aub and H3K4me3 (Supplemental Figure 21). Subsequently, we examined the enrichment levels of H3K4me3 in genes with reduced H2Aub in the *al2/4/5/6/7* and *ring1a/1b* mutants. Our analysis revealed that genes with decreased H2Aub levels in the *ring1a/1b* mutant were found in regions with varying levels of H3K4me3 enrichment, while genes exhibiting reduced H2Aub in the *al2/4/5/6/7* mutant were enriched at genes with high H3K4me3 levels (Figure 5J). These analyses suggest that AL proteins facilitate H2Aub at a subset of PRC1 target genes with high levels of H3K4me3, and thus reveal a crucial molecular mechanism underlying the genome-wide connection between H2Aub and H3K4me3.

### AL-mediated H2Aub and REF6-mediated H3K27me3 demethylation

Our AP-MS results indicated that AL proteins can interact the histone H3K27 demethylase REF6 in Arabidopsis plants (Figure 2A, Supplemental Dataset 1). We confirmed the interaction by pull-down assays and demonstrated that the C-terminal Zn-finger-containing domain of REF6 (REF6-3: 853-1360 aa) interacts with the AL2-PAL domain (Figure 6A). To understand how AL proteins coordinate with REF6 to regulate the chromatin state, we analyzed the previous REF6 ChIP-seq data (Cui et al., 2016) and compared the distribution of AL proteins and REF6 throughout the Arabidopsis genome. We found that 60.0% of (1,702/2,836) of REF6-enriched genes overlapped with AL-enriched genes (Figure 6B). By evaluating the H3K4me3 level at different subsets of AL- and REF6-enriched genes, we observed that AL- and REF6-shared genes and AL-specific genes had a high level of H3K4me3 compared to AL- and REF6-specific genes (Figure 6C). This finding is consistent with the concept that the binding of AL proteins to H3K4me3 contributes to their association with chromatin.

**Figure 6.**
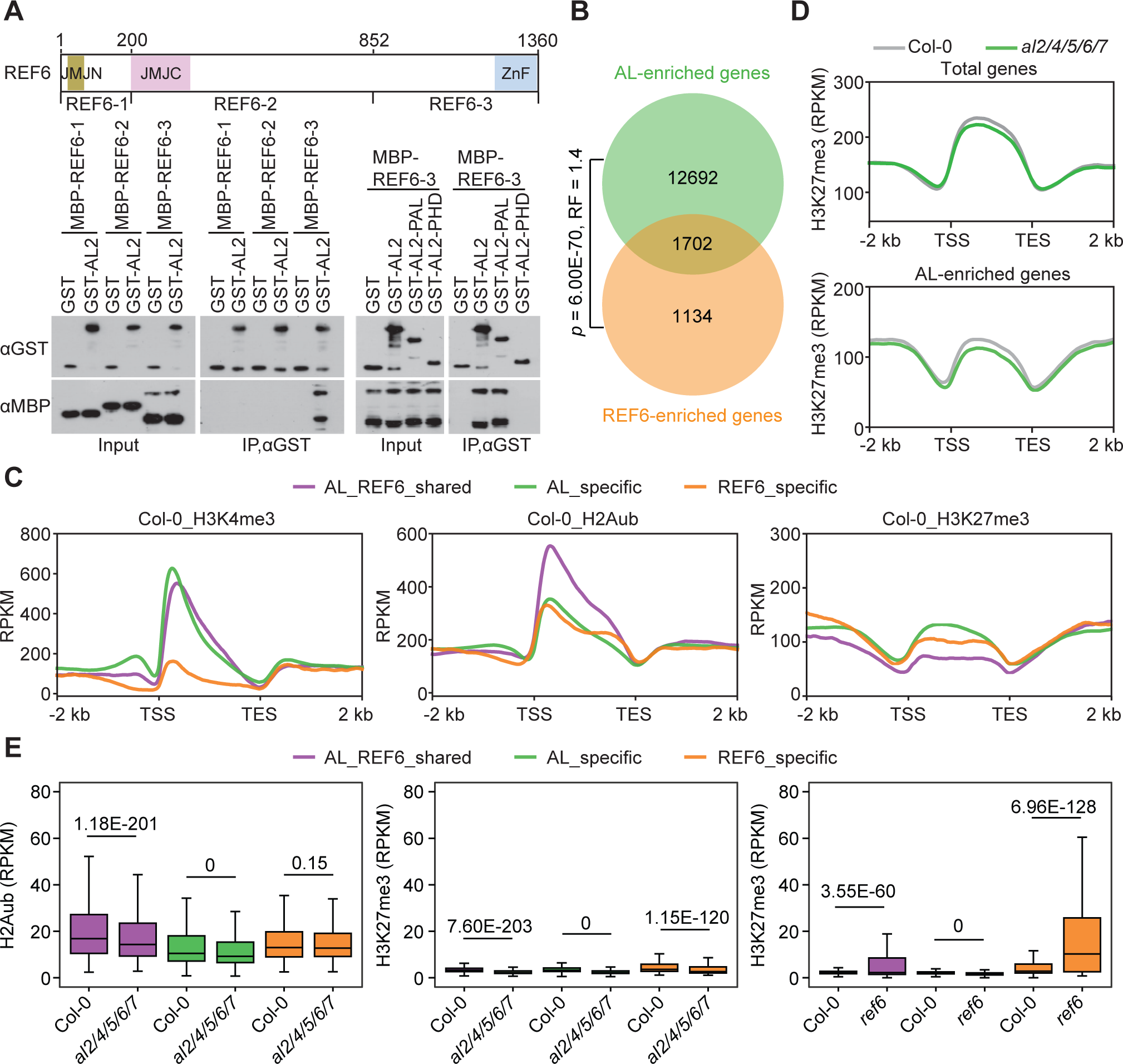
Integrated analysis of AL-mediated H2Aub and REF6-mediated H3K27 demethylation. **(A)** Identification of REF6 and AL2 interaction domains by pull-down assays. The schematic diagram of full-length and truncated versions of REF6 (upper panel) and the pull-down results (lower panel) are shown. **(B)** Analysis of overlapping genes enriched by AL and REF6. Statistical significance was determined by one-tailed hypergeometric test. RF represents the observed number of overlapping genes to the number expected by chance. **(C)** Meta plots showing the distribution patterns of H3K4me3, H2Aub, and H3K27me3 across AL- and REF6-shared genes, AL-specific genes, and REF6-specific genes. TSS, transcription start site; TES, transcription end site. **(D)** Meta plots showing the distribution patterns of H3K27me3 in wild type and *al2/4/5/6/7* mutant across total Arabidopsis genes and AL-enriched genes. TSS, transcription start site; TES, transcription end site. **(E)** Box plots illustrating the levels of H2Aub (left) and H3K27me3 (middle) in wild type and *al2/4/5/6/7* mutant, as well as the levels of H3K27me3 (right) in wild type and *ref6* mutant. In the box plots, the center lines and box edges represent the medians and interquartile range (IQR), respectively. Whiskers extend to values within 1.5 times the IQR. *P* values were determined two-tailed Mann Whitney U test (paired).

Considering that AL proteins and REF6 are involved in H2Aub deposition and H3K27 demethylation, respectively, we performed ChIP-seq for H2Aub and H3K27me3 in the *al2/4/5/6/7* mutant and the wild type to determine whether AL proteins cooperate with REF6 to regulate the histone modifications (Figure 6D, Supplemental Dataset 3, 4). Our analysis revealed that AL- and REF6-shared genes have a high level of H2Aub compared to AL- and REF6-specific genes (Figure 6C), indicating that the high level of H2Aub is associated with the presence of REF6 at AL- and REF6-shared genes. Furthermore, we found that the H2Aub level was significantly reduced in the *al2/4/5/6/7* mutant relative to the wild type at both AL- and REF6-shared genes and AL-specific genes but not at REF6-specific genes (Figure 6E). These results suggest that REF6 is dispensable for AL-mediated H2Aub and that the presence of REF6 is related to the high level of H2Aub on chromatin.

The analysis of H3K27me3 in the wild type plants indicated that both AL- and REF6-target genes exhibited a low level of H3K27me3 (Figure 6C). However, in the *ref6* mutant, the H3K27me3 level was significantly increased compared to the wild type at AL- and REF6-shared regions and REF6-specific regions, but not at AL-specific regions (Figure 6E). This finding supports the crucial role of REF6 as an H3K27me3 demethylase. Interestingly, in the *al2/4/5/6/7* mutant, the H3K27me3 level did not increase relative to the wild type in any of the three gene classes (Figure 6E). These results highlight the coordination between AL-mediated H2Aub and REF6-mediated histone H3K27me3 demethylation.

### AL proteins play a regulatory role in H2A.Z distribution

Our AP-MS data indicated that the SWR1 complex components, involved in H2A.Z deposition, were co-purified with AL proteins (Figure 2A, Supplemental Dataset 1). This finding is consistent with previous studies demonstrating the co-purification of AL proteins with SWR1 complex components (Supplemental Figure 10) (Potok et al., 2019; Luo et al., 2020). In our pull-down assay, we observed a direct interaction between the SWR1 catalytic subunit PIE1 and the PAL domain of AL2 through its N-terminal HSA-containing domain (PIE1-1: 1-520 aa) (Figure 7A). Interestingly, although our AP-MS indicated an interaction between AL proteins and CHR11, a catalytic subunit of ISWI complex that is involved in nucleosome sliding (Li et al., 2014; Luo et al., 2020), the pull-down assay did not detect a direct interaction between AL proteins and CHR11 (Supplemental Figure 15). This suggests that the interaction between AL proteins and CHR11 is indirect. Considering that CHR11 functions as a subunit of both the ISWI complex and the SWR1 complex (Luo et al., 2020), the direct interaction between AL proteins and the SWR1 catalytic subunit PIE1 may be partially responsible for the interaction of AL proteins and CHR11.

**Figure 7.**
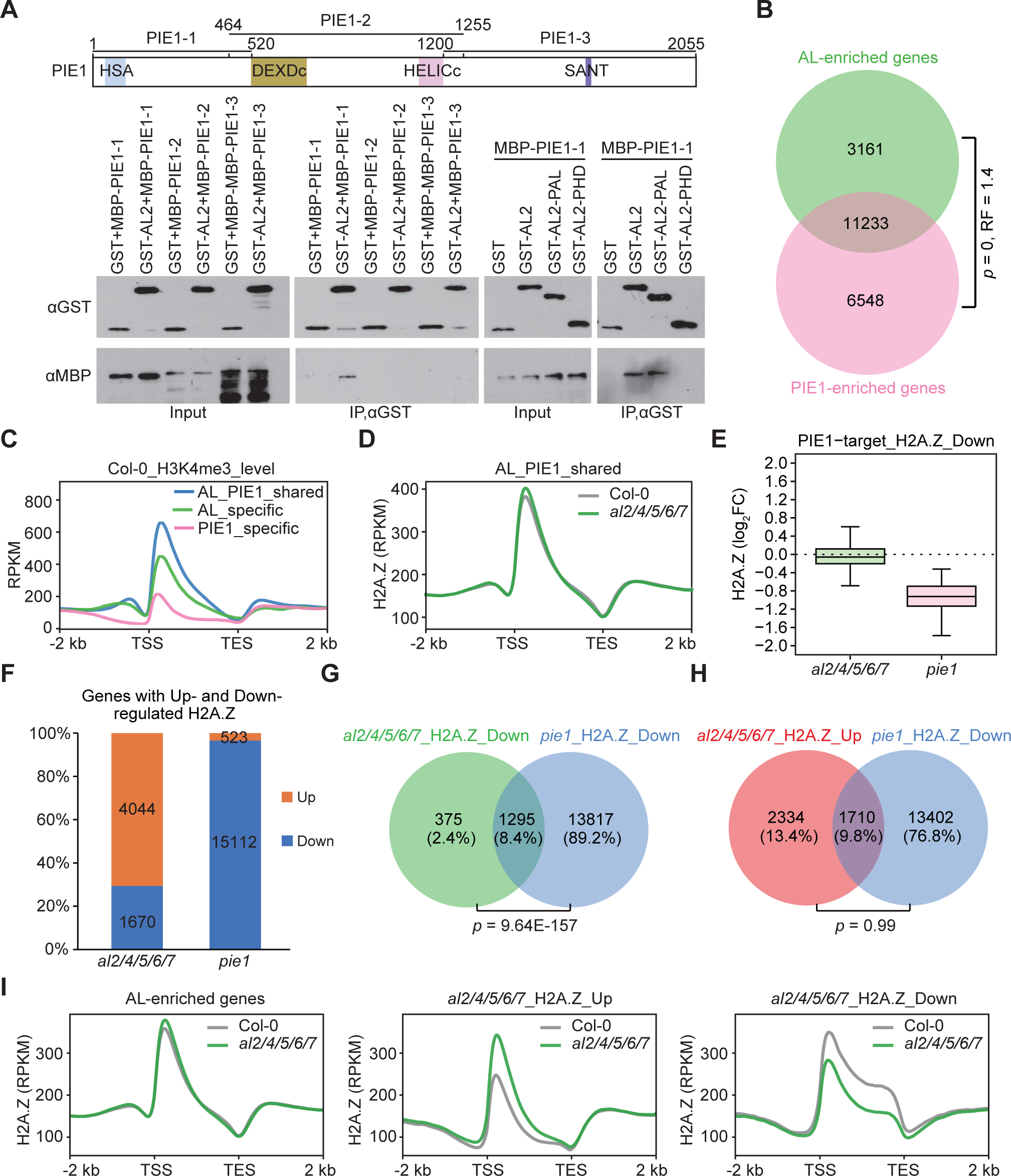
AL proteins play a regulatory role in the distribution of H2A.Z. **(A)** Determination of the interaction domains of PIE1 and AL2 by pull-down assays. The schematic representation of full-length and truncated versions of PIE1 is shown on the top, while the pull-down results are shown at the bottom. **(B)** The overlap between AL-enriched genes and PIE1-enriched genes. Statistical significance was determined by one-tailed hypergeometric test. RF represents the ratio of the observed number of overlapping genes to the number expected by chance. **(C)** Meta plots showing the distribution patterns of H3K4me3 in wild type across AL- and PIE1-shared genes, AL-specific genes, and PIE1-specific genes. TSS, transcription start site; TES, transcription end site. **(D)** Meta plots showing the distribution patterns of H2A.Z in wild type and *al2/4/5/6/7* mutant at AL- and PIE1-shared genes. TSS, transcription start site; TES, transcription end site. **(E)** Box plots illustrating the impact of *al2/4/5/6/7* and *pie1* on the levels of H2A.Z at PIE1 target genes. PIE1 target genes exhibiting down-regulated levels of H2A.Z in the *pie1* mutant were subjected to the analysis. In the box plots, the center lines and box edges represent the medians and interquartile range (IQR), respectively. Whiskers extend to values within 1.5 times the IQR. **(F)** The counts of genes with up- and down-regulated levels of H2A.Z in the *al2/4/5/6/7* and *pie1* mutants relative to the wild type. **(G, H)** Determination of the overlap between genes with down-regulated H2A.Z levels in the *al2/4/5/6/7* mutant and those with down-regulated H2A.Z levels in the *pie1* mutant **(G)**, and between genes with up-regulated H2A.Z levels in the *al2/4/5/6/7* mutant and those with down-regulated H2A.Z level in the *pie1* mutant **(H)**. *P* values were determined by one-tailed hypergeometric test. **(I)** Meta plots showing the distribution patterns of H2A.Z in wild type and *al2/4/5/6/7* mutant at AL-enriched genes (left) and genes with increased H2A.Z levels (middle) and with decreased H2A.Z levels (right) in the *al2/4/5/6/7* mutant. TSS, transcription start site; TES, transcription end site.

To investigate the occupancy of AL proteins at SWR1 target genes on a genome-wide scale, we compared our ChIP-seq data for AL proteins with previously published ChIP-seq data for the SWR1 catalytic subunit PIE1 (Luo et al., 2020). Our analysis revealed a significant overlap between AL- and PIE1-enriched genes (*p* = 0; RF = 1.4), with the majority (11,233/17,781) of PIE1-enriched genes also showing enrichment with AL proteins (Figure 7B). Furthermore, when examining the H3K4me3 levels across different classes of AL- and PIE1-enriched genes independently, we observed that AL- and PIE1-shared genes, as well as AL-specific genes, displayed higher levels of H3K4me3 compared to PIE1-specific genes (Figure 7C). This suggests a preference of AL proteins for binding to H3K4me3-enriched PIE1 target genes, which is consistent with their known ability to bind to H3K4me3. Additionally, we performed ChIP-seq for H2A.Z in *al2/4/5/6/7* mutant and wild-type plants (Supplemental Dataset 5). In the wild type, H2A.Z ChIP-seq signals detected in the current study highly overlapped with those identified in our previous study (Luo et al., 2020). Although the overall H2A.Z level of PIE1 target genes was substantially reduced in the *pie1* mutant (Luo et al., 2020), it did not show a significant reduction in the *al2/4/5/6/7* mutant (Figure 7D, 7E). This suggests that AL proteins are unlikely to function as canonical subunits of SWR1 complex in mediating H2A.Z deposition.

Our ChIP-seq analysis identified 4,044 genes with increased H2A.Z levels (FC > 1.2, FDR < 0.05) and 1,670 genes with decreased H2A.Z levels (FC < 0.8, FDR < 0.05) in *al2/4/5/6/7* mutant compared to the wild type (Figure 7F, Supplemental Dataset 5). Based on previous ChIP-seq data (Luo et al., 2020), 15,112 genes exhibited reduced H2A.Z levels in the *pie1* mutant (Figure 7F). Although the number of genes with reduced H2A.Z levels is much lower in the *al2/4/5/6/7* mutant than in *pie1* mutant, the majority (1,670/5,714) of genes with reduced H2A.Z levels in the *al2/4/5/6/7* mutant also exhibited reduced levels of H2A.Z in the *pie1* mutant (Figure 7G), suggesting that AL proteins contribute to H2A.Z deposition at a small subset of PIE1 target genes. The genes with increased H2A.Z levels in the *al2/4/5/6/7* mutant do not significantly overlap with the genes with a reduced H2A.Z level in the *pie1* mutant (*p* = 0.99, hypergeometric test) (Figure 7H), indicating that AL proteins have a negative effect on H2A.Z deposition in a PIE1-independent manner. Given that the INO80 complex is reported to mediate the exchanging of H2A.Z for the canonical H2A (Papamichos-Chronakis et al., 2011; Willige et al., 2021; Xue et al., 2021), the negative impact of AL proteins on H2A.Z deposition is likely related to their interaction with the INO80 complex components, as determined by the current study (Figure 2A, Supplemental Figure 9). Interestingly, we observed that at genes with increased H2A.Z levels in the *al2/4/5/6/7* mutant, H2A.Z formed a sharp peak proximal to the TSS (Figure 8I). In contrast, at genes with decreased H2A.Z levels in the *al2/4/5/6/7* mutant, H2A.Z was enriched not only at the TSS-proximal region but also at the intragenic region (Figure 8, Supplemental Figure 22). These findings suggest that AL proteins differentially affect H2A.Z deposition at the TSS-proximal region and at the intragenic region.

**Figure 8.**
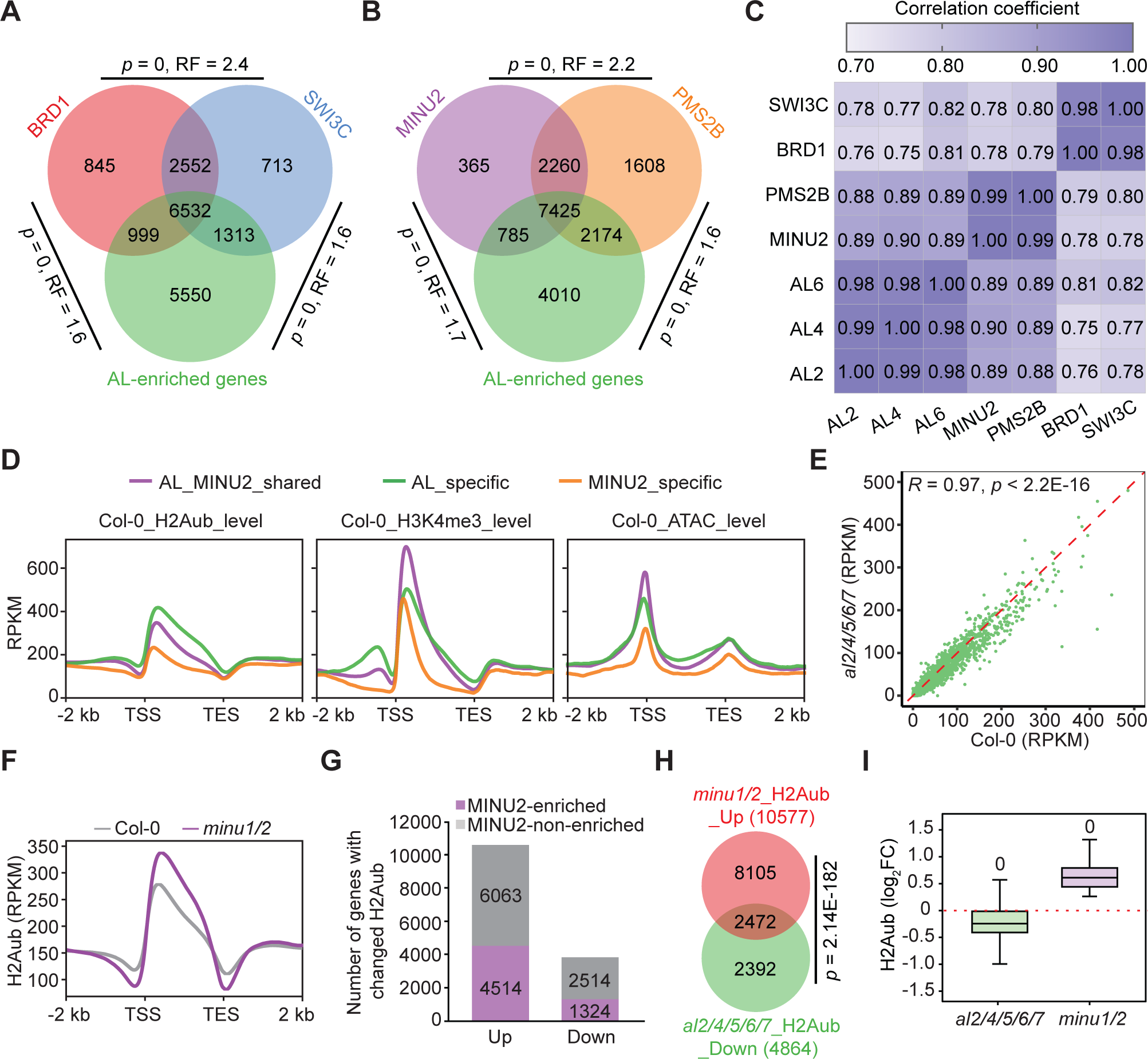
Analysis of coordination between AL proteins and SWI/SNF complexes. **(A,B)** Examination of the overlap between genes enriched with AL proteins and those enriched with the BAS complex **(A)** or the MAS complex **(B)**. *P* values were determined by one-tailed hypergeometric test. RF denotes the proportion of overlapping genes observed in comparison to the number anticipated by random chance. Genes enriched with the BAS complex components BRD1 and SWI3C are termed BAS-enriched genes, while genes enriched with the MAS complex components MINU2 and PMS2B are referred to as MAS-enriched genes. **(C)** Correlation analysis of ChIP-seq signals between pairwise comparisons of AL2, AL4, AL6, MINU2, PMS2B, BRD1 and SWI3C. Pearson correlation coefficients are shown. **(D)** Meta plots showing the distribution patterns of H2Aub, H3K4me3 and ATAC-seq signals in wild type across AL- and MINU2-shared target genes, AL-specific target genes, and MINU2-specific target genes. TSS, transcription start site; TES, transcription end site. **(E)** Scatter plots illustrating the comparison of accessibility signals between the *al2/4/5/6/7* mutant and wild type based on the whole-genomic level analysis. **(F)** Meta plots showing the distribution of H2Aub in wild type and *minu1/2* mutant across total Arabidopsis genes. TSS, transcription start site; TES, transcription end site. **(G)** The counts of MINU2-enriched genes and MINU2-non-enriched genes among those exhibiting up- and down-regulated H2Aub levels in the *minu1/2* mutant relative to wild type. **(H)** Analysis of the overlap between genes with down-regulated H2Aub levels in the *al2/4/5/6/7* mutant and those with up-regulated H2Aub levels in the *minu1/2* mutant. Statistical significance was determined by one-tailed hypergeometric test. **(I)** Box plots showing the effect of *minu1/2* on the levels of H2Aub in a subset of AL-enriched genes that exhibit reduced H2Aub levels in the *al2/4/5/6/7* mutant relative to wild type. In the box plots, the center lines and box edges represent the medians and interquartile range (IQR), respectively. Whiskers extend to values within 1.5 times the IQR.

### The coordination between AL proteins and SWI/SNF complex

Our AP-MS results indicated that the components of the BAS and MAS-type SWI/SNF complexes, but not of the SAS-type SWI/SNF complex, were co-purified with AL proteins (Figure 2A, Supplemental Dataset 1). Furthermore, previous AP-MS data (Guo et al., 2022) also showed that AL proteins were co-purified with the SWI/SNF complex components, including BRM, BRIP2, SWP73A, BSH, SWI3A, and ARP4 (Supplemental Figure 9). We conducted pull-down assays to determine whether AL proteins interact with truncated versions of the BAS catalytic enzyme BRM (BRM-1: 1-744 aa, BRM-2: 718-1,611aa, BRM-3: 1,596-2,193aa) and of the MAS catalytic enzyme MINU2 (MINU2-1: 1-381aa, MINU2-2:382-676, MINU2-3: 677-1,064 aa). The pull-down assays indicated that AL proteins interact with the C-terminal domain of both BRM (BRM-3) and MINU2 (MINU2-3) (Supplemental Figure 11), confirming that AL proteins directly interact with BRM and MINU2.

By comparing the ChIP-seq data for AL proteins with our previous ChIP-seq data for the BAS components BRD1 and SWI3C and the MAS components MINU2 and PMS2B (Guo et al., 2022; Fu et al., 2023), we found that AL-enriched genes significantly overlapped with genes occupied by the BAS and MAS components (Figure 8A, 8B). A correlation analysis of the ChIP-seq data indicated that AL proteins are highly correlated with the MAS components and to a lesser extent with the BAS components, with the ChIP-seq signals of components within the same complex clustered together (Figure 8C). Moreover, we evaluated the H3K4me3 level of different classes of AL and MINU2 target genes and found that AL- and MINU2-shared target genes exhibited higher levels of H3K4me3 than AL- and MINU2-specific target genes (Figure 8D), which is consistent with notion that AL proteins are associated with H3K4me3-enriched MAS target genes.

Considering that the MAS complex is responsible for mediating chromatin accessibility (Guo et al., 2022), we performed assay for transposase-accessible chromatin followed by sequencing (ATAC–seq) in the *al2/4/5/6/7* mutant and the wild type to investigate whether AL proteins regulate chromatin accessibility. The ATAC-seq analysis did not identify a significant number of differentially accessible regions in *al2/4/5/6/7* mutant relative to the wild type (Figure 8E, Supplemental Dataset 6). This suggests that AL proteins do not function as canonical subunits of the MAS complex that regulates chromatin accessibility. Therefore, we performed ChIP-seq for H2Aub in the *minu1 minu2* (*minu1/2*) mutant and the wild type to determine whether the MAS complex affects AL-mediated H2Aub. Interestingly, we found that the genome-wide H2Aub level was substantially increased in the *minu1/2* mutant relative to the wild type (Figure 8F). In total, we identified 10,576 genes with increased H2Aub levels in the *minu1/2* mutant compared to the wild type (Figure 8G). The genes with increased H2Aub levels in the *minu1/2* mutant significantly overlapped with those with a reduced H2Aub level in the *al2/4/5/6/7* mutant (Figure 8H, 8I). These findings suggest that the MAS complex exhibits a previously uncharacterized role in H2A deubiquitination and that AL proteins and the MAS complex have antagonistic effects on H2Aub. The antagonistic effects are probably involved in maintaining the equilibrium of H2Aub at the whole-genome level.

## Discussion

Polycomb group (PcG) proteins are evolutionarily conserved epigenetic regulators that faciliate transcriptional gene repression in eukaryotes (Liu et al., 2021). Typically, PcG proteins form two distinct multi-protein complexes, PRC1 and PRC2, responsible for H2A monoubiquitination and H3K27 trimethylation, respectively (Molitor and Shen, 2013; Zhou et al., 2018; Barbour et al., 2020). In metazoans, the PRC1 complex contains an H3K27me3 reader, which mediates H2A monoubiquitination downstream to H3K27me3 (Wang et al., 2004; Molitor and Shen, 2013). However, in Arabidopsis, H3K27me3 is not required for establishing H2Aub at most of H2Aub target genes (Zhou et al., 2017; Liu et al., 2021). Additionally, unlike H3K27me3, H2Aub does not function as a canonical histone mark associated with transcriptional repression (Yin et al., 2021; Yin et al., 2023). Therefore, it is important to determine how the PRC1 complex mediates H2Aub selectively at a subset of protein-coding genes. In this study, we found that the H3K4me3-binding ability of AL proteins is involved in their association with H3K4me3-enriched genes at the whole-genome level. Furthermore, we demonstrated that AL proteins function as subunits of the PRC1 complex and are crucial for mediating H2A monoubiquitination in Arabidopsis plants. Considering the high positive correlation between the enrichments of H3K4me3 and H2Aub throughout the Arabidopsis genome, we suggest that the incorporation of H3K4me3-binding AL proteins into the PRC1 complex facilitates the link between H3K4me3 and H2Aub in plants.

Although we have demonstrated that the AL proteins function as subunits of the PRC1 complex and play a crucial role in PRC1-mediated H2Aub at the whole-genome level, it remains unclear how the AL proteins are involved in PRC1-mediated H2Aub. Since the AL proteins contain the conserved PHD finger with H3K4me3-binding ability, it is possible that the AL proteins mediate the recruitment of the PRC1 complex to H3K4me3-enriched genomic regions, thereby facilitating the link between H3K4me3 and H2Aub. However, our ChIP-seq did not support the role of the AL proteins in the association of RING1A with chromatin, suggesting that the AL proteins are involved in mediating H2Aub followed by the recruitment of the PRC1 complex to chromatin. Further studies are required to determine whether the AL proteins are crucial for facilitating the H2Aub activity of the PRC1 complex. Intriguingly, the AL proteins were also shown to interact with the PEAT complex, which is involved in H2A deubiquitination (Zheng et al., 2023). Within the PEAT complex, UBP5 is the enzyme responsible for the genome-wide H2A deubiquitination while the PWWP proteins are accessory subunits involved in H2A deubiquitination (Zheng et al., 2023). Our pull-down assay has demonstrated the direct interaction between the AL proteins and the PWWP proteins (Supplemental Figure 16). Therefore, it is also possible that the AL proteins promote H2Aub by suppressing the H2A deubiquitination activity of the PEAT complex. In future, biochemical and genomic analyses were required to clarify how the AL proteins are involved in the regulation of H2Aub at the whole-genome level.

RELATIVE OF EARLY FLOWERING 6 (REF6), a Jumonji-type histone H3K27 demethylase, is responsible for the demethylation of the repressive histone mark H3K27me3 (Lu et al., 2011). Previous studies have indicated that the zinc finger domain of REF6 is responsible for its target selection in a sequence-specific manner (Cui et al., 2016; Yan et al., 2018). Moreover, REF6 has also been shown to bind a subset of REF6 target loci in a zinc finger-independent manner (Yan et al., 2018). Although previous studies have indicated that active histone marks, including H3K4me3 and H3K9Ac, are enriched at REF6 target loci (Cui et al., 2016; Yan et al., 2018), how these active histone marks are associated with these loci remains largely unclear. Our study indicated that the AL proteins interact with REF6 (Figure 6A). Given that the AL proteins have the H3K4me3-binding ability *in vitro* and *in vivo*, the interaction between AL proteins and REF6 reveals a link between H3K4me3 and REF6-mediated H3K27 demethylation. However, because the AL proteins do not have a significant impact on H3K27me3 (Figure 6), the binding of AL proteins to H3K4me3 is unlikely to function upstream to REF6-medaited H3K27 demethylation. The finding of the enrichment of H3K4me3 at AL- and REF6-shared target genes suggests that the interaction between AL proteins and REF6 may contribute to the link between H3K4me3 and H3K27 demethylation (Figure 6C). Intriguingly, H2Aub is exclusively enriched at AL- and REF6-shared target genes but not at AL- or REF6-specific target genes, suggesting a potential role of REF6 in AL-dependent H2Aub (Figure 6C, 6E). Further studies are required to determine how the interaction between AL proteins and REF6 is involved in the coordination of H3K4me3, H2Aub, and H3K27 demethylation at their common target genes.

The histone H2A variant H2A.Z is deposited by the conserved SWR1 complex in eukaryotes (Mizuguchi et al., 2004; Kapoor and Shen, 2014). In Arabidopsis, the AL proteins were previously co-purified with the SWR1 complex component ARP6 (Potok et al., 2019), which is consistent with our AP-MS results (Figure 2A, 2B, Supplemental Figure 10, Supplemental Dataset 1). However, the potential role of AL proteins in regulating SWR1-mediated H2A.Z deposition remains unclear. Our pull-down assay showed that the AL proteins directly interact with the SWR1 catalytic subunit PIE1 (Figure 7A). Furthermore, our genome-wide analysis of H2A.Z indicated that the AL proteins do not function as canonical subunits of the SWR1 complex but are more likely to play a regulatory role (Figure 7). The AL proteins exhibit a negative effect at the TSS-flanking region and a positive effect at the intragenic region, suggesting that they regulate the H2A.Z deposition in a locus-dependent manner (Figure 7I, Supplemental Figure 22). While the positive effect of AL proteins on H2A.Z deposition can be explained by their interaction with the SWR1 complex components, how the AL proteins negatively regulate the H2A.Z deposition is elusive. Because the AL proteins negatively regulate H2A.Z deposition primarily at genes not occupied by PIE1, we predict that they suppress H2A.Z deposition through an SWR1-independent manner. Interestingly, our AP-MS results also identified the INO80 chromatin-remodeling complex (Figure 2A, Supplemental Figure 9E, Supplemental Dataset 1), which is thought to facilitate nucleosome sliding and the exchange of H2A.Z for H2A (Yang et al., 2020; Shang et al., 2021). Therefore, it is possible that the AL proteins negatively impact the H2A.Z deposition though interacting with the INO80 complex.

Three types of SWI/SNF complexes, BAS, SAS, and MAS, have been found to facilitate chromatin accessibility at different genomic regions relative to the TSS (Guo et al., 2022; Fu et al., 2023). The BAS and MAS complexes primarily mediate chromatin accessibility mainly at the TSS-proximal region, which is enriched with active histone marks including H3K4me3, while the SAS complex primarily mediates chromatin accessibility at the upstream promoter region, which is depleted of H3K4me3 (Guo et al., 2022). Our study revealed that the AL proteins selectively interact with the BAS and MAS complex components but not with the SAS complex components, in line with the observation that the AL proteins bind to the TSS-proximal region enriched with H3K4me3 (Figure 2A, Supplemental Dataset 1). Since the AL proteins were found to be dispensable for the MAS-mediated chromatin accessibility, it is unlikely that the AL proteins function upstream of the BAS and MAS complexes (Figure 8E). Given that the AL proteins interact with the BAS and MAS complexes through the PAL domain and that the PAL domain is crucial for the association of AL proteins with chromatin, it is possible that the interaction of the BAS and MAS with the AL proteins plays a role in the association of AL proteins with chromatin (Supplemental Figure 11). The possibility is supported by the finding that the enrichment of AL2 at its target genomic loci is reduced in the *minu1/2* mutant compared to the wild type. Although the involvement of other AL-interacting chromatin-related proteins in the association of AL proteins with chromatin has not been determined, it is reasonable to infer that multiple AL-interacting chromatin-related proteins have a synergistic effect on the association of AL proteins with chromatin.

The H3K4me3 mark is widely distributed at numerous protein-coding genes in euchromatin regions (Ruthenburg et al., 2007; Zhang et al., 2009). However, the association of H3K4me3 with other chromatin modifications to regulate transcription remain elusive. Our study not only reveals that the AL proteins are genome-wide H3K4me3-binding proteins but also demonstrates that they regulate multiple chromatin modifications, thereby elucidating a complex network that links diverse chromatin modifications. In particular, our study demonstrated that the AL proteins function as subunits of the PRC1 complex and thereby play a major role in regulating H2Aub, while they are unlikely to be crucial for regulating H2A.Z deposition, H3K27me3, and chromatin accessibility. Given that the PAL domain is required for the association of AL proteins with chromatin, the interaction of the PAL domain with chromatin-related proteins is likely to mediate the association of AL proteins with chromatin. Therefore, the AL proteins are likely to be downstream effecters or act in parallel to H2A.Z deposition, H3K27me3, and chromatin accessibility, rather than upstream regulators of them. Overall, our study reveals that the plant-specific AL proteins play a crucial role in mediating the connection between H3K4me3 and other chromatin modifications, providing a foundation for further investigating the interaction network of diverse chromatin modifications in plants.

## Materials and methods

### Plant materials and growth conditions

All the *Arabidopsis* plants used in this study belonged to the Col-0 ecotype. T-DNA insertion mutants, including *al5* (SALK_081524), *al6* (SALK_040877) and *al7* (SALK_032503), were obtained from the Arabidopsis Biological Resource Center (ABRC). Through CRISPR/Cas9-mediated genome editing, we generated *al1*, *al2*, *al3*, and *al4* single mutants. Multiple mutants, including double, triple, quadruple, and quintuple mutants were developed through a combination of genome editing and genetic crossing. Owing to severe developmental defects observed in *al* multiple mutants, the high-order mutants *al1/3/4/5/6/7* and *al1/3/4/5/6/7;2^+/-^* mutants were derived from the segregation of heterozygotes. Arabidopsis seedlings were grown on MS (Murashige and Skoog) medium plates at 22 °C for 16 h in the light and 8 h in the dark. Primers utilized for genotyping are detailed in Supplemental Dataset 7.

The full-length genomic sequences of *AL1*, *AL2*, *AL3*, *AL4*, *AL5*, *AL6*, *AL7*, *SNL2*, *VAL1*, *ZF2*, *GATA1* and *RING1A*, along with their native promoters, were cloned into the *pCAMBIA1305* vector with a 3’-terminal 3×FLAG tag or a 3’-terminal GFP tag. Additionally, the genomic sequences encoding AL2-PALΔ, AL2-PAL-M, AL2-PHDΔ, and AL2-PHD-M proteins were inserted into *pCAMBIA1305* vector with a 3’-terminal 3×FLAG tag. These constructs were transformed into Agrobacterium strain GV3101 and subsequently into Arabidopsis plants via the flower-dipping method, yielding transgenic plants. Transgenic plants were selected based on resistance to hygromycin (30 mg/l) and ampicillin (50 mg/l). The constructs were confirmed by sequencing, with primers used in the cloning listed in Supplemental Dataset 7.

### AP–MS analysis

Eleven-day-old WT and transgenic Arabidopsis seedlings, weighing 6 grams, were harvested and processed into a fine powder using liquid nitrogen. The powder was resuspended in 25 ml of ice-cold plant lysis buffer (50 mM Tris-HCl pH 7.5, 150 mM NaCl, 5 mM MgCl_2_, 10% glycerol, 0.1% NP-40, 5 mM dithiothreitol (DTT), 1 mM phenylmethylsulfonyl fluoride (PMSF) and 0.1% Roche protease inhibitor cocktail tablet) and rotated for 1-2 h at 4°C to ensure complete lysis. After the lysate was centrifuged at 12,000 g for 15 min, the supernatant was filtered through Miracloth (Merck Millipore, 475855) and incubated with anti-FLAG M2 agarose (Sigma-Aldrich, A2220, 1:200 dilution) at 4 °C for 2.5 h. After incubation, the agarose beads were washed five times with cold plant lysis buffer to eliminate non-specific binding proteins. Proteins were eluted using 3×FLAG peptides (Sigma, F4799) and separated by 10% SDS–PAGE gel, followed by silver staining with the ProteoSilver Silver Stain Kit (Sigma, PROT-SIL1).

Mass spectrometry analysis was performed as previously described (Zhang et al., 2012). Briefly, the silver-stained protein bands from the gel were excised, de-stained, and subjected to overnight tryptic digestion at 37°C. The resultant peptides were subsequently eluted through a capillary column and sprayed into a Q Exactive Mass Spectrometer, which was equipped with a nano-ESI ion source (Thermo Fisher Scientific). The mass spectra obtained were searched against the International Protein Index (IPI) database for Arabidopsis. The mass spectrometry data were visualized using heatmaps generated with GraphPad Prism (v8). The diagram of protein-protein interaction as determined by AP-MS was drawn using Cytoscape (v3.6.1) software. The AP-MS datasets for the components of various complexes, including SWI/SNF (Guo et al., 2022), PEAT (Tan et al., 2018; Zheng et al., 2023), SWR1 (Luo et al., 2020), AuPC (Qi et al., 2022), HDAC (Ning et al., 2019; Feng et al., 2021), and INO80 complexes (Shang et al., 2021) were compiled from our previous studies.

### Protein purification and *in vitro* pull-down assay

For protein purification, the full-length and truncated coding sequence of Arabidopsis *AL2*, *AL4*, *RING1A*, *BMI1B*, *REF6*, *BRM*, *MINU2*, *SWI3B*, *SNL2*, *HDA19*, *HDA6*, *PIE1*, *CHR11*, *VAL1*, *NDX*, *ZF2*, and *GATA1* were cloned into *pGEX-6p-1* in fusion with a GST tag, *pSMT3* in fusion with a His tag, and *pET30a* in fusion with MBP and His tags. Primers used for the construction of the recombinant plasmids are listed in Supplemental Dataset 7. The constructs expressing PWWP1, PWWP1-3, and UBP5 are derived from our previous studies (Tan et al., 2018; Zheng et al., 2023). The constructs were validated by Sanger sequencing and transformed into *Escherichia coli* strain BL21 (Rosetta DE3). Positive bacterial strains were grown in LB medium at 37°C until the OD_600_ reached 1.0, after which the fusion proteins were induced with 0.1 mM IPTG at 16°C for 16 h. The cells were harvested by centrifugation, the pellet was suspended in GST-tag lysis buffer (20 mM Tris–HCl at pH 7.4, 500 mM NaCl, 1 mM DTT and 1 mM PMSF), His-tag lysis buffer (20 mM Tris–HCl at pH 7.8, 500 mM NaCl, 20 mM imidazole, 1 mM DTT and 1 mM PMSF), or MBP-tag lysis buffer (20 mM Tris–HCl at pH 7.4, 200 mM NaCl, 1mM EDTA pH 8.0, 1 mM DTT and 1 mM PMSF). The lysate was sonicated for 15 min (3s on, 7s off) and then centrifuged at 18,000 g for 1 h at 4 °C to collect the supernatant, which was then incubated with GST-beads (GE Healthcare, 17-0756-01) or His-beads (Millipore, 70666-4) or MBP-beads (Smart-Lifesciences, SA077025) for 2 hours at 4 °C. The protein-bound beads were washed six times with GST-tag lysis buffer or His-tag lysis buffer to remove the non-specific binding proteins. GST-tag elution buffer (GST-tag lysis buffer containing 20 mM Glutathione) or His-tag elution buffer (His-tag lysis buffer containing 250 mM imidazole) or MBP-tag elution buffer (MBP-tag lysis buffer containing maltose) were added to elute the binding proteins.

For the *in vitro* pull-down assay, a mixture of proteins bearing different tags was prepared and incubated in GST-tag lysis buffer at 4°C for 1 h. Following the incubation, a subset of the mixture was reserved as the input control. The remainder of the sample was then incubated with GST-beads and gently rotated for 1 h at 4°C to facilitate binding. Post-incubation, the GST-beads were washed six times with cold GST-tag lysis buffer to eliminate any non-specific interactions. The bound proteins were eluted by incubating the beads with GST-tag lysis buffer containing 20 mM glutathione for 30 min at 4 °C. Subsequently, both the eluted and reserved input control were resolved on SDS–PAGE gels. The separated proteins were then subjected to immunoblotting using GST antibody (M20007L; Abmart; 1:5,000 dilution) and His antibody (F013; ORIGENE; 1:5,000 dilution) or MBP antibody (AE016; Abclonal; 1:5,000 dilution).

### *In vitro* histone peptide binding assay

For histone peptide binding assays, 1 μg of biotin-labelled histone peptides, either modified or unmodified, were mixed with 1 μg of GST-tagged purified protein in peptide binding buffer (50 mM Tris-HCl at pH 7.5, 150 mM NaCl, 0.05% NP-40, 1 mM DTT and 1 mM PMSF). The mixture was then gently rotated at 4 °C for 4 h to allow for binding. A control group, consisting of purified protein without the addition of histone peptides, was utilized to serve as a negative control. Following the incubation, the mixture was further incubated with 30 μl of Streptavidin MagneSphere paramagnetic particles (PROMEGA, Z548D), with gentle rotation maintained at 4 °C for 1 h to selectively capture the biotinylated peptides. To ensure the removal of any non-specific interactions, the protein-bound beads were washed five times using the same peptide binding buffer. The beads were then subjected to a boiling step to elute the bound protein, which were subsequently analyzed by western blotting using GST-specific antibodies. The histone peptides utilized in this study, including H3 (1–19 aa), H3K4me1 (1–21 aa), H3K4me2 (1–21 aa), H3K4me3 (1–21 aa), were custom synthesized (Scilight-Peptide).

### ChIP-seq and data analysis

ChIP experiments were performed as previously described (Wu et al., 2023). In brief, 4 g of 11-day-old Arabidopsis seedlings grown on MS medium was incubated in the cross-linking buffer (0.4 M sucrose, 10 mM Tris–HCl pH 8.0, 10 mM MgCl_2_, 1 mM PMSF, 1% formaldehyde) for 12 min under vacuum. The cross-linking reaction was quenched by the addition of 0.125 M glycine. The seedlings were then ground into a fine power in liquid nitrogen and resuspended in buffer I (10 mM Tris–HCl pH 8.0, 10 mM MgCl_2_, 0.4 M sucrose, 1 mM DTT, 0.1 mM PMSF, 1% Roche protease inhibitor cocktail tablet) for 45 min at 4°C. The suspension was filtered through Microcloth (Millipore, 475855) and centrifuged at 4,000 g for 20 min. The resulting pellet was washed four times with buffer II (10 mM Tris–HCl pH 8.0, 10 mM MgCl_2_, 0.25 M sucrose, 1% triton X-100, 1 mM DTT, 0.1mM PMSF, 1% Roche protease inhibitor cocktail tablet) until the green color was completely removed. Next, the pellet was resuspended in buffer III (10 mM Tris–HCl pH 8.0, 2mM MgCl_2_, 1.7 M sucrose, 0.15% triton X-100, 1 mM DTT, 0.1mM PMSF, 1% Roche protease inhibitor cocktail tablet), overlaid with an equal volume of buffer III, and centrifuged at 14,000 g at 4°C for 1 h.

The precipitate was resuspended in sonication buffer and subjected to sonication using a Bioruptor sonicator for 28 cycles. Following the sonication, the sample was centrifuged at 14,000 g for 15 min at 4°C, and the supernatant was subsequently diluted 5-fold in dilution buffer (16.7mM Tris-HCl pH 8.0, 167 mM NaCl, 1.2 mM EDTA pH 8.0, 1.1% triton X-100, 1 mM DTT, 0.1mM PMSF, 1% Roche protease inhibitor cocktail tablet). This diluted supernatant was then incubated overnight with a specific antibody at 4°C. Protein A (Thermo, 10001D) or Protein G (Thermo, 10004D) beads were conjugated to the antibody, and these antibody-conjugated beads were washed sequentially with a series of buffers: low salt buffer (20 mM Tris-HCl pH 8.0, 150 mM NaCl, 0.1% SDS, 1% triton X-100, 2 mM EDTA pH 8.0), high salt buffer (20 mM Tris-HCl pH 8.0, 500 mM NaCl, 0.1% SDS, 1% triton X-100, 2 mM EDTA pH 8.0), LiCl buffer (10 mM Tris-HCl pH 8.0, 0.25 M LiCl, 1% sodium deoxycholate, 1 mM EDTA pH 8.0, 1% NP-40), and TE buffer (10 mM Tris-HCl pH 8.0, 1 mM EDTA pH 8.0).

The protein–DNA complex was eluted and then subjected to reverse cross-linking at 65°C overnight. The proteins in the eluate were digested using proteinase K, and the DNA was extracted using a phenol/chloroform/isoamyl (Biorigin; 25:24:1) reagent. The NEXTFLEX® Rapid DNA Seq Kit 2.0 (NOVA-5188-01) was utilized for the preparation of the next-generation sequencing library from the purified DNA. The library preparation process included DNA end repair and adenylation, adapter ligation, bead-based DNA size selection, and PCR amplification, with detailed protocols provided in the accompanying manual. High-throughput sequencing was performed on the Illumina-NovaSeq platform using a paired-end 150 (PE150) sequencing method. For the ChIP-seq of H2A.Z, Dynabeads Protein G were used in conjunction with an H2A.Z antibody, which was produced in rats against a histone peptide as previously described (Deal et al., 2007). For the ChIP-seq of H2Aub, H3K27me3, and GFP-tagged AL proteins (AL2, AL4, and AL6), Dynabeads Protein A were used in conjunction with an H2Aub antibody (8240s; CST; diluted at 1:500), an H3K27me3 antibody (07-449; Millipore; diluted at 1:2,000), and a GFP antibody (Abcam; ab290; diluted 1:600), respectively. For the ChIP-seq of FLAG-tagged AL1, AL2, AL3, AL6, AL2-PAL-M, AL2-PALΔ, AL2-PHD-M, AL2-PHDΔ, and RING1A, Dynabeads Protein G were conjugated with a FLAG antibody (Sigma; F1804; diluted 1:1,000).

For ChIP–seq data analysis, clean reads were aligned to the Arabidopsis genome (TAIR10) using Bowtie2 (v2.3.4), allowing for one mismatch after adapter removal and lowLquality read filtering (Langmead and Salzberg, 2012). PCR duplicates were identified and removed using Picard Tools (v2.23.0) with MarkDuplicates (https://broadinstitute.github.io/picard/). Read counts were normalized to RPKM (reads per kilobase per million mapped reads) based on the number of clean reads aligned to the genome in each library. For GFP or Flag ChIP-seq, enriched peaks were identified using MACS2 (v2.2.7.1) with input reads serving as a negative control (Zhang et al., 2008). For H2Aub, H3K27me3 and H2A.Z ChIP-seq, differentially enriched peaks were identified between WT and mutant samples using SICER2 (v1.0.2) with a filter (FC < 0.8 or > 1.2, FDR < 0.05) (Zang et al., 2009). H3K4me3 ChIP-seq data (Chen et al., 2017), H3K9Ac ChIP-seq data (Wu et al., 2021), H3K36me3 ChIP-seq data (Qi et al., 2022), REF6 and H3K27me3 ChIP-seq data (Cui et al., 2016), PIE1 and H2A.Z ChIP-seq data (Luo et al., 2020), and BRD1, SWI3C, MINU2 and PMS2B ChIP-seq data (Guo et al., 2022) were derived from prior research. The ChIP-seq results presented in this study were based on two independent biological replicates. Visualization of ChIP-seq data was performed using various tools. Meta plots and heatmaps were generated with the plotProfile and plotHeatmap tools from DeepTools (v3.5.1). Venn diagrams were created using the R package ggvenn (v0.1.9). Box plots and scatter plots were crafted with the R package ggplot2 (v3.3.5). The correlation heatmap was produced using the heatmap.2 function from the R package gplots (v3.1.1).

### ATAC-seq and data analysis

ATAC-seq experiments were performed as previously described (Guo et al., 2022). Approximately 0.1 g of 11-day-old Arabidopsis seedlings, cultivated on MS medium, were finely chopped into pieces in cold nuclei extraction buffer (10 mM Tris–HCl pH 8.0, 10 mM MgCl_2_, 0.25 M sucrose, 3% triton X-100, 1% Roche protease inhibitor cocktail tablet). The resulting mixture was filtered through a 40 μm cell sieve into a collection tube and centrifuged at 1,000 g for 10 min at 4°C. The pellet was washed with the same cold nuclei extraction buffer (10 mM Tris–HCl pH 8.0, 10 mM MgCl_2_, 0.25 M sucrose, 3% triton X-100, 1% Roche protease inhibitor cocktail tablet) and then resuspended in nuclei purification buffer (20 mM MOPS pH 7.0, 40 mM NaCl, 90 mM KCl, 2 mM EDTA, 0.5 mM EGTA, 0.5 mM spermidine, 0.2 mM spermine, 1% Roche protease inhibitor cocktail tablet). A sample aliquot was taken for microscopic examination to determine the appropriate volume of the nuclear suspension for subsequent steps. Approximately 10,000 to 50,000 nuclei were used for Tn5 tagmentation and library construction with the TruePrep DNA library Prep Kit V2 for Illumina (Vazyme, TD501). High-throughput sequencing was performed on the Illumina-NovaSeq platform using the PE150 method.

For ATAC–seq data analysis, clean reads were aligned to the Arabidopsis genome (TAIR10) with Bowtie2 (v2.3.4), allowing for one mismatch after adapter trimming and removal of lowLquality reads (Langmead and Salzberg, 2012). PCR duplicates were eliminated using Picard Tools (v2.23.0) with the MarkDuplicates function (https://broadinstitute.github.io/picard/). ATAC-seq peaks were identified using MACS2 (v2.2.7.1) with the following parameters: --nomodel --shift -100 –extsize 200 --bdg --keep-dup all (Zhang et al., 2008). Differentially accessible genomic regions were identified using the DiffBind package (v3.0) in the R program (Stark and Brown, 2011). The study presents ATAC-seq data derived from two independent biological replicates. Visualization of the data was achieved using box plots and scatter plots generated with the R package ggplot2 (v3.3.5).

### Domain prediction, sequence alignment, and phylogenetic analysis

Protein sequences utilized in this study were sourced from the TAIR (https://www.arabidopsis.org) and the NCBI database (https://www.ncbi.nlm.nih.gov). The domains within each protein were predicted using NCBI’s conserved domain Search tools (https://www.ncbi.nlm.nih.gov/Structure/cdd/wrpsb.cgi) and Uniprot (https://www.uniprot.org). Multiple protein sequence alignments were conducted using the DNAMAN software (version 7). A phylogenetic tree for ALFIN-LIKE proteins in Arabidopsis were generated by MEGA (version 7) with the neighbor-joining method.

### Accession numbers

The Arabidopsis sequence data employed in this study are derived from TAIR10. The accession numbers of genes reported in this study are as follows: *AL1* (AT5G05610), *AL2* (AT3G11200), *AL3* (AT3G42790), *AL4* (AT5G26210), *AL5* (AT5G20510), *AL6* (AT2G02470), *AL7* (AT1G14510), *RING1A* (AT5G44280), *BMI1B* (AT1G06770), *RING1B* (AT1G03770), *BMI1A* (AT2G30580), *PIE1* (AT3G12810), *CHR11* (AT3G06400), *ARP6* (AT3G33520), *CHR17* (AT5G18620), *MBD9* (AT3G01460), *RIN1* (AT5G22330), *RIN2* (AT5G67630), *SWC2* (AT2G36740), *GAS41* (AT5G45600), *SWC4* (AT2G47210), *SWC6* (AT5G37055), *TRA1A* (AT2G17930), *TRA1B* (AT4G36080), *REF6* (AT3G48430), *MINU1* (AT3G06010), *MINU2* (AT5G19310), *SWI3A* (AT2G47620), *SWI3B* (AT2G33610), *BSH* (AT3G17590), *SHH2* (AT3G18380), *PMS1A* (AT1G50620), *PMS2A* (AT3G08020), *PMS2B* (AT3G52100), *BRD5* (AT1G58025), *MIS* (AT1G32730), *SSM* (AT1G06500), *LFR* (AT3G22990), *BRM* (AT2G46020), *SWI3C* (AT1G21700), *BRD1* (AT1G20670), *BRD2* (AT1G76380), *BRD13* (AT5G55040), *BRIP2* (AT5G17510), *BRIP1* (AT3G03460), *MSI1* (AT5G58230), *RXT3* (AT5G08450), *SNL3* (AT1G24190), *SNL4* (AT1G70060), *SNL5* (AT1G59890), *SNL6* (AT1G10450), *SNL2* (AT5G15020), *HDA19* (AT4G38130), *HDA6* (AT5G6310), *FLD* (AT3G10390), *SDG26* (AT1G76710), *EFL4* (AT1G17455), *EFL2* (AT1G72630), *UBP5* (AT2G40930), *PWWP1* (AT3G03140), *ARID2* (AT2G17410), *ARID3* (AT1G20910), *ARID4* (AT1G76510), *EPCR1* (AT4G32620), *PWWP2* (AT1G51745), *PWWP3* (AT3G21295), *TRB1* (AT1G49950), *HAM1* (AT5G64610), *FUG1* (AT1G17220), *WDR5A* (AT3G49660), *IES2B* (AT3G06660), *TRO* (AT1G51450), *RBL* (AT3G21060), *YY1* (AT4G06634), *ARP9* (AT5G43500), *ARP5* (AT3G12380), *UCH1* (AT5G16310), *UCH2* (AT1G65650), *INO80* (AT3G57300), *EEN* (AT4G38495), *JMJ24* (AT1G09060), *NFRKB1* (AT3G45830), *NFRKB2* (AT5G13950), *INB1* (AT4G18400), *INB3* (AT3G51500), *INB2B* (AT5G57910), *ATX2* (AT1G05830), *GATA1* (AT3G24050), *ZF2* (AT3G19580), *VAL1* (AT2G30470), *NDX* (AT4G03090).

## Data availability

Raw ChIP-seq data have been deposited in the Gene Expression Omnibus (GEO) database with the accession code GSE270449. The access token is ‘wpmrgmucbretryp’. Raw ATAC-seq data have been deposited in the GEO database with the accession code GSE270450. The access token is ‘kjazwegcdzgphkz’.

## Acknowledgements

We thank Dr. Ligeng Ma for providing the *ring1a/1b* double mutant seeds. This work was supported by the National Natural Science Foundation of China (32025003 to XJH).

## Author contributions

XMS and XJH were responsible for the project conception, experimental design, and manuscript preparation. XMS carried out the experiments. DYY performed the bioinformatic analysis. LL and SC performed the mass spectrometry analysis. MY and YZ provided assistance in the generation of *BMI1A* and *RING1A* transgenic plants and analysis of RING1A ChIP-seq data.

## Competing interests

The authors declare no competing interests.

## Supplemental Information

Supplemental Figure 1. Genotypic analysis of *al1*, *al2*, *al3*, *al4*, *al5*, *al6*, and *al7* mutants.

Supplemental Figure 2. Morphological phenotypes of the *al* single mutants.

Supplemental Figure 3. Morphological phenotypes of the *al* double mutants.

Supplemental Figure 4. Phenotypic complementation test of the *al2/4* double mutant.

Supplemental Figure 5. Morphological phenotypes of the *al* triple mutants.

Supplemental Figure 6. Morphological phenotypes of the *al* quadruple mutants.

Supplemental Figure 7. Statistical analysis of morphological phenotypes in the *al* quintuple mutants.

Supplemental Figure 8. Statistical evaluation of the impact of individual *al* mutations on morphological change in *al* mutant plants.

Supplemental Figure 9. Determination of the interactions of AL proteins with PEAT, AuPC, SWI/SNF, HDAC, and INO80 complex components by AP-MS.

Supplemental Figure 10. Determination of the interactions of AL proteins with SWR1 complex components, PRC1 complex components, and transcription factors by AP-MS.

Supplemental Figure 11. Determination of the interactions of AL proteins with transcription factors and SWI/SNF complex components by pull-down assays.

Supplemental Figure 12. Sequence analysis of Arabidopsis AL proteins.

Supplemental Figure 13. Phylogenetic analysis of AL1 orthologs in plants.

Supplemental Figure 14. Determination of the interaction between AL2 and AL4 by pull-down assays.

Supplemental Figure 15. Determination of the interactions of AL proteins with HDAC complex components and AuPC complex components by pull-down assays.

Supplemental Figure 16. Determination of the interactions between AL proteins and components of PEAT or PRC1 complexes and between RING1A and BMI1B by pull-down assays.

Supplemental Figure 17. The W206A and D210N mutations within the PHD domain disrupt the binding of AL2 to methylated H3K4 peptides *in vitro*.

Supplemental Figure 18. Co-occupancy of different AL proteins on chromatin at the whole-genome level.

Supplemental Figure 19. Determination of the impact of the deletion or mutation of the PHD domain on the association of AL2 with chromatin at the whole-genome level.

Supplemental Figure 20. Comparison of the effect of *al2/4/5/6/7* and *ring1a/1b* on H2Aub.

Supplemental Figure 21. Analysis of the correlation between H2Aub and H3K4me3 at the whole-genome level.

Supplemental Figure 22. Analysis of the genomic distribution of H2A.Z level changes in the *al2/4/5/6/7* mutant relative to the wild type.

Supplemental Dataset 1. Full list of Arabidopsis proteins identified by affinity purification followed by mass spectrometry analysis.

Supplemental Dataset 2. Peaks identified by ChIP-seq.

Supplemental Dataset 3. Peaks with reduced and increased H2Aub levels in *al2/4/5/6/7*, *ring1a/1b* and *minu1/2* mutants relative to the wild type.

Supplemental Dataset 4. Peaks with reduced and increased H3K27me3 levels in *al2/4/5/6/7* mutant relative to the wild type.

Supplemental Dataset 5. Peaks with reduced and increased H2A.Z levels in *al2/4/5/6/7* mutant relative to the wild type.

Supplemental Dataset 6. Differentially accessible regions identified by ATAC-seq.

Supplemental Dataset 7. List of primers used in this study.

## Notes

### Competing Interest Statement

The authors have declared no competing interest.

## References

Bannister, A.J., and Kouzarides, T. (2011). Regulation of chromatin by histone modifications. Cell Res. 21, 381–395.

Barbour, H., Daou, S., Hendzel, M., and Affar, E.B. (2020). Polycomb group-mediated histone H2A monoubiquitination in epigenome regulation and nuclear processes. Nat. Commun. 11, 5947.

Berr, A., Shafiq, S., Pinon, V., Dong, A., and Shen, W.H. (2015). The trxG family histone methyltransferase SET DOMAIN GROUP 26 promotes flowering via a distinctive genetic pathway. Plant J. 81, 316–328.

Bratzel, F., Lopez-Torrejon, G., Koch, M., Del Pozo, J.C., and Calonje, M. (2010). Keeping cell identity in Arabidopsis requires PRC1 RING-finger homologs that catalyze H2A monoubiquitination. Curr. Biol. 20, 1853–1859.

Cao, R., Tsukada, Y., and Zhang, Y. (2005). Role of Bmi-1 and Ring1A in H2A ubiquitylation and Hox gene silencing. Mol. Cell 20, 845–854.

Chen, D., Molitor, A., Liu, C., and Shen, W.H. (2010). The Arabidopsis PRC1-like ring-finger proteins are necessary for repression of embryonic traits during vegetative growth. Cell Res. 20, 1332–1344.

Chen, L., Luo, J., Cui, Z., Xue, M., Wang, L., Zhang, X., Pawlowski, W.P., and He, Y. (2017). ATX3, ATX4, and ATX5 Encode Putative H3K4 Methyltransferases and Are Critical for Plant Development. Plant Physiol. 174, 1795-1806.

Chiang, Y.H., Zubo, Y.O., Tapken, W., Kim, H.J., Lavanway, A.M., Howard, L., Pilon, M., Kieber, J.J., and Schaller, G.E. (2012). Functional characterization of the GATA transcription factors GNC and CGA1 reveals their key role in chloroplast development, growth, and division in Arabidopsis. Plant Physiol. 160, 332–348.

Clapier, C.R., and Cairns, B.R. (2009). The biology of chromatin remodeling complexes. Annu. Rev. Biochem. 78, 273–304.

Cui, X., Lu, F., Qiu, Q., Zhou, B., Gu, L., Zhang, S., Kang, Y., Cui, X., Ma, X., Yao, Q., Ma, J., Zhang, X., and Cao, X. (2016). REF6 recognizes a specific DNA sequence to demethylate H3K27me3 and regulate organ boundary formation in Arabidopsis. Nat. Genet. 48, 694-699.

Dambacher, S., Hahn, M., and Schotta, G. (2010). Epigenetic regulation of development by histone lysine methylation. Heredity 105, 24–37.

Deal, R.B., Topp, C.N., McKinney, E.C., and Meagher, R.B. (2007). Repression of flowering in Arabidopsis requires activation of FLOWERING LOCUS C expression by the histone variant H2A.Z. Plant Cell 19, 74–83.

Di Lorenzo, A., and Bedford, M.T. (2011). Histone arginine methylation. FEBS Lett. 585, 2024–2031.

Diego-Martin, B., Pérez-Alemany, J., Candela-Ferre, J., Corbalán-Acedo, A., Pereyra, J., Alabadí, D., Jami-Alahmadi, Y., Wohlschlegel, J., and Gallego-Bartolomé, J. (2022). The TRIPLE PHD FINGERS proteins are required for SWI/SNF complex-mediated+ 1 nucleosome positioning and transcription start site determination in Arabidopsis. Nucleic Acids Res. 50, 10399-10417.

Feng, C., Cai, X.W., Su, Y.N., Li, L., Chen, S., and He, X.J. (2021). Arabidopsis RPD3-like histone deacetylases form multiple complexes involved in stress response. J Genet Genomics 48, 369–383.

Fu, W., Yu, Y., Shu, J., Yu, Z., Zhong, Y., Zhu, T., Zhang, Z., Liang, Z., Cui, Y., Chen, C., and Li, C. (2023). Organization, genomic targeting, and assembly of three distinct SWI/SNF chromatin remodeling complexes in Arabidopsis. Plant Cell. 35, 2464–2483.

Godwin, J., Govindasamy, M., Nedounsejian, K., March, E., Halton, R., Bourbousse, C., Wolff, L., Fort, A., Krzyszton, M., Lopez Corrales, J., Swiezewski, S., Barneche, F., Schubert, D., and Farrona, S. (2024). The UBP5 histone H2A deubiquitinase counteracts PRCs-mediated repression to regulate Arabidopsis development. Nat. Commun. 15, 667.

Gómez-Zambrano, Á., Crevillén, P., Franco-Zorrilla, J.M., López, J.A., Moreno-Romero, J., Roszak, P., Santos-González, J., Jurado, S., Vázquez, J., and Köhler, C. (2018). Arabidopsis SWC4 binds DNA and recruits the SWR1 complex to modulate histone H2A. Z deposition at key regulatory genes. Mol. plant 11, 815–832.

Grossniklaus, U., and Paro, R. (2014). Transcriptional silencing by polycomb-group proteins. Cold Spring Harb. Perspect Biol. 6, a019331.

Guo, J., Cai, G., Li, Y.Q., Zhang, Y.X., Su, Y.N., Yuan, D.Y., Zhang, Z.C., Liu, Z.Z., Cai, X.W., Guo, J., Li, L., Chen, S., and He, X.J. (2022). Comprehensive characterization of three classes of Arabidopsis SWI/SNF chromatin remodelling complexes. Nat. Plants 8, 1423-1439.

He, Y., Michaels, S.D., and Amasino, R.M. (2003). Regulation of Flowering Time by Histone Acetylation in Arabidopsis. Science 302, 1751–1754.

Hohenstatt, M.L., Mikulski, P., Komarynets, O., Klose, C., Kycia, I., Jeltsch, A., Farrona, S., and Schubert, D. (2018). PWWP-DOMAIN INTERACTOR OF POLYCOMBS1 Interacts with Polycomb-Group Proteins and Histones and Regulates Arabidopsis Flowering and Development. Plant Cell 30, 117–133.

Jansen, A., and Verstrepen, K.J. (2011). Nucleosome positioning in Saccharomyces cerevisiae. Microbiol Mol. Biol. Rev. 75, 301–320.

Jin, R., Yang, H., Muhammad, T., Li, X., Tuerdiyusufu, D., Wang, B., and Wang, J. (2024). Involvement of Alfin-Like Transcription Factors in Plant Development and Stress Response. Genes 15, 184.

Kapoor, P., and Shen, X. (2014). Mechanisms of nuclear actin in chromatin-remodeling complexes. Trends Cell Biol. 24, 238–246.

Karanyi, Z., Mosolygo, L.A., Fero, O., Horvath, A., Boros-Olah, B., Nagy, E., Hetey, S., Holb, I., Szaker, H.M., Miskei, M., Csorba, T., and Szekvolgyi, L. (2022). NODULIN HOMEOBOX is required for heterochromatin homeostasis in Arabidopsis. Nat. Commun. 13, 5058.

Kayum, M.A., Park, J.I., Ahmed, N.U., Jung, H.J., Saha, G., Kang, J.G., and Nou, I.S. (2015). Characterization and stress-induced expression analysis of Alfin-like transcription factors in Brassica rapa. Mol. Genet. Genomics 290, 1299–1311.

Lachner, M., and Jenuwein, T. (2002). The many faces of histone lysine methylation. Curr. Opin. Cell Biol. 14, 286–298.

Langmead, B., and Salzberg, S.L. (2012). Fast gapped-read alignment with Bowtie 2. Nat. Methods 9, 357–359.

Lee, W.Y., Lee, D., Chung, W.I., and Kwon, C.S. (2009). Arabidopsis ING and Alfin1Llike protein families localize to the nucleus and bind to H3K4me3/2 via plant homeodomain fingers. Plant J. 58, 511–524.

Li, G., and Reinberg, D. (2011). Chromatin higher-order structures and gene regulation. Curr. Opin. Genet. Dev. 21, 175–186.

Li, G., Liu, S., Wang, J., He, J., and Xu, L. (2014). ISWI proteins participate in the genome-wide nucleosome distribution in Arabidopsis. Plant J. 78, 706–714.

Li, J., Wang, Z., Hu, Y., Cao, Y., and Ma, L. (2017). Polycomb group proteins RING1A and RING1B regulate the vegetative phase transition in Arabidopsis. Front. plant Sci. 8, 867.

Liang, X., Lei, M., Li, F., Yang, X., Zhou, M., Li, B., Cao, Y., Gong, S., Liu, K., Liu, J., Qi, C., and Liu, Y. (2018). Family-Wide Characterization of Histone Binding Abilities of PHD Domains of AL Proteins in Arabidopsis thaliana. Protein J. 37, 531–538.

Liang, Z., Yuan, L., Xiong, X., Hao, Y., Song, X., Zhu, T., Yu, Y., Fu, W., Lei, Y., Xu, J., Liu, J., Li, J.F., and Li, C. (2022). The transcriptional repressors VAL1 and VAL2 mediate genome-wide recruitment of the CHD3 chromatin remodeler PICKLE in Arabidopsis. Plant Cell 34, 3915–3935.

Liu, R., Li, X., Chen, W., and Du, J. (2018). Structure and mechanism of plant histone mark readers. Sci. China Life Sci. 61, 170–177.

Liu, C., Lu, F., Cui, X., and Cao, X. (2010). Histone methylation in higher plants. Annu. Rev. Plant Biol. 61, 395–420.

Liu, S., Trejo-Arellano, M.S., Qiu, Y., Eklund, D.M., Kohler, C., and Hennig, L. (2021). H2A ubiquitination is essential for Polycomb Repressive Complex 1-mediated gene regulation in Marchantia polymorpha. Genome Biol. 22, 253.

Lu, F., Cui, X., Zhang, S., Jenuwein, T., and Cao, X. (2011). Arabidopsis REF6 is a histone H3 lysine 27 demethylase. Nat. Genet. 43, 715–719.

Luger, K., Mäder, A.W., Richmond, R.K., Sargent, D.F., and Richmond, T.J. (1997). Crystal structure of the nucleosome core particle at 2.8 Å resolution. Nature 389, 251–260.

Luo, Y.X., Hou, X.M., Zhang, C.J., Tan, L.M., Shao, C.R., Lin, R.N., Su, Y.N., Cai, X.W., Li, L., and Chen, S. (2020). A plantLspecific SWR1 chromatinLremodeling complex couples histone H2A. Z deposition with nucleosome sliding. EMBO J. 39, e102008.

March-Dīaz, R., and Reyes, J.C. (2009). The beauty of being a variant: H2A.Z and the SWR1 complex in plants. Mol. Plant 2, 565-577.

Mehdi, S., Derkacheva, M., Ramstrom, M., Kralemann, L., Bergquist, J., and Hennig, L. (2016). The WD40 Domain Protein MSI1 Functions in a Histone Deacetylase Complex to Fine-Tune Abscisic Acid Signaling. Plant Cell 28, 42–54.

Mizuguchi, Gaku, Shen, Xuetong, Landry, Joe, Wu, Wei-Hua, Sen, and Subhojit. (2004). ATP-Driven Exchange of Histone H2AZ Variant Catalyzed bySWR1 Chromatin Remodeling Complex. Science 303, 343–348.

Molitor, A., and Shen, W.H. (2013). The polycomb complex PRC1: composition and function in plants. J Genet. Genomics 40, 231–238.

Molitor, A.M., Bu, Z., Yu, Y., Shen, W.H., and Goodrich, J. (2014). Arabidopsis AL PHD-PRC1 Complexes Promote Seed Germination through H3K4me3-to-H3K27me3 Chromatin State Switch in Repression of Seed Developmental Genes. PLoS Genet. 10, e1004091.

Ning, Y.Q., Chen, Q., Lin, R.N., Li, Y.Q., Li, L., Chen, S., and He, X.J. (2019). The HDA19 histone deacetylase complex is involved in the regulation of flowering time in a photoperiod-dependent manner. Plant J. 98, 448-464.

Noh, Y.S., and Amasino, R.M. (2003). PIE1, an ISWI family gene, is required for FLC activation and floral repression in Arabidopsis. Plant Cell 15, 1671–1682.

Papamichos-Chronakis, M., Watanabe, S., Rando, O.J., and Peterson, C.L. (2011). Global regulation of H2A.Z localization by the INO80 chromatin-remodeling enzyme is essential for genome integrity. Cell 144, 200-213.

Peng, L., Wang, L., Zhang, Y., Dong, A., Shen, W.H., and Huang, Y. (2018). Structural Analysis of the Arabidopsis AL2-PAL and PRC1 Complex Provides Mechanistic Insight into Active-to-Repressive Chromatin State Switch. J. Mol. Biol. 430, 4245–4259.

Perrella, G., Lopez-Vernaza, M.A., Carr, C., Sani, E., Gossele, V., Verduyn, C., Kellermeier, F., Hannah, M.A., and Amtmann, A. (2013). Histone deacetylase complex1 expression level titrates plant growth and abscisic acid sensitivity in Arabidopsis. Plant Cell 25, 3491–3505.

Peterson, C.L., and Laniel, M.-A. (2004). Histones and histone modifications. Curr. Biol. 14, R546–R551.

Portela, A., and Esteller, M. (2010). Epigenetic modifications and human disease. Nat. Biotechnol. 28, 1057–1068.

Potok, M.E., Wang, Y., Xu, L., Zhong, Z., Liu, W., Feng, S., Naranbaatar, B., Rayatpisheh, S., Wang, Z., Wohlschlegel, J.A., Ausin, I., and Jacobsen, S.E. (2019). Arabidopsis SWR1-associated protein methyl-CpG-binding domain 9 is required for histone H2A.Z deposition. Nat. Commun. 10, 3352.

Qi, P.L., Zhou, H.R., Zhao, Q.Q., Feng, C., Ning, Y.Q., Su, Y.N., Cai, X.W., Yuan, D.Y., Zhang, Z.C., Su, X.M., Chen, S.S., Li, L., Chen, S., and He, X.J. (2022). Characterization of an autonomous pathway complex that promotes flowering in Arabidopsis. Nucleic Acids Res. 50, 7380-7395.

Rosana March-Díaz, M.G.-D., Francisco J. Florencio,, and Reyes, J.C. (2007). SEF, a New Protein Required for Flowering Repression in Arabidopsis, Interacts with PIE1 and ARP6. Plant Physiol. 143, 893-901.

Roudier, F., Ahmed, I., Bérard, C., Sarazin, A., Mary-Huard, T., Cortijo, S., Bouyer, D., Caillieux, E., Duvernois-Berthet, E., and Al-Shikhley, L. (2014). Integrative epigenomic mapping defines four main chromatin states in Arabidopsis. EMBO J. 30, 1928–1938.

Ruthenburg, A.J., Allis, C.D., and Wysocka, J. (2007). Methylation of lysine 4 on histone H3: intricacy of writing and reading a single epigenetic mark. Mol. Cell 25, 15–30.

Shang, J.Y., Lu, Y.J., Cai, X.W., Su, Y.N., Feng, C., Li, L., Chen, S., and He, X.J. (2021). COMPASS functions as a module of the INO80 chromatin remodeling complex to mediate histone H3K4 methylation in Arabidopsis. Plant Cell 33, 3250-3271.

Shang, J.Y., and He, X.J. (2022). Chromatin-remodeling complexes: Conserved and plant-specific subunits in Arabidopsis. J. Integr. Plant Biol. 64, 499–515.

Sijacic, P., Holder, D.H., Bajic, M., and Deal, R.B. (2019). Methyl-CpG-binding domain 9 (MBD9) is required for H2A.Z incorporation into chromatin at a subset of H2A.Z-enriched regions in the Arabidopsis genome. PLoS Genet. 15, e1008326.

Spedaletti, V., Polticelli, F., Capodaglio, V., Schininà, M.E., Stano, P., Federico, R., and Tavladoraki, P. (2008). Characterization of a lysine-specific histone demethylase from Arabidopsis thaliana. Biochemistry 47, 4936–4947.

Stark, R., and Brown, G. (2011). DiffBind differential binding analysis of ChIP-Seq peak data.

Suganuma, T., and Workman, J.L. (2011). Signals and combinatorial functions of histone modifications. Annu. Rev. Biochem. 80, 473–499.

Sureshkumar, S., Bandaranayake, C., Lv, J., Dent, C.I., Bhagat, P.K., Mukherjee, S., Sarwade, R., Atri, C., York, H.M., Tamizhselvan, P., Shamaya, N., Folini, G., Bergey, B.G., Yadav, A.S., Kumar, S., Grummisch, O.S., Saini, P., Yadav, R.K., Arumugam, S., Rosonina, E., Sadanandom, A., Liu, H., and Balasubramanian, S. (2024). SUMO protease FUG1, histone reader AL3 and chromodomain protein LHP1 are integral to repeat expansion-induced gene silencing in Arabidopsis thaliana. Nat. Plants 10, 749-759.

Tan, L.M., Zhang, C.J., Hou, X.M., Shao, C.R., Lu, Y.J., Zhou, J.X., Li, Y.Q., Li, L., Chen, S., and He, X.J. (2018). The PEAT protein complexes are required for histone deacetylation and heterochromatin silencing. EMBO J. 37, e98770.

Wang, H., Wang, L., Erdjument-Bromage, H., Vidal, M., Tempst, P., Jones, R., and Zhang, Y. (2004). Role of histone H2A ubiquitination in Polycomb silencing. Nature 431, 873–878.

Wang, P., Lu, S., Li, W., Ma, Z., Mao, J., and Chen, B. (2023). Genome-wide characterization of Alfin-like (AL) genes in apple and functional identification of MdAL4 in response to drought stress. Plant Cell Rep. 42, 395–408.

Wang, Z., Cao, H., Sun, Y., Li, X., Chen, F., Carles, A., Li, Y., Ding, M., Zhang, C., Deng, X., Soppe, W.J., and Liu, Y.X. (2013). Arabidopsis paired amphipathic helix proteins SNL1 and SNL2 redundantly regulate primary seed dormancy via abscisic acid-ethylene antagonism mediated by histone deacetylation. Plant Cell 25, 149–166.

Weber, C.M., and Henikoff, S. (2014). Histone variants: dynamic punctuation in transcription. Genes Dev. 28, 672–682.

Wei, W., Zhang, Y.Q., Tao, J.J., Chen, H.W., Li, Q.T., Zhang, W.K., Ma, B., Lin, Q., Zhang, J.S., and Chen, S.Y. (2015). The Alfin-like homeodomain finger protein AL5 suppresses multiple negative factors to confer abiotic stress tolerance in Arabidopsis. Plant J. 81, 871-883.

Willige, B.C., Zander, M., Yoo, C.Y., Phan, A., and Chory, J. (2021). PHYTOCHROME-INTERACTING FACTORs trigger environmentally responsive chromatin dynamics in plants. Nat. Genet. 53, 955–961.

Wu, C.J., Liu, Z.Z., Wei, L., Zhou, J.X., Cai, X.W., Su, Y.N., Li, L., Chen, S., and He, X.J. (2021). Three functionally redundant plant-specific paralogs are core subunits of the SAGA histone acetyltransferase complex in Arabidopsis. Mol. Plant 14, 1071-1087.

Wu, C.J., Yuan, D.Y., Liu, Z.Z., Xu, X., Wei, L., Cai, X.W., Su, Y.N., Li, L., Chen, S., and He, X.J. (2023). Conserved and plant-specific histone acetyltransferase complexes cooperate to regulate gene transcription and plant development. Nat. Plants 9, 442-459.

Xue, M., Zhang, H., Zhao, F., Zhao, T., Li, H., and Jiang, D. (2021). The INO80 chromatin remodeling complex promotes thermomorphogenesis by connecting H2A. Z eviction and active transcription in Arabidopsis. Mol. Plant 14, 1799–1813.

Yan, C., Yang, N., Li, R., Wang, X., Xu, Y., Zhang, C., Wang, X., and Wang, Y. (2022). AlfinLlike transcription factor VqAL4 regulates a stilbene synthase to enhance powdery mildew resistance in grapevine. Mol. Plant Pathol. 24, 123–141.

Yan, W., Chen, D., Smaczniak, C., Engelhorn, J., Liu, H., Yang, W., Graf, A., Carles, C.C., Zhou, D.X., and Kaufmann, K. (2018). Dynamic and spatial restriction of Polycomb activity by plant histone demethylases. Nat. Plants 4, 681–689.

Yang, C., Bratzel, F., Hohmann, N., Koch, M., Turck, F., and Calonje, M. (2013). VAL- and AtBMI1-mediated H2Aub initiate the switch from embryonic to postgerminative growth in Arabidopsis. Curr. Biol. 23, 1324–1329.

Yang, C., Yin, L., Xie, F., Ma, M., Huang, S., Zeng, Y., Shen, W.-H., Dong, A., and Li, L. (2020). AtINO80 represses photomorphogenesis by modulating nucleosome density and H2A. Z incorporation in light-related genes. Proc. Natl. Acad. Sci. USA 117, 33679–33688.

Yin, X., Romero-Campero, F.J., de Los Reyes, P., Yan, P., Yang, J., Tian, G., Yang, X., Mo, X., Zhao, S., Calonje, M., and Zhou, Y. (2021). H2AK121ub in Arabidopsis associates with a less accessible chromatin state at transcriptional regulation hotspots. Nat. Commun. 12, 315.

Yin, X., Romero-Campero, F.J., Yang, M., Baile, F., Cao, Y., Shu, J., Luo, L., Wang, D., Sun, S., Yan, P., Gong, Z., Mo, X., Qin, G., Calonje, M., and Zhou, Y. (2023). Binding by the Polycomb complex component BMI1 and H2A monoubiquitination shape local and long-range interactions in the Arabidopsis genome. Plant Cell 35, 2484-2503.

Zang, C., Schones, D.E., Zeng, C., Cui, K., Zhao, K., and Peng, W. (2009). A clustering approach for identification of enriched domains from histone modification ChIP-Seq data. Bioinformatics 25, 1952–1958.

Zhang, C., Cao, L., Rong, L., An, Z., Zhou, W., Ma, J., Shen, W.H., Zhu, Y., and Dong, A. (2015). The chromatin remodeling factor AtINO80 plays crucial roles in genome stability maintenance and in plant development. Plant J. 82, 655–668.

Zhang, C.J., Ning, Y.Q., Zhang, S.W., Chen, Q., Shao, C.R., Guo, Y.W., Zhou, J.X., Li, L., Chen, S., and He, X.J. (2012). IDN2 and its paralogs form a complex required for RNA–directed DNA methylation. PLoS Genet. 8, e1002693.

Zhang, D., Gao, Z., Zhang, H., Yang, Y., Yang, X., Zhao, X., Guo, H., Nagalakshmi, U., Li, D., Dinesh-Kumar, S.P., and Zhang, Y. (2023). The MAPK-Alfin-like 7 module negatively regulates ROS scavenging genes to promote NLR-mediated immunity. Proc. Natl Acad. Sci. USA 120, e2214750120.

Zhang, X., Bernatavichute, Y.V., Cokus, S., Pellegrini, M., and Jacobsen, S.E. (2009). Genome-wide analysis of mono-, di-and trimethylation of histone H3 lysine 4 in Arabidopsis thaliana. Genome biol. 10, 1–14.

Zhang, Y., Liu, T., Meyer, C.A., Eeckhoute, J., Johnson, D.S., Bernstein, B.E., Nusbaum, C., Myers, R.M., Brown, M., Li, W., and Liu, X.S. (2008). Model-based analysis of ChIP-Seq (MACS). Genome Biol. 9, R137.

Zheng, S.Y., Guan, B.B., Yuan, D.Y., Zhao, Q.Q., Ge, W., Tan, L.M., Chen, S.S., Li, L., Chen, S., Xu, R.M., and He, X.J. (2023). Dual roles of the Arabidopsis PEAT complex in histone H2A deubiquitination and H4K5 acetylation. Mol. Plant 16, 1847-1865.

Zhou, W., Wu, J., Zheng, Q., Jiang, Y., Zhang, M., and Zhu, S. (2016). Genome-wide identification and comparative analysis of Alfin-like transcription factors in maize. Genes Genom. 39, 261–275.

Zhou, Y., Romero-Campero, F.J., Gomez-Zambrano, A., Turck, F., and Calonje, M. (2017). H2A monoubiquitination in Arabidopsis thaliana is generally independent of LHP1 and PRC2 activity. Genome Biol. 18, 69.

Zhou, Y., Wang, Y., Krause, K., Yang, T., Dongus, J.A., Zhang, Y., and Turck, F. (2018). Telobox motifs recruit CLF/SWN-PRC2 for H3K27me3 deposition via TRB factors in Arabidopsis. Nat. Genet. 50, 638–644.

Zhu, D., Wen, Y., Yao, W., Zheng, H., Zhou, S., Zhang, Q., Qu, L.J., Chen, X., Wu, Z. (2023). Distinct chromatin signatures in the Arabidopsis male gametophyte. Nat. Genet. 55, 706–720.

