## Supplemental Figures for "The Arabidopsis histone H3K4me3-binding ALFIN-like proteins mediate histone H2A ubiquitination and coordinate diverse chromatin modifications"

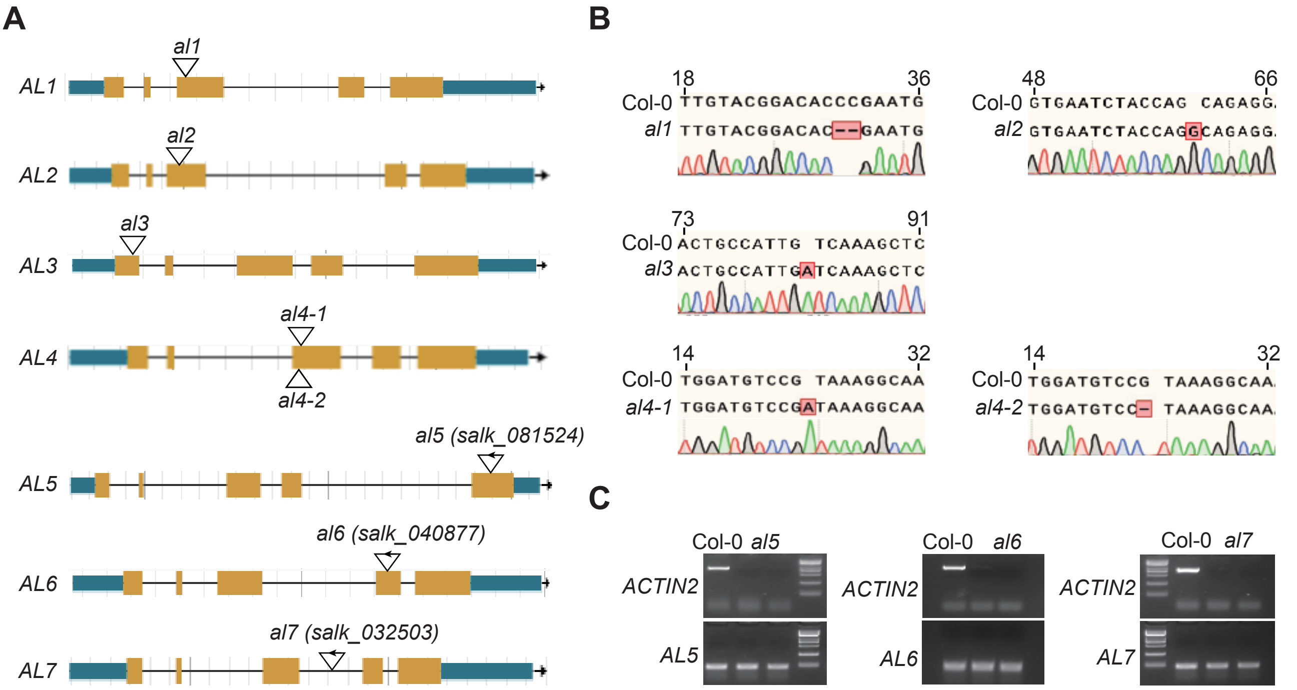


**Supplemental Figure 1. Genotypic analysis of *al1*, *al2*, *al3*, *al4*, *al5*, *al6*, and *al7* mutants.** **(A)** Schematics diagrams of mutations in *AL1*, *AL2*, *AL3*, *AL4*, *AL5*, *AL6* and *AL7* genes. **(B)** Validation of the *al1*, *al2*, *al3*, and *al4* mutations by Sanger sequencing. These mutations were generated by CRISPR/Cas9-mediated genome eding. The mutation sites are highlighted by red boxes. **(C)** Validation of *al5*, *al6*, and *al7* mutations by RT-PCR. The expression levels of *AL5*, *AL6*, and *AL7* were determined by RT-PCR in the wild type and corresponding T-DNA mutants. The expression level of *ACTIN2* is shown as a control.

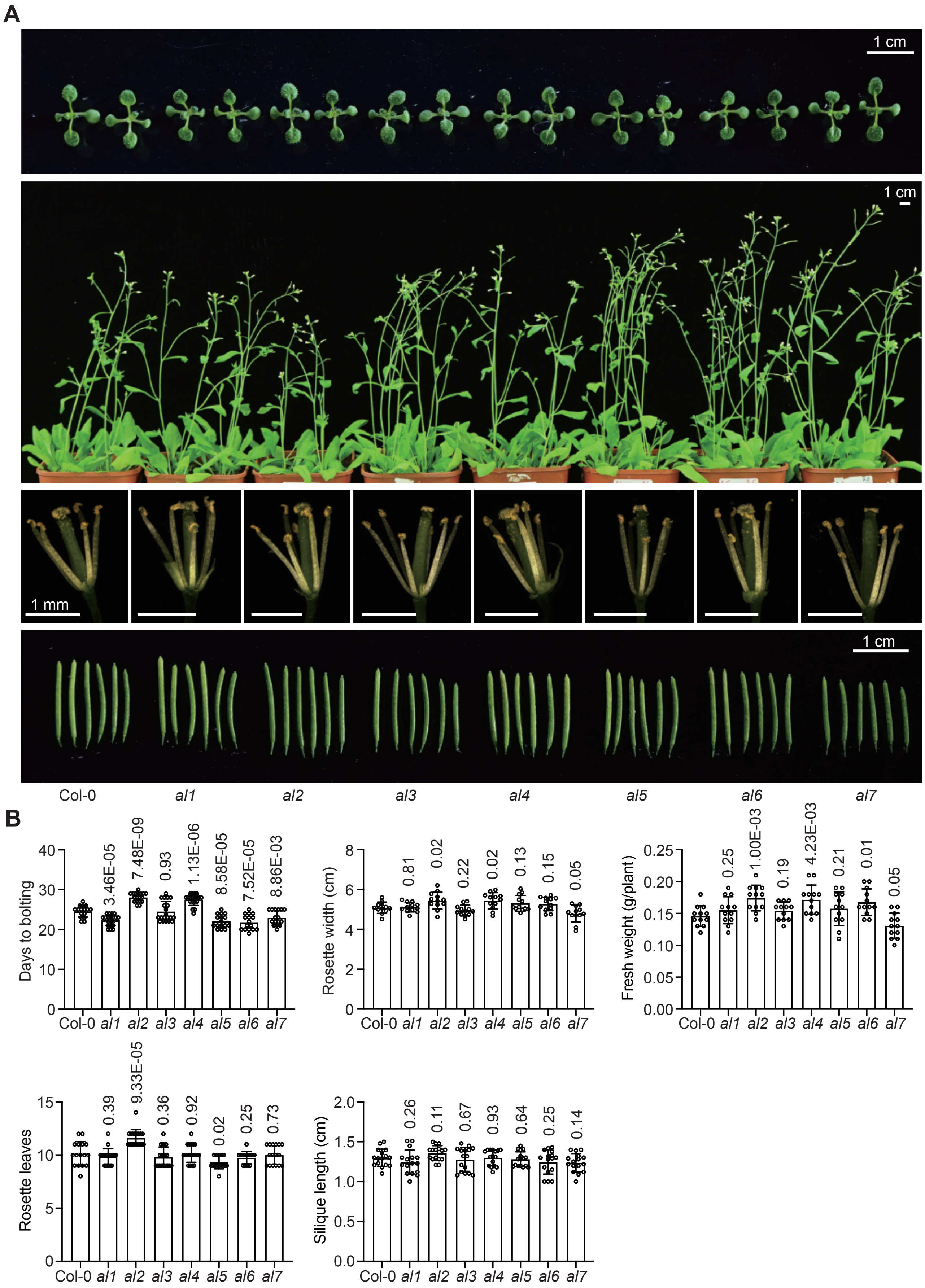


**Supplemental Figure 2. Morphological phenotypes of the *al* single mutants. (A)** From top to bottom, 11-day-old plants (top), 30-day-old plants, stamens and pistils under a microscope, and mature siliques (bottom) are shown for wild-type and *al* single mutants. **(B)** The statistical results of rosette width (20-day-old plants, *n* = 12), fresh weight (20-old-day plants, *n* = 12), days to bolting (*n* >= 16), the number of rosette leaves (*n* >= 16), and siliques length after maturing (*n* = 16). Values are mean ± SD. *P* values determined by two-tailed Student's t-test indicate the difference between the mutants and the wild-type control.

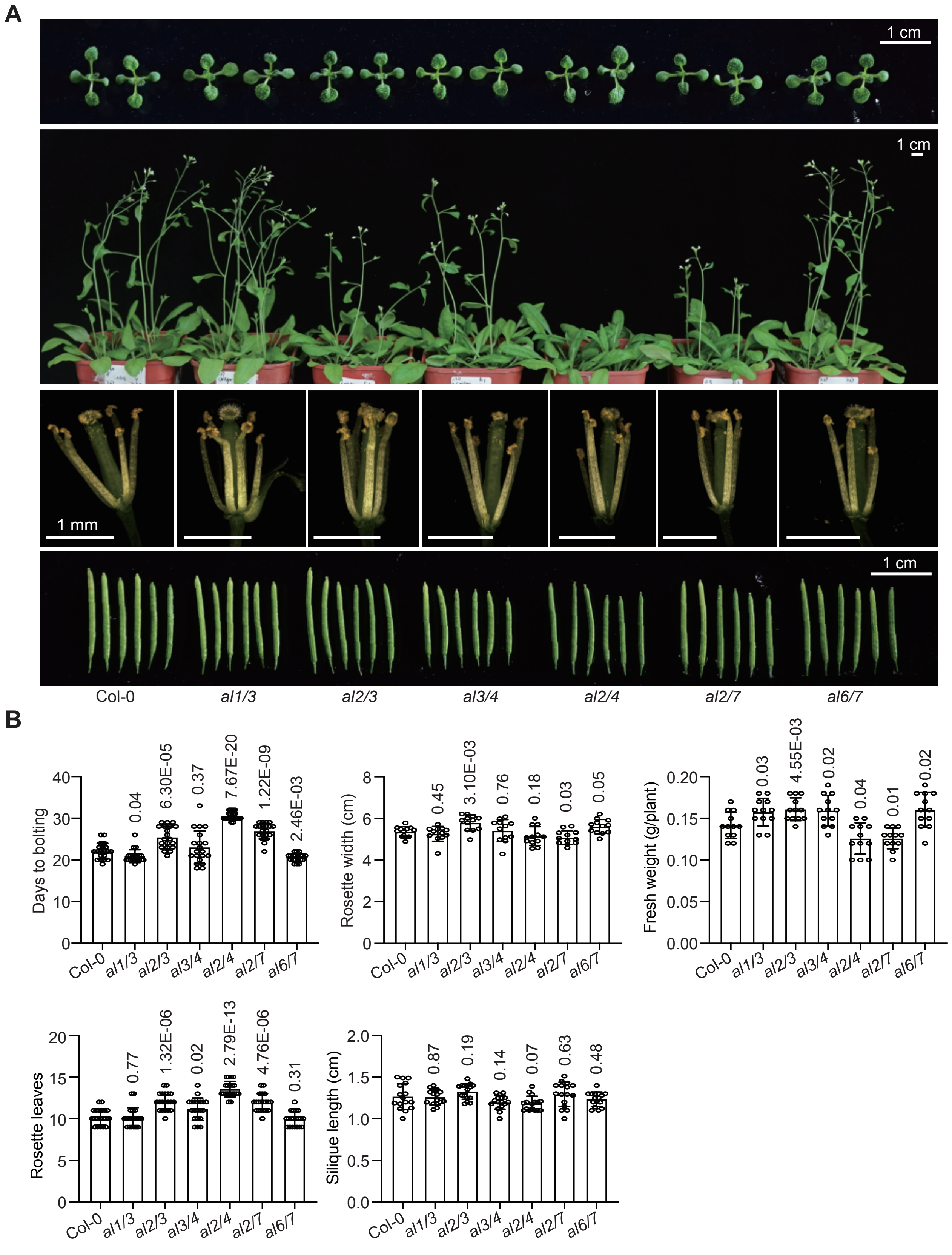


**Supplemental Figure 3. Morphological phenotypes of the *al* double mutants. (A)** From top to bottom, 11-day-old plants (top), 30-day-old plants, stamens and pistils under a microscope, and mature siliques (bottom) are shown for wild-type and *al* double mutants. **(B)** The statistical results of rosette width (20-day-old plants, *n* = 12), fresh weight (20-old-day plants, *n* = 12), days to bolting (*n* = 20), the number of rosette leaves (*n* = 20), and siliques length after maturing (*n* = 16). Values are mean ± SD. *P* values determined by two-tailed Student's t-test indicate the difference between the mutants and the wild-type control.

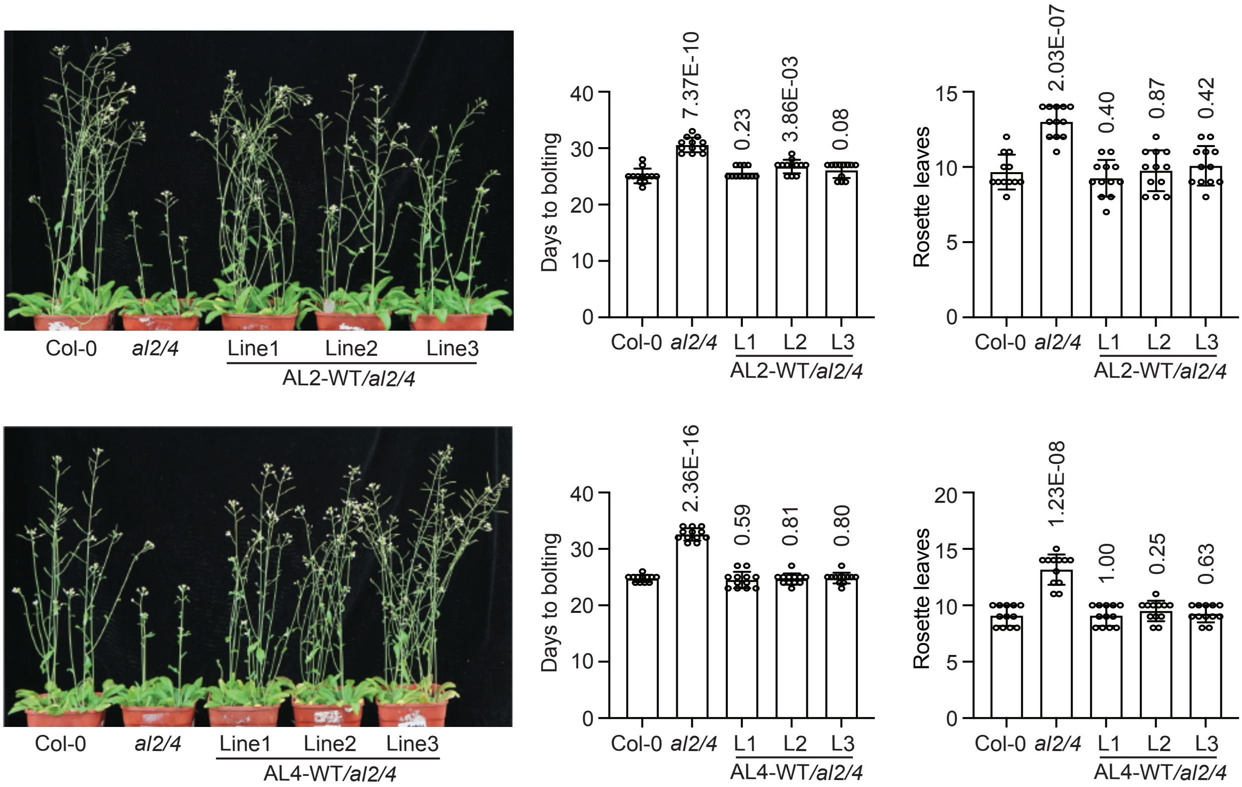


**Supplemental Figure 4. Phenotypic complementation test of the *al2/4* double mutant.** The flowering-time phenotype of the wild type, the *al2/4* mutant, and the *AL2* and *AL4* transgenic complementation lines. The morphological phenotype (left) and the statistical results of the number of days to bolting and the number of rosette leaves (right) are shown. The sample size is 12. Values are mean ± SD. *P* values were determined by two-tailed Student's t-test.

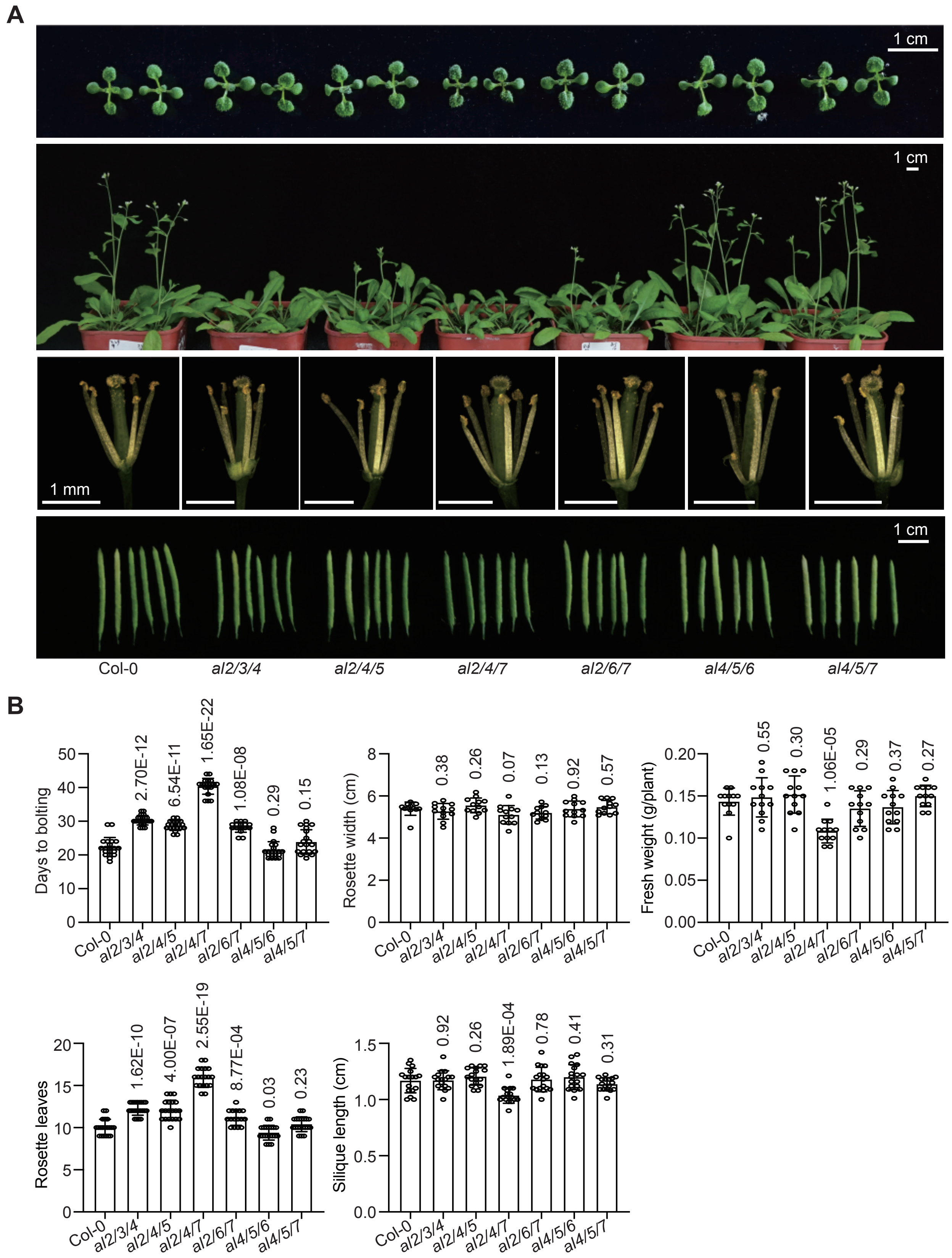


**Supplemental Figure 5. Morphological phenotypes of the *al* triple mutants. (A)** From top to bottom, 11-day-old plants (top), 30-day-old plants, stamens and pistils under a microscope, and mature siliques (bottom) are shown for wild-type and *al* triple mutants. **(B)** The statistical results of rosette width (20-day-old plants, *n* = 12), fresh weight (20-old-day plants, *n* = 12), days to bolting (*n* >= 16), the number of rosette leaves (*n* >= 16), and siliques length after maturing (*n* = 16). Values are mean ± SD. *P* values determined by two-tailed Student's t-test indicate the difference between the mutants and the wild-type control.

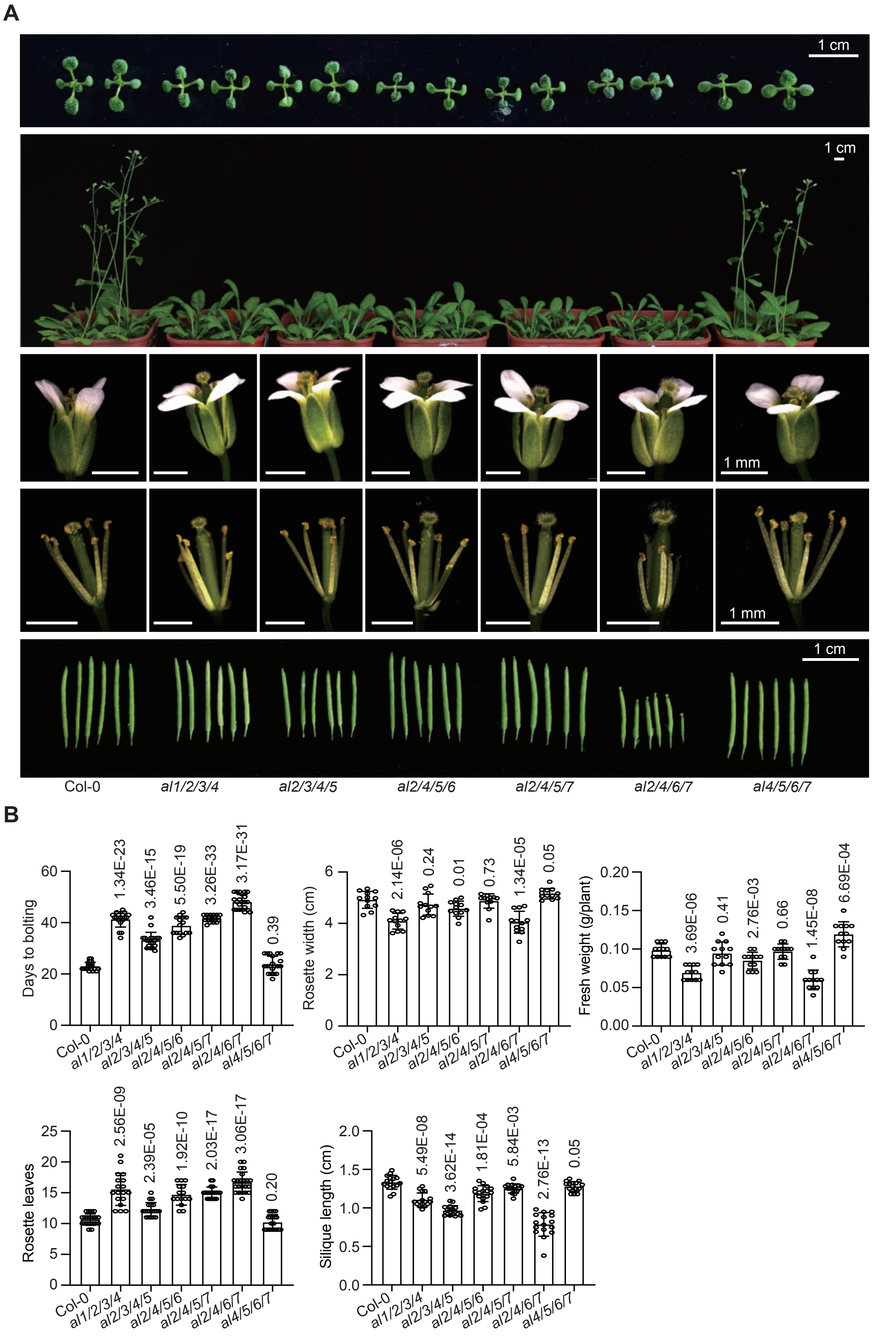


**Supplemental Figure 6. Morphological phenotypes of the *al* quadruple mutants. (A)** From top to bottom, 11-day-old plants (top), 31-day-old plants, flowers, stamens and pistils under a microscope, and mature siliques (bottom) are shown for wild-type and *al* quadruple mutants. **(B)** The statistical results of rosette width (20-day-old plants, *n* = 12), fresh weight (20-old-day plants, *n* = 12), days to bolting (*n* >= 15), the number of rosette leaves (*n* >= 15), and siliques length after maturing (*n* = 16). Values are mean ± SD. *P* values determined by two-tailed Student's t-test indicate the difference between the mutants and the wild-type control.

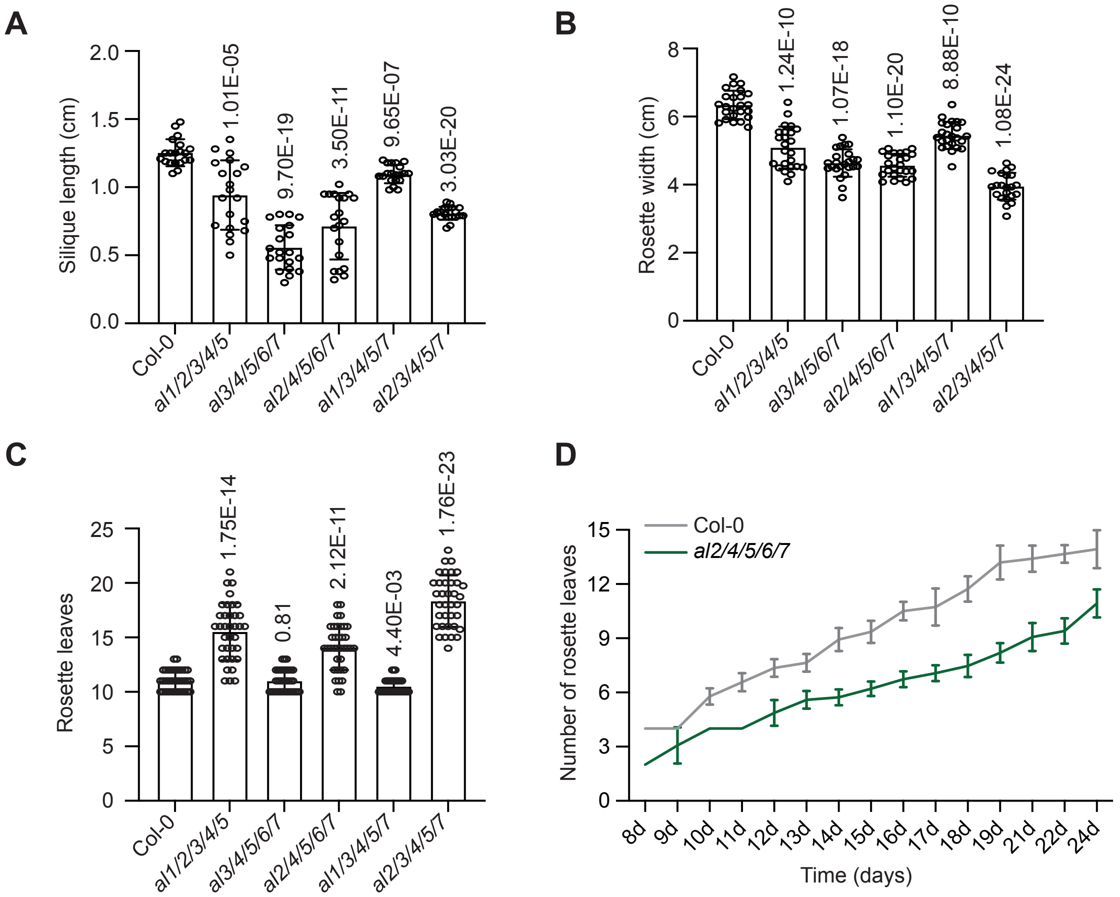


**Supplemental Figure 7. Statistical analysis of morphological phenotypes in the *al* quintuple mutants. (A-C)** The statistical analysis results of silique length, rosette width, and the number of rosette leaves between the *al* quintuple mutants and the wild type. The sample sizes for evaluating the length of mature siliques **(A)**, the rosette width **(B)**, and the number of rosette leaves **(C)** are 20, 24, and 36, respectively. Values are mean ± SD. *P* values determined by two-tailed Student's t-test indicate the difference between the mutants and the wild-type control. **(D)** The statistical analysis of the growth rate for rosette leaves in the *al2/4/5/6/7* mutant compared to the wild type.

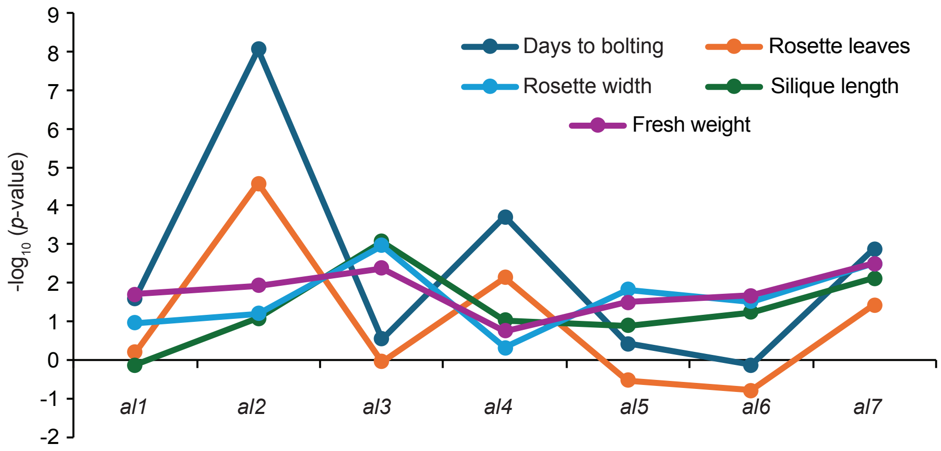


**Supplemental Figure 8. Statistical evaluation of the impact of individual *al* mutations on morphological change in *al* mutant plants.** The contribution coefficient for each *al* mutation was calculated based on the statistical significance of observed morphological changes between different *al* mutants, and was represented by $-\log_{10}$ (*p*-value). The morphological phenotypes include days to bolting, the number of rosette leaves, silique length, rosette width, and plant fresh weight. *P* values were determined based on all the *al* mutants tested in this study (Supplemental Figure 2-7).

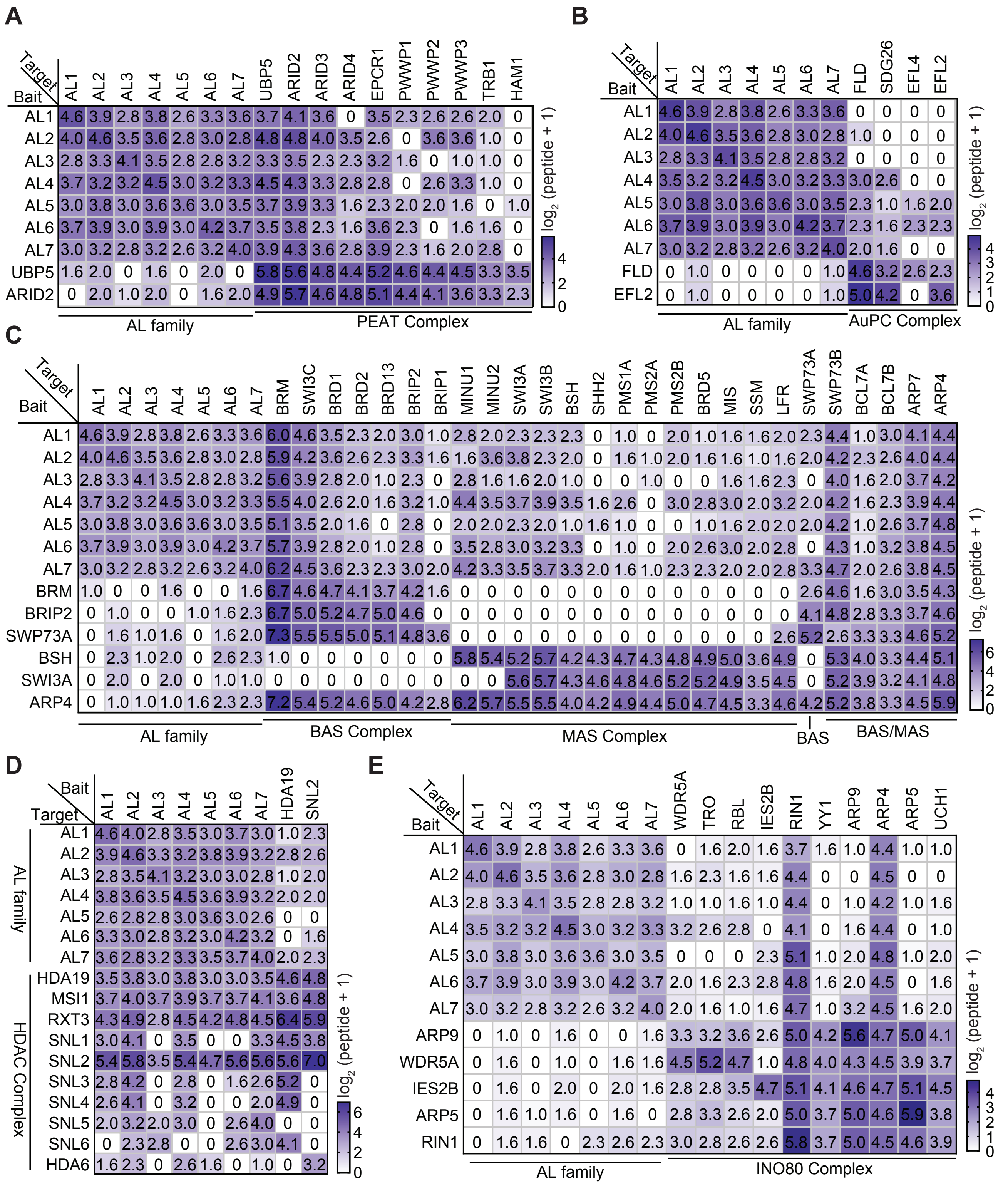


**Supplemental Figure 9. Determination of the interactions of AL proteins with PEAT, AuPC, SWI/SNF, HDAC, and INO80 complex components by AP-MS. (A-E)** Heatmap showing the co-purification of AL proteins with components of various chromatin-related complexes, including PEAT **(A)**, AuPC **(B)**, SWI/SNF **(C)**, HDAC **(D)**, and INO80 **(E)**, based on our previous AP-MS findings. The co-purification of these chromatin-related complexes with AL proteins, as determined by AP-MS in this study, is included in the heatmap. The color intensity represents the enrichment of normalized peptides detected by AP-MS.

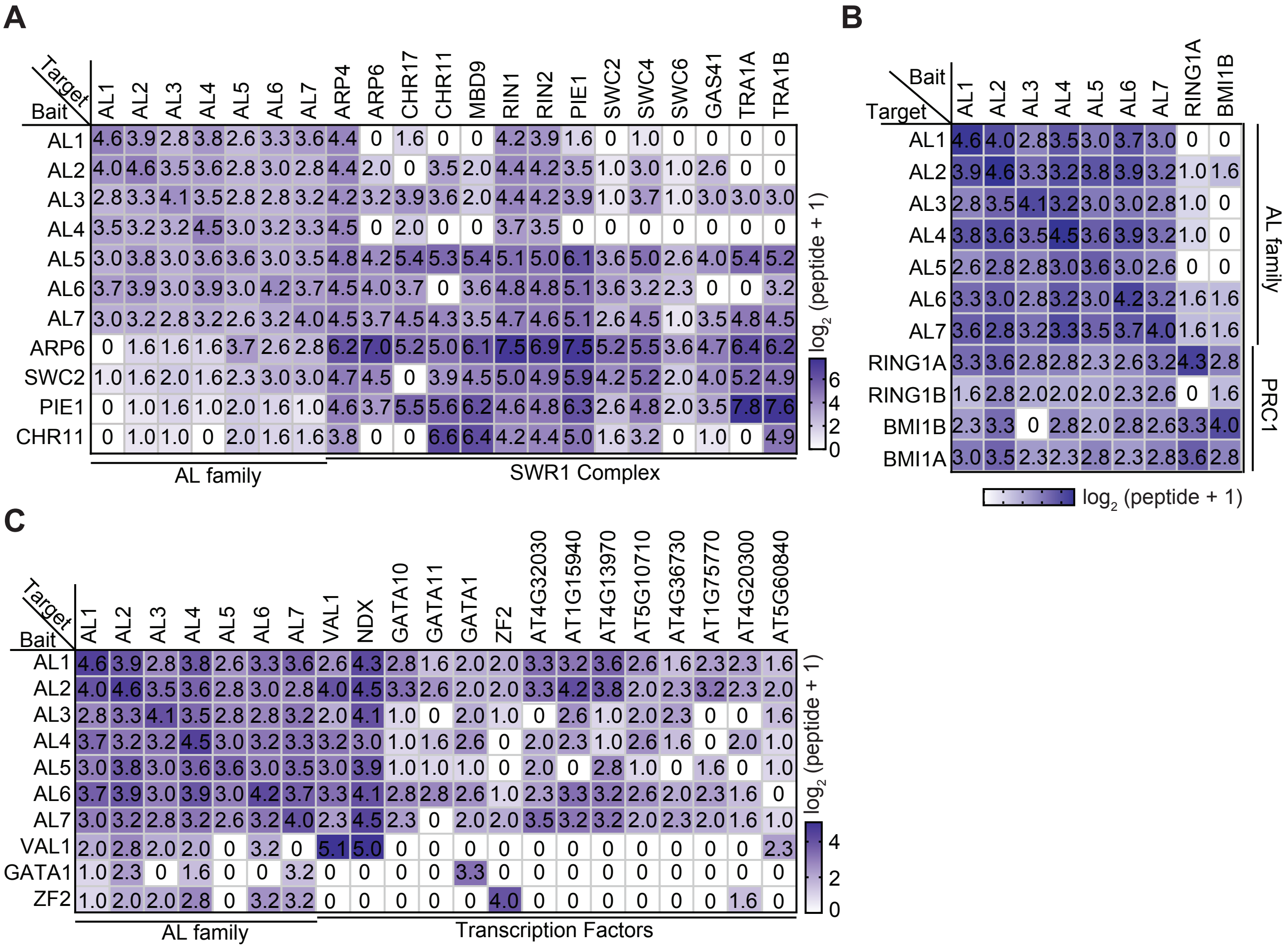


**Supplemental Figure 10. Determination of the interactions of AL proteins with SWR1 complex components, PRC1 complex components, and transcription factors by AP-MS. (A-C)** Heatmap showing the co-purification of AL proteins with SWR1 complex components **(A)**, PRC1 complex components **(B)**, and various transcription factors **(C)**. The color density represents the enrichment of normalized peptides identified by AP-MS. The co-purification of AL proteins with the SWR1 complex components, including PIE1, SWC2, CHR11, and ARP6, was detected by AP-MS in previous studies. The co-purification of AL proteins with PRC1 complex components and transcription factors was determined by AP-MS data generated in this study. Transgenic plants expressing indicated bait proteins were subjected to AP-MS.

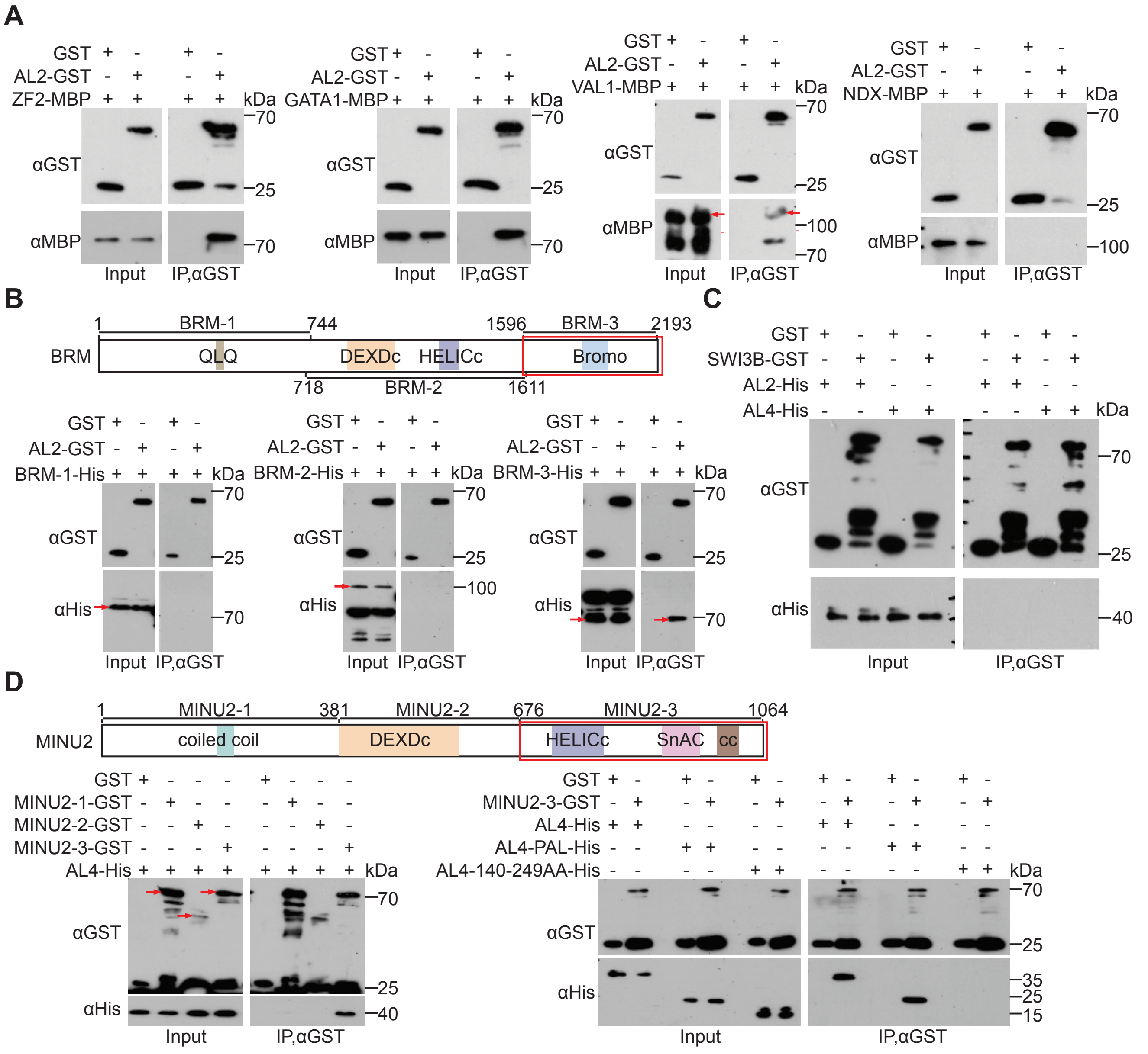


**Supplemental Figure 11. Determination of the interactions of AL proteins with transcription factors and SWI/SNF complex components by pull-down assays. (A)** The interactions of AL2 with the transcription factors ZF2, GATA1, VAL1, and NDX, as determined by pull-down assays. **(B)** Determination of the interaction between BRM and AL2 by pull-down assays. A schematic diagram indicates a series of truncated versions of BRM used in the pull-down assays, with the interaction domain marked with a red box. **(C)** Determination of the interactions between SWI3B and AL2 or AL4 by pull-down assays. **(D)** The interaction domains between MINU2 and AL4 are mapped using truncated protein versions. A schematic diagram shows truncated versions of MINU2 employed in pull-down assays, with the interaction domain marked with a red box.

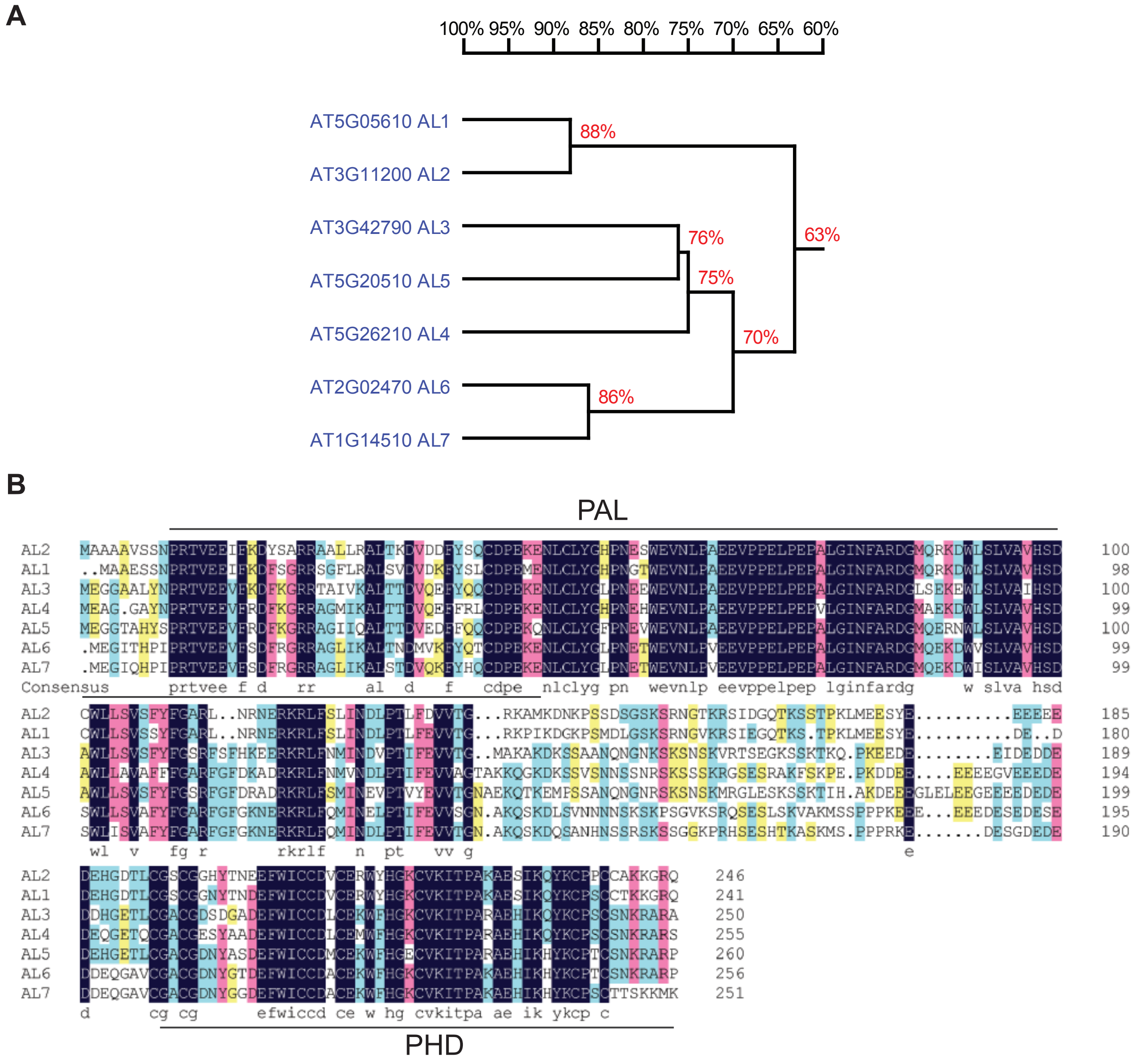


**Supplemental Figure 12. Sequence analysis of Arabidopsis AL proteins. (A)** Phylogenetic analysis of Arabidopsis AL proteins. A phylogenetic tree was drawn using the neighbor-joining method (with 1,000 bootstrap replicates) was drawn with the MEGA (version 7) software. **(B)** Sequence alignment of Arabidopsis AL proteins. In this alignment, identical amino acid residues are highlighted in black, while residues with varying degrees of conservation are denoted by pink, blue, and yellow shading. The positions of the PAL and PHD domains are indicated. The alignment was carried out using the DNAMAN (version 7) software.

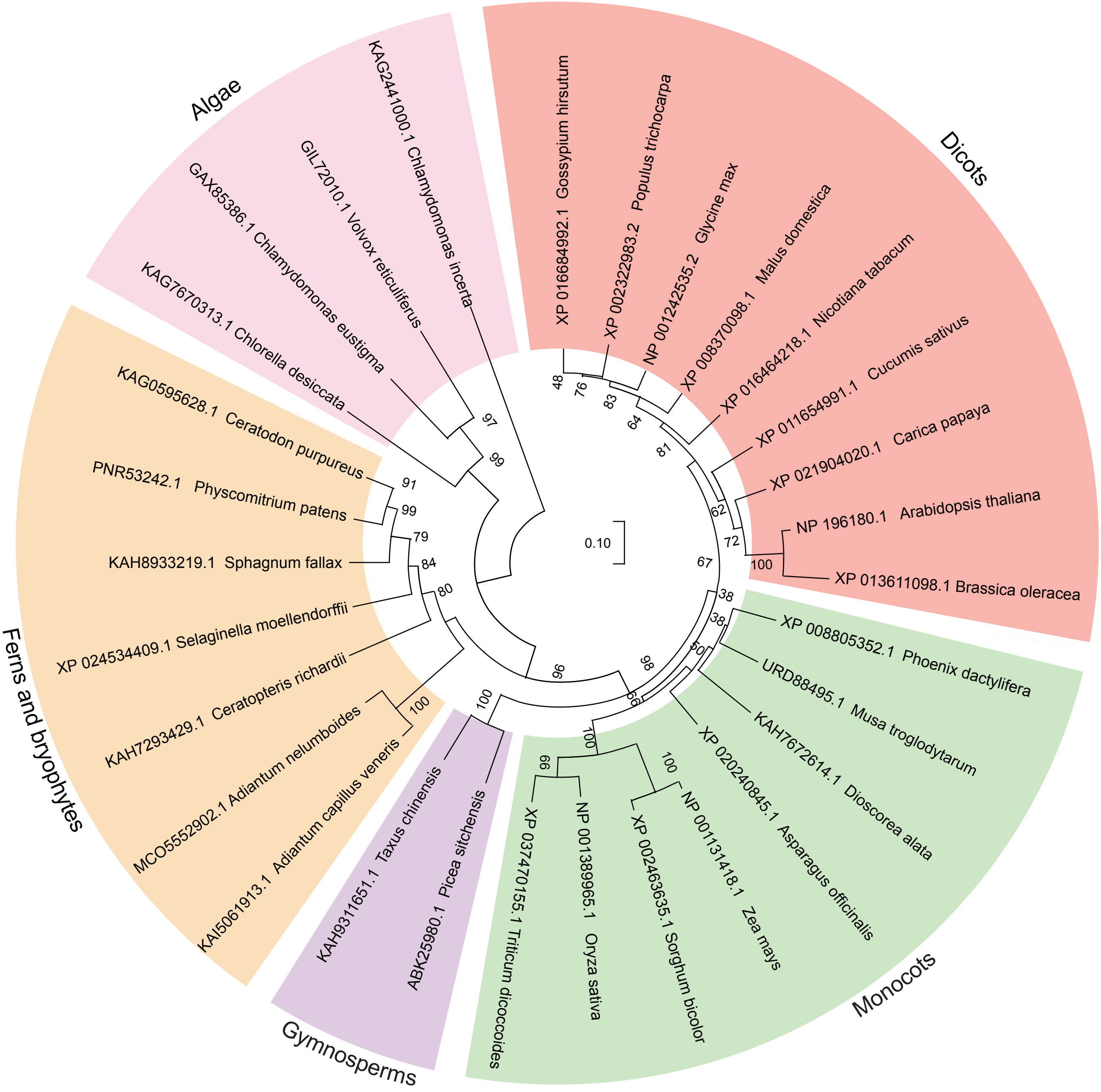


**Supplemental Figure 13. Phylogenetic analysis of AL1 orthologs in plants.** The Arabidopsis AL1 and the AL1 orthologs in algae, bryophytes, ferns, gymnosperms, monocots, and dicots were subjected to phylogenetic analysis. The orthologs of Arabidopsis AL1 in other plants were obtained by BLAST. Accession numbers are from the NCBI database. The phylogenetic tree was generated using the neighbor-joining method (with 1,000 bootstrap replicates) was drawn with the MEGA (version 7) software.

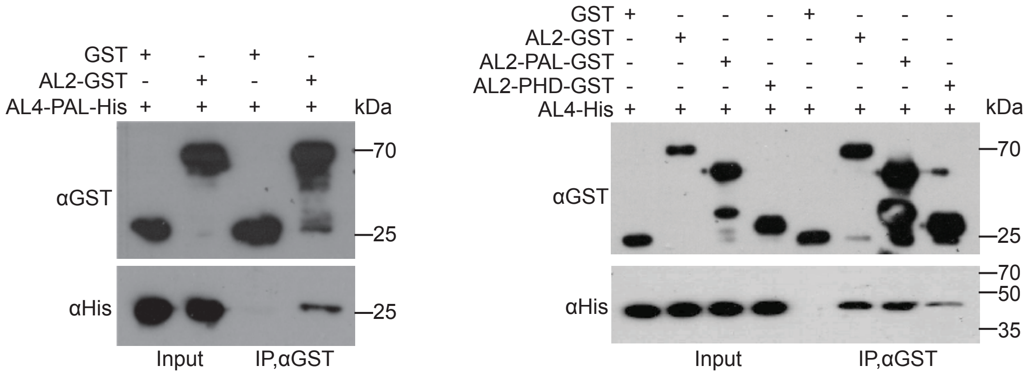


**Supplemental Figure 14. Determination of the interaction between AL2 and AL4 by pull-down assays.** The pull-down assays were conducted using full-length and truncated versions of AL2 and AL4. The truncated versions of AL2 and AL4 used in the pull-down assays include the PAL domain of AL4, the PAL domain of AL2, and the PHD domain of AL2.

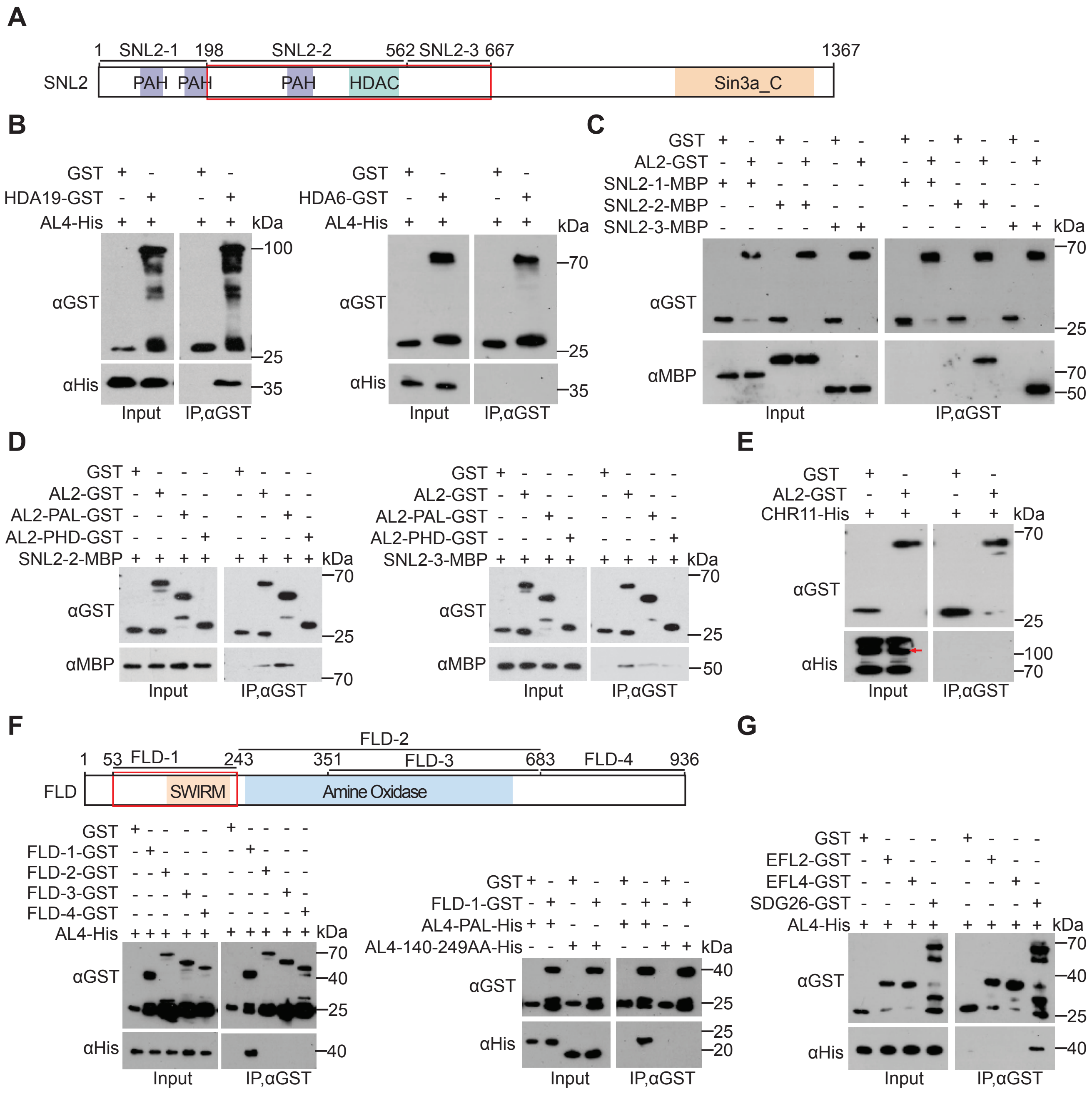


**Supplemental Figure 15. Determination of the interactions of AL proteins with HDAC complex components and AuPC complex components by pull-down assays. (A)** Schematic diagram of truncated versions of SNL2 used in pull-down assays. Conserved domains are shown in the diagram. The AL2 interaction domain within SNL2 is highlighted by a red box. **(B)** Determination of the interactions between AL4 and the HDAC catalytic subunits HDA19 and HDA6 by pull-down assays. The full-length AL4, HDA19, and HDA6 were used in the pull-down assay. **(C, D)** Determination of the interactions between AL2 and the HDAC accessory subunit SNL2 by pull-down assays. Truncated versions of SNL2 (SNL2-1, SNL2-2, and SNL2-3) were used in the pull-down assays. **(E)** Evaluation of the interaction between AL2 and CHR11 by pull-down assays. **(F, G)** Validation of the interactions between AL2 and AuPC complex components by pull-down assays. The AuPC complex components tested in the pull-down assays include FLD **(F)**, as well as EFL2, EFL4 and SDG26 **(G)**. A schematic diagram of truncated versions of FLD used in the pull-down assays is indicated, with the interaction domain of FLD marked with a red box.

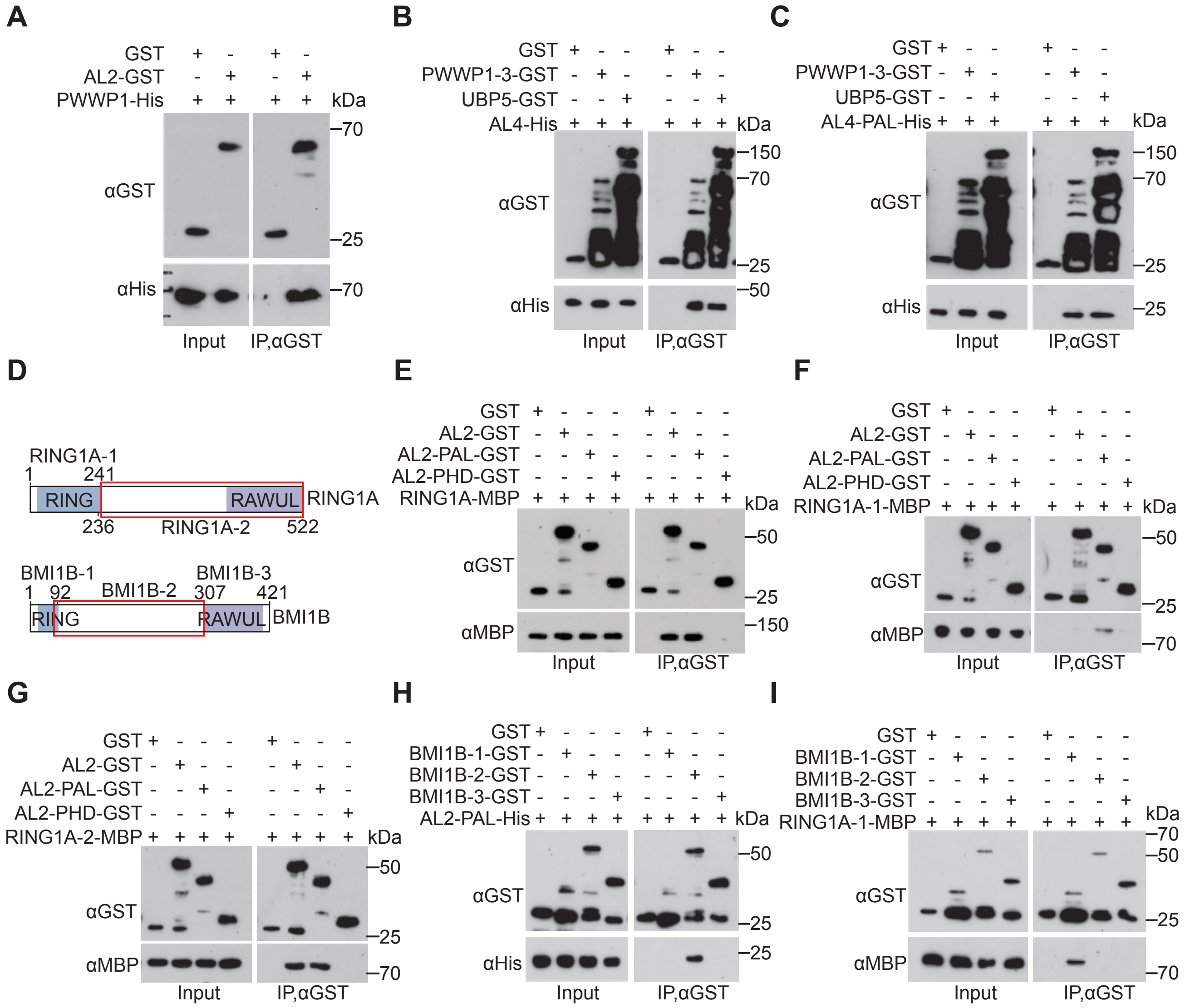


**Supplemental Figure 16. Determination of the interactions between AL proteins and components of PEAT or PRC1 complexes and between RING1A and BMI1B by pull-down assays. (A-C)** Pull-down assays showing the interactions between AL proteins and the PEAT complex components PWWP1 and UBP5. **(D)** Schematic diagrams of truncated versions of RING1A and BMI1B used in the pull-down assays. Conserved domains are shown in the diagrams. Red boxes represent the interaction domains within RING1A and BMI1B that are responsible for interaction with AL proteins. **(E-G)** Determination of the interaction between AL2 and RING1A by pull-down assays. The interaction domains of AL2 and RING1A were determined using both full-length and truncated versions of the proteins. **(H)** Determination of the interaction between AL2 and BMI1B by pull-down assays. The interaction between the PAL domain of AL2 and three truncated versions of BMI1B were detected. **(I)** Characterization of the interaction between the PRC1 complex components RING1A and BMI1B by pull-down assays. Truncated versions of RING1A and BMI1B were subjected to the analysis.

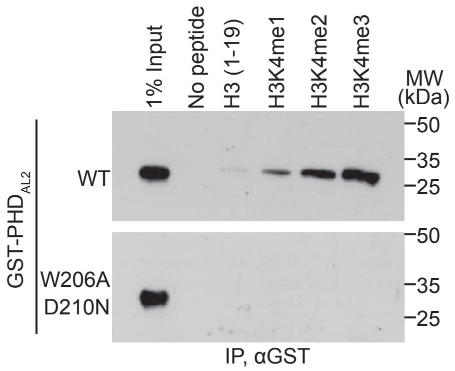


**Supplemental Figure 17. The W206A and D210N mutations within the PHD domain disrupt the binding of AL2 to methylated H3K4 peptides *in vitro*.** Histone peptide pull-down assays were conducted to assess the binding affinity of the wild-type PHD domain, as well as the mutated PHD domain harboring the W206A and D210N mutations, towards methylated H3K4 peptides. Both the wild-type and mutated PHD domains were expressed with a GST tag. Biotinylated histone peptides, methylated or unmethylated at H3K4, were used for the pull-down assays.

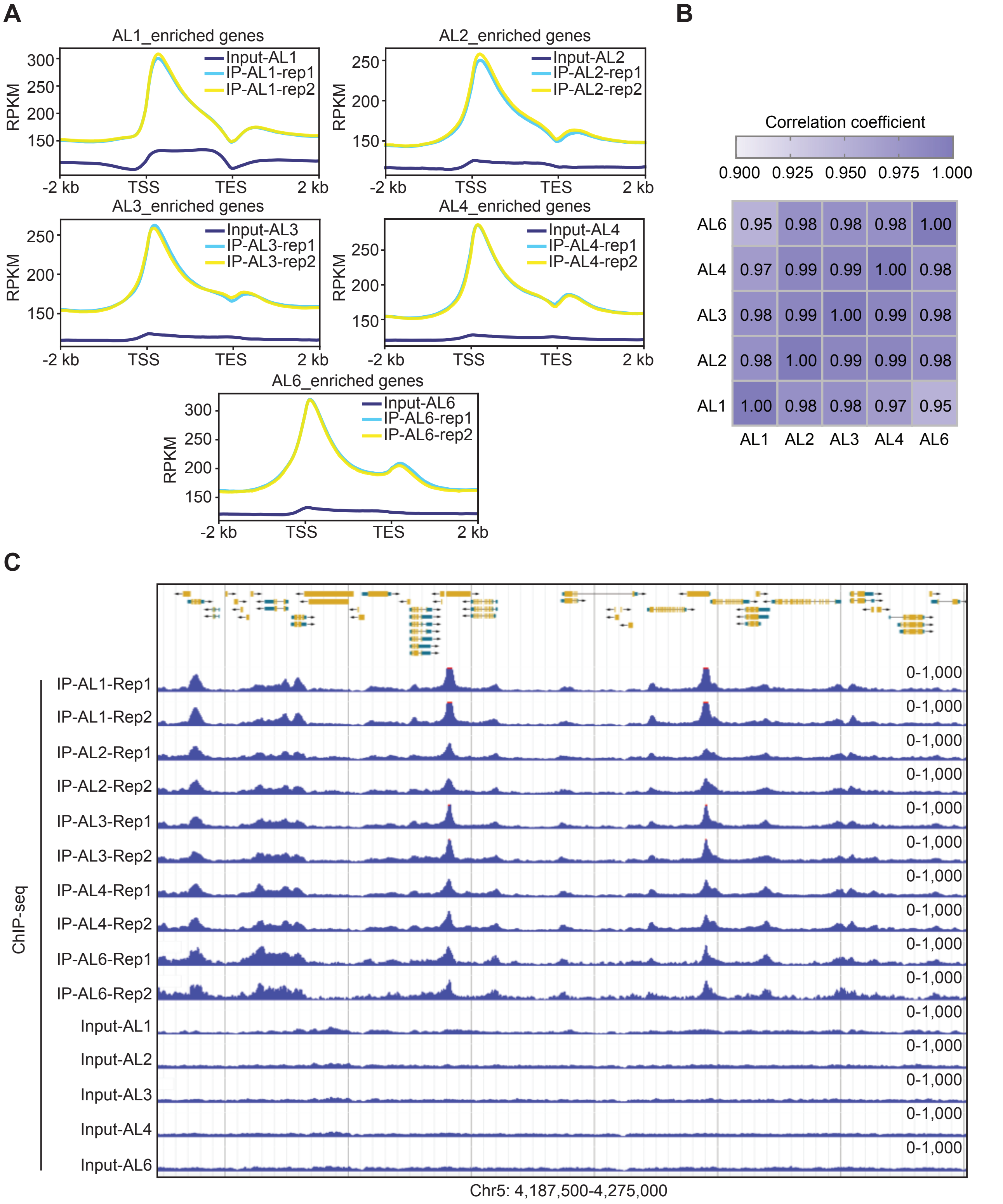


**Supplemental Figure 18. Co-occupancy of different AL proteins on chromatin at the whole-genome level. (A)** Meta plots depicting the distribution patterns of AL1, AL2, AL3, AL4, and AL6 across the genic region. Data from two independent biological replicates are shown. TSS, transcription start site; TES, transcription end site. **(B)** Pairwise correlation analysis of AL1, AL2, AL3, AL4, and AL6 ChIP-seq signals, with Pearson correlation coefficients displayed. **(C)** Genome browser visualization of the ChIP-seq signals for AL1, AL2, AL3, AL4, and AL6 at a representative genomic region. The scale of normalized reads is indicated.

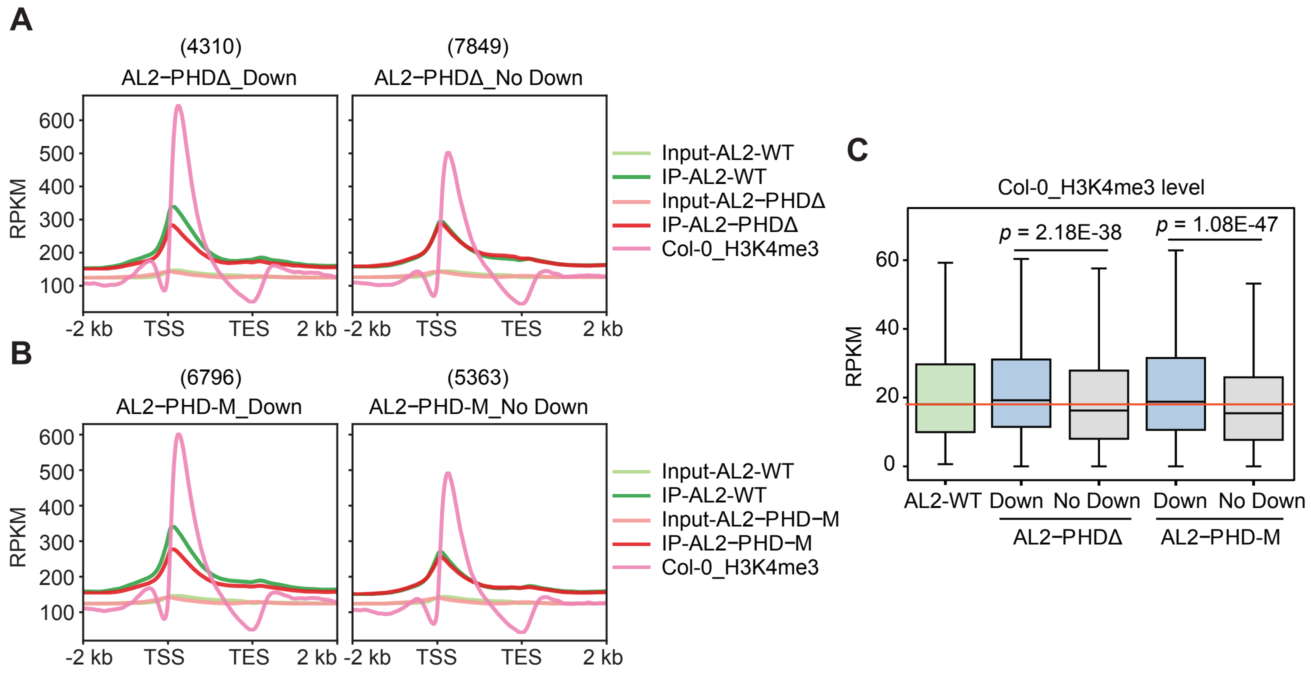


**Supplemental Figure 19. Determination of the impact of the deletion or mutation of the PHD domain on the association of AL2 with chromatin at the whole-genome level. (A, B)** Meta plots illustrating the enrichment levels of the wild-type AL2, the PHD-deleted AL2 (AL2-PHDΔ), the PHD-mutated AL2 (AL2-PHD-M), and H3K4me3 across AL2-enriched genes. Based on whether the PHD deletion **(A)** or mutation **(B)** reduce the association of AL2 with AL2-enriched genes, AL2-enriched genes were divided into two categories for the analysis. **(C)** Box plots showing the H3K4me3 levels at different categories of AL2-enriched genes. In the box plots, the center lines and box edges represent the medians and interquartile range (IQR), respectively. Whiskers extend to values within 1.5 times the IQR. *P* values were determined two-tailed Mann Whitney U test (no-paired).

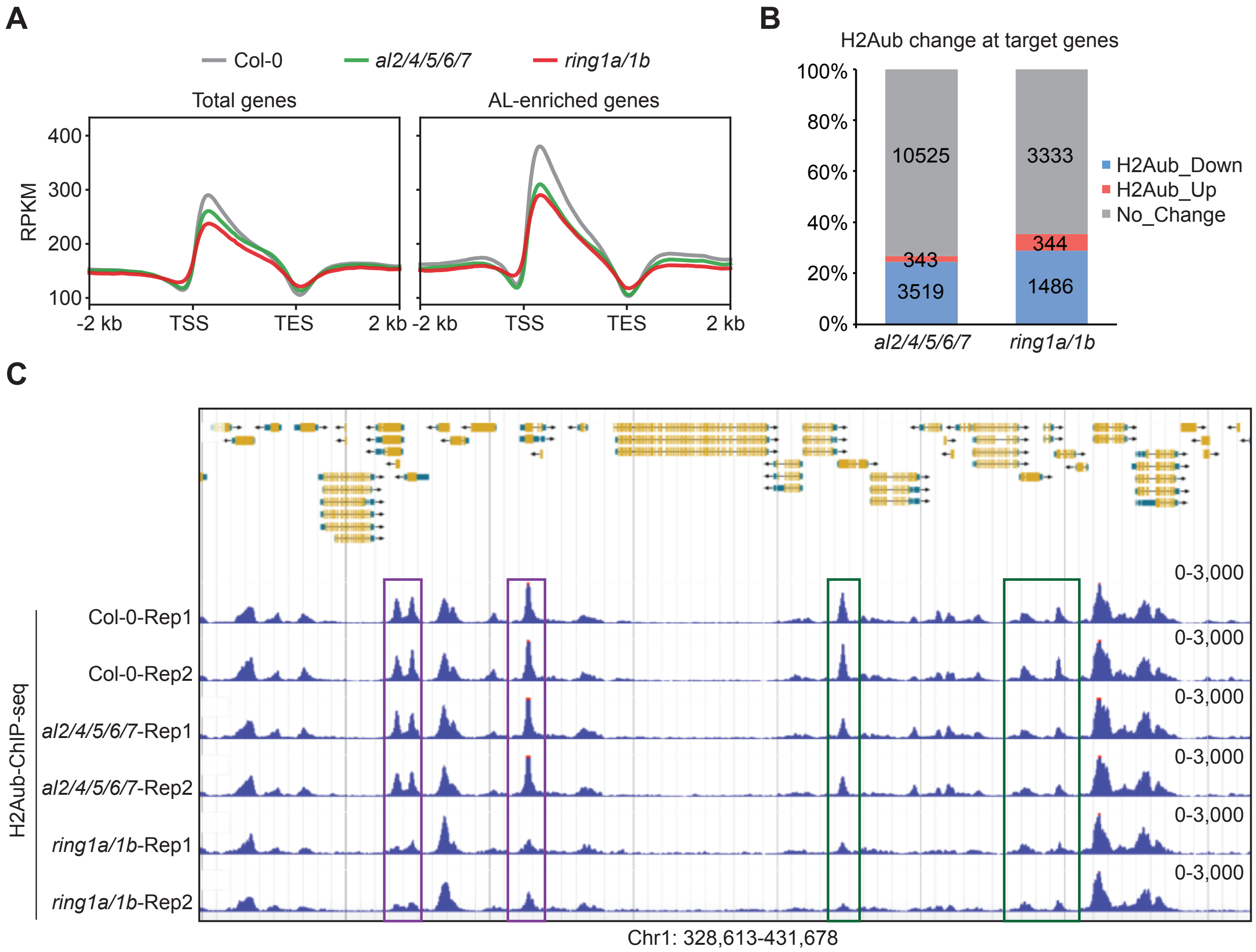


**Supplemental Figure 20. Comparison of the effect of *al2/4/5/6/7* and *ring1a/1b* on H2Aub. (A)** Meta plots showing the distribution patterns of H2Aub across total genes and AL-enriched genes in Col-0, *al2/4/5/6/7* and *ring1a/1b*. TSS, transcription start site; TES, transcription end site. **(B)** Analysis of the ratio of AL- and RING1A-enriched genes with up- and down-regulated H2Aub levels in *al2/4/5/6/7* and *ring1a/1b* mutants. The graph indicates the number of genes with up- (FC > 1.2, FDR < 0.05) and down-regulated (FC < 0.8, FDR < 0.05) H2Aub levels in the *al2/4/5/6/7* mutant relative to the wild type at AL-enriched genes, and in the *ring1a/1b* mutant relative to the wild type at RING1A-enriched genes. **(C)** Genome browser visualization of H2Aub ChIP-seq signals in wild type, *al2/4/5/6/7*, and *ring1a/1b* mutants within a representative genomic region. Purple boxes highlight genomic regions with AL-independent H2Aub, while green boxes denote regions with AL-dependent H2Aub. The scale of normalized reads is indicated.

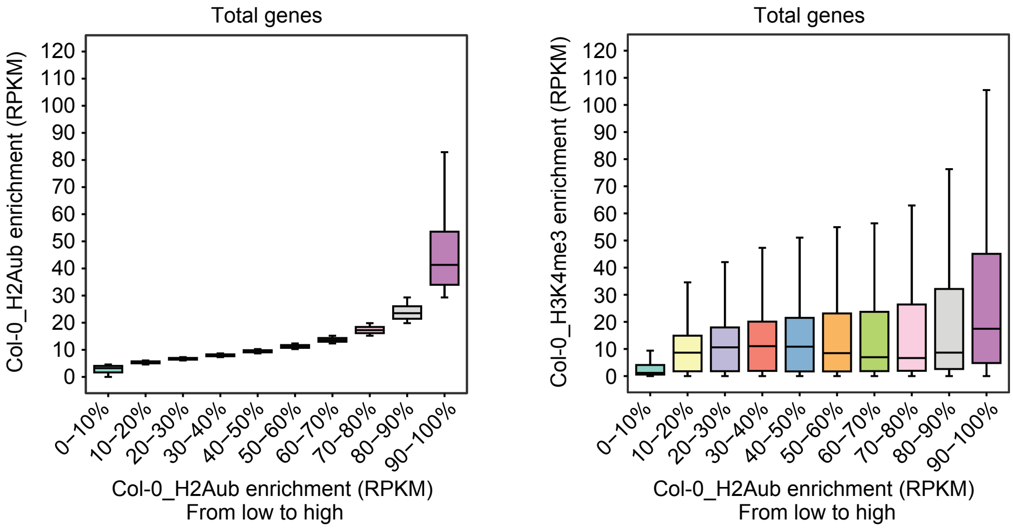


**Supplemental Figure 21. Analysis of the correlation between H2Aub and H3K4me3 at the whole-genome level.** The H2Aub levels of total Arabidopsis genes ($n$ = 32548) in the wild type were categorized into deciles in ascending order, as indicated in the left panel. The corresponding H3K4me3 levels for these deciles are shown in the right panel. In box plots, center lines and box edges are medians and the interquartile range (IQR), respectively. Whiskers extend to values within 1.5 times the IQR.

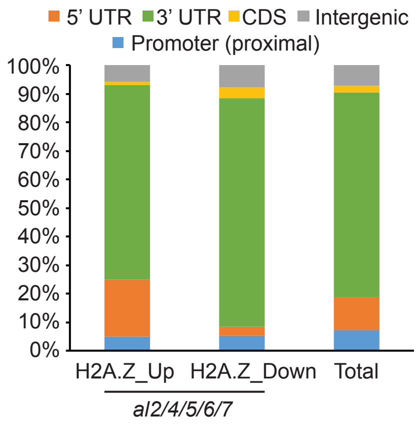


**Supplemental Figure 22. Analysis of the genomic distribution of H2A.Z level changes in the *al2/4/5/6/7* mutant relative to the wild type.** The genomic regions exhibiting up- and down-regulated levels of H2A.Z in the *al2/4/5/6/7* mutants compared to the wild type were annotated as promoter, 5'-UTR, CDS (coding sequence), 3'-UTR, and intergenic regions. The distribution of random genomic regions is shown as a control.
